# Pathwise approximations to solving the filtering problem for the stochastic chemostat

**DOI:** 10.1101/2023.01.17.524503

**Authors:** José Augusto Fontenele Magalhães, Muhammad Fuady Emzir, Francesco Corona

**Author notes:** 2000 MSC:* 60H15, 93E11, 65C05, 92D25, 37N25.

## Abstract

This paper concerns the inverse problem of characterising the state of a bioreactor from observations. In laboratory settings, the bioreactor is represented by a device called a chemostat. We consider a differential description of the evolution of the state of the chemostat under environmental fluctuations. We model the state evolution as a stochastic process driven by Brownian motion. Under this model, the state is described by its probability distribution in time, given the distribution of the initial state. The corresponding probability density function solves a deterministic partial differential equation (PDE), the Kolmogorov forward equation. This description is refined by incorporating observations. More formally, we seek the conditional distribution of the state given an observation process as the solution to a filtering problem, with the corresponding conditional probability density function solving a non-linear stochastic PDE, the Kushner–Stratonovich equation. In numerical studies of bioreactors, the well-posedness of these differential equations is frequently assumed rather than verified. Consequently, it remains unclear whether unique solutions exist and whether numerical approximations preserve key qualitative properties, such as positivity (e.g. non-negative chemical concentrations). This paper establishes the existence and uniqueness of solutions, ensuring well-posedness before presenting numerical approximations. Furthermore, we approximate the pathwise solution to the filtering problem by three approaches: a PDE-based approximation combining finite difference and splitting methods, a classical sequential Monte Carlo method, and a linearisation method. Our results show that the first two provide a richer probabilistic description of the state than the linearisation method, making them suitable for bioreactor control architectures.

## 1. Introduction

An inverse problem for a dynamic system refers to the construction of a set of equations given a differential description of the evolution of the system and a collection of observations related to quantities of interest. In this paper, the quantity of interest is the state variable of such a system denoted as *signal*, with inferences about the signal being obtained from indirect and noisy observations.

The study of this type of inverse problem has been the focus of considerable theoretical and numerical work under the name *stochastic filtering*. The aim is to obtain expectations with respect to the probability distribution of the signal at the current time, this distribution being conditional on noisy observations accumulated up to that time. The theoretical features of this problem are well-known: such expectations require integration with respect to a stochastic process modelling the observations, and the probability density function of the conditional probability distribution of the signal solves a non-linear stochastic partial differential equation, the Kushner–Stratonovich equation (KSE). A comprehensible account of this problem is available in [1, 2].

Filtering theorists are typically concerned with analysing the differential equations governing the filtering problem. Among other topics, they study the existence, uniqueness, and growth behaviour of solutions to the stochastic differential equations that describe the evolution of the underlying process and the observations. This is crucial for numerical approximations of the expectations of interest as most analyses of existing algorithms rely on such properties [1, 3].

Filtering practitioners have adopted the pathwise formulation of the filtering problem. This is because when observing most real-world signals, the single “path” for the observations never exactly aligns with the observation model: signals are sampled at a high frequency, with sampled points artificially connected, for instance, by straight-line segments. While this formulation enables various lines of research [4, 5], this work focuses on fixing the observation path to reduce the aforementioned KSE to a partial differential equation (PDE) with a random coefficient.

The conditional probability density function in pathwise filtering can be derived in closed-form for certain problems, for instance, those with a linearly evolving observation process (the observation is a noisy, scaled version of the signal) [6, 7]. However, even in this scenario, the signal is only allowed to have a non-linear evolution under some restrictive conditions (see Beneš condition [6]), rarely met in real-world systems.

With limited closed-form solutions, much research has been directed towards constructing algorithms to approximate the solution in more general settings. The approximations can be obtained via, but not limited to, three approaches: the extended Kalman filter (EKF), particle filters (PFs), and methods for solving PDEs.

The EKF is the classical approximation method. It linearises certain non-linearities in the model to ensure that the traditional Kalman–Bucy filter [7] can be applied. PFs, also known as sequential Monte Carlo methods, have been widely developed since their conception in the 1990s [8, 9] and are among the most popular alternatives to the EKF. In PFs, particles (also called samples) first evolve according to a desirable transition kernel and then selectively interact in accordance with their likelihood with respect to the observation path. The EKF, PFs and their derivatives have practical applications across diverse domains, including target tracking in robotics [10], weather forecasting [11], and disease outbreak monitoring in epidemiology [12].

A less common approach consists of approximations via a splitting method: the PDE with a random coefficient is decomposed into two parts: a deterministic and a normalisation one. The first part is independent of the observations and may therefore be solved offline. The second part consists of computing a correction and normalisation factor from the observation path, ensuring that the resulting density integrates to unity. Several frameworks have been proposed to perform this decomposition and efficiently transfer the offline solution to the online stage, ranging from the early splitting methods of Le Gland [13, 14] to the real-time framework developed by Yau and Yau [15, 16, 17]. The offline PDE can be solved in low-dimensional state spaces using conventional numerical methods such as finite-difference [18] or spectral discretisations [19]. More recently, several deep learning-based approaches have also been proposed to approximate the solution of the offline PDE, providing a mesh-free alternative to classical discretisation techniques [20, 21]. Regardless of the numerical approximation employed for the offline problem, grid-based methods remain an effective approach for low-dimensional filtering problems [22, 23], as they provide an explicit approximation of the filtering density over the entire state space, enabling visualisation and further analysis without restrictive assumptions on its form.

This study considers the practical implementation of filtering with a focus on a classical splitting method. Our goal is to obtain the probabilities and expectations for a process performed in a bioreactor, in which one or more populations of microorganisms grow in a nutrient medium consisting of a cocktail of molecules. In particular, we consider a laboratory device representing this bioreactor: a chemostat. Prior to the genomic era, the chemostat was a major player in the use of continuous culture systems to characterise certain kinds of microbes [24, 25]. Being a thoroughly tested continuous culture system, it has regained some interest in this post-genomic era. For instance, chemostat-grown cultures lead to reproducible and homogeneous results which are necessary to identify the roles played by genes in the biology of organisms [26].

We consider a chemostat under environmental fluctuations, specifically a chemostat subjected to Brownian motion processes with multiplicative noise, herein called the stochastic chemostat. We adopt the dynamic model proposed in [27], as its differential equations have been shown, by means of martingale theory, to admit a unique solution within a suitably chosen space. In particular, [27] ensures that their model would allow for its realisations remaining within a biologically meaningful domain (e.g. positive chemical concentration values). Finally, to describe the relationship between states and observations, the dynamics are coupled with an observation model in which the observations are also subjected to Brownian motion.

The existing state-of-the-art of filtering for the stochastic chemostat comprises an approximation to the first two moments of the conditional probability distribution of the signal by EKF and a more complex approximation by PFs [29]. However, to the best of our knowledge, no previous work has addressed the theoretical guarantees of existence, uniqueness, and growth behaviour of the solutions to our differential equations of interest, nor has the pathwise solution to the filtering problem been numerically realised by grid-based schemes.

The following are the contributions of this paper:

- First, Theorem 3.1 shows that the differential equations for the stochastic chemostat have a unique solution within a suitably chosen space. We provide an alternative proof to the results in [27] via Lyapunov functions, aligning with existing stochastic population dynamics literature (e.g. [30] on the Lotka-Volterra model). Further, our proof introduces tools that will be useful later in the analysis, particularly in establishing the well-posedness of the PDEs.
- Secondly, Proposition 3.1 addresses the well-posedness of the Kolmogorov forward equation for the stochastic chemostat. We also discuss the implication of this result on the pathwise form of the Kushner–Stratonovich equation. Valid conclusions from numerical simulations of these PDEs require guarantees of existence and uniqueness of their solution. To the best of our knowledge, previous studies have assumed well-posedness without formal derivation. In contrast, our work explicitly investigates if these assumptions hold.
- Lastly, we demonstrate the pathwise approximations using the splitting method, comparing them against EKF and PF results. This allows us to move beyond merely visualising expectations. Instead, the shape of the probability density across the state space is available, providing a more comprehensive understanding of the system dynamics and the associated uncertainty.

This paper is structured as follows. Section 2 provides the theoretical foundations for understanding the filtering problem. In particular, Section 2.1 introduces assumptions ensuring the existence, uniqueness, and growth properties of solutions to key differential equations. Section 2.2 introduces the filtering problem and its pathwise formulation, covering both the theory and approximation methods. Section 3 applies the concepts from Section 2 to the stochastic chemostat. In particular, Section 3.1 discusses the properties of solutions to the differential equations describing the evolution of the state of the stochastic chemostat. Finally, Section 3.2 presents the pathwise filtering problem for the stochastic chemostat, showing its numerical solution based on the aforementioned approximation methods. Section 4 concludes the paper.

## 2. Preliminaries: Stochastic filtering

Filtering refers to a methodology in which observations and a model of a dynamic system are combined to make inferences about its state. The state variable of this dynamic system is denoted as the signal. In stochastic filtering, the signal is modelled by a continuous-time stochastic process *X* = (*X*_*t*_, *t* ≥ 0), with *t* denoting time. While the signal cannot be directly observed, an indirect observation can be obtained. This observation is assumed to be described by a continuous-time stochastic process *Y* = (*Y*_*t*_, *t* ≥ 0) denoted as the observation process. Furthermore, *Y* is assumed to be a function of *X* corrupted by an observation noise, modelled by a stochastic process 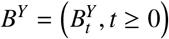. For inferences on *X* from *Y*, we let 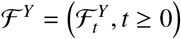 be the filtration generated by *Y*, where the *σ*-field 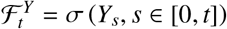 is all the information from observations up to time *t*. The resulting filtering problem consists of determining the conditional distribution *π*_*t*_ of *X*_*t*_ given 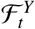, In a mean-square sense, the estimate 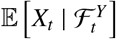 of the value of the signal at time *t*, the conditional mean of *X*_*t*_ given 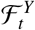, is optimal. Other inferences about *X* from *Y* are quantities of the form 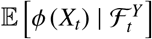 for some function *ϕ* over the state space of the signal. Notably, the conditional distribution of the signal is also modelled by a stochastic process *π* = (*π*_*t*_, *t ≥* 0).

This section provides a compact overview of this problem and the background needed for our application. Section 2.1 reviews a number of notions and results about stochastic differential equations, such as existence, uniqueness, and Markovianity of solutions. Section 2.2 concerns the filtering problem. Since the evolution of the system is described in continuous-time, we present filtering equations using a body of techniques from stochastic calculus outlined in Section 2.1. In particular, Section 2.2.1 presents these equations when we assume that (1) the signal process *X* takes values in an *N*_*X*_-dimensional state space S_*X*_ and that *X* is the solution to a stochastic differential equation driven by an 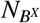-dimensional Brownian motion process 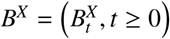, and (2) the observation process *Y* takes values in 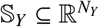 and that *Y* has an evolution which depends on both *X* and an 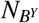-dimensional Brownian motion process 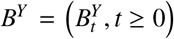. Section 2.2.2 particularises the filtering equations for a case in which the pair (*X, Y*) is Markovian. The main computational methods for solving these equations are also outlined.

### 2.1. Stochastic differential equations (SDEs)

To set the notation, we consider an identity modelling the evolution of some deterministic process *z* = (*z*_*t*_, *t* ≥ 0),

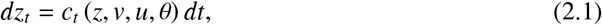

where *z*_*t*_ ∈ S_*z*_ is the value at time *t* of the process of interest, *dz*_*t*_ is called the differential of the process *z* at time *t, dt* is an infinitesimal time increment, *v* = (*v*_*t*_(*z*), *t* ≥ 0) with *v*_*t*_ ∈ S_*v*_ is an auxiliary process possibly dependent on the entire process *z, u* = (*u*_*t*_ *(z*_*s,s*≤*t*_ *), t* ≥ 0) with *u*_*t*_ ∈ S_*u*_ is a process which describes the evolution of state-feedback control actions and is possibly dependent on past values of *z*, and *θ* = (*θ*_*t*_ = const, *t* ≥ 0) with *θ*_*t*_ ∈ S_*θ*_ is a process which characterises the model parameters that are constant in time. The coefficient *c*_*t*_ is the functional which specifies the dynamics. At this point, it suffices to assume that S_*z*_, S_*v*_, S_*u*_ and S_*θ*_ are complete separable metric spaces. Without loss of generality, processes *u* and *θ* are omitted hereafter. Informally, note that popular forms of ordinary differential equations (ODEs) can be recovered from identity (2.1). For instance, if processes *z* and *v* are smooth enough, then *c*_*t*_ is the derivative of *z* with respect to time evaluated at time *t*. Furthermore, if *v* is specified up to time *t* and takes the form *v* = (*v*_*t*_ *(z*_*s,s*≤*t*_), then *c*_*t*_ (*z, v*) = *c*_*t*_ *(z, v*_*t*_ *(z*_*s,s*≤*t*_ )). For the particular case of first-order differential equations

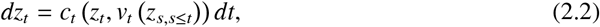

knowing the past (*v*_*s*_, *s* ≤ *t*) of *v* and any value *z*_*s*_, *s* ≤ *t*, allows to determine the conditions under which a process *z* satisfying ODE (2.2) exists and is unique (Picard’s theorem in, for instance, [31]): typically, the initial (at *s* = 0) condition *z*_0_ is provided.

Stochastic differential equations can be understood as deterministic differential equations into which a degree of randomness is introduced. When the process *v* is a stochastic process *V* = (*V*_*t*_, *t* ≥ 0) and/or the initial condition *z*_0_ is a random variable, the process *z* which satisfies identity (2.1) becomes a stochastic process *Z* = (*Z*_*t*_, *t* ≥ 0). An additional source of randomness can be introduced by including another stochastic process *B* = (*B*_*t*_, *t* ≥ 0). This procedure results in the class of differential equations known as stochastic differential equations (SDEs), whose solutions involve ordinary and stochastic integrals. We consider stochastic integration with respect to *B*, particularly for the case in which *B* is a standard Brownian motion process. We provide a full characterisation of such a process in the Supplementary Material. Lastly, we let (Ω, ℱ, ℙ) be the appropriate probability space for all processes considered in this paper.

To provide an instrumental account of the theory of SDEs, we start by letting (1) *C*_*Z*_ = *C*_*Z*_ ([0, ∞), S_*Z*_) be the space of continuous functions defined on the time index set [0, ∞) and taking values in 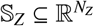; and (2) *D*_*V*_ = *D*_*V*_ ([0, ∞), S_*V*_) be the space of right-continuous functions also on [0, ∞), with left-hand limits, and taking values in a complete separable metric space S_*V*_ such that regular conditional probabilities exist. We use ℬ (*C*_*Z*_) and ℬ (*D*_*V*_) to denote the *σ*-fields of Borel sets of *C*_*Z*_ and *D*_*V*_.

We introduce into identity (2.1) a random initial condition *z*_0_ = *z*_0_ (*ω*)_*ω*∈Ω_, an S_*V*_-valued stochastic process *V* = (*V*_*t*_, *t* ≥ 0) with sample paths *V* (*ω*) ∈ *D*_*V*_ for ℙ-almost every *ω* ∈ Ω, and a process written in terms of an 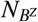-dimensional standard Brownian motion process 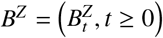, where 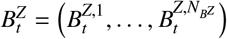. The resulting SDE which models the evolution of *Z* is

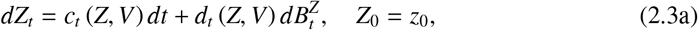

or, equivalently, in coordinate form, for each *n*_*Z*_ = 1, …, *N*_*Z*_,

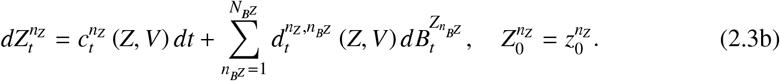

In SDE (2.3), the functionals 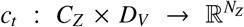 and 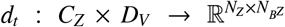 are ℬ (*C*_*Z*_) × ℬ (*D*_*V*_)-measurable coefficients; that is, we have that 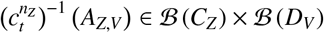 and 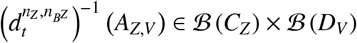, for every *A* ∈ ℬ (ℝ).

In the following, we limit ourselves to the class of non-anticipative functionals *c*_*t*_ and *d*_*t*_; that is, the values of *c*_*t*_ and *d*_*t*_ only depend on the paths *V* (*ω*) and *Z* (*ω*) up to time *t* (we provide a detailed mathematical definition in the Supplementary Material).

#### 2.1.1. Existence and uniqueness of the solution

The notion of a solution to SDE (2.3), as well as the conditions for its existence and uniqueness, can be stated whenever certain conditions on *c*_*t*_ and *d*_*t*_ are satisfied. These conditions are identified after noting that SDE (2.3) is a shorthand to the following stochastic integral equation:

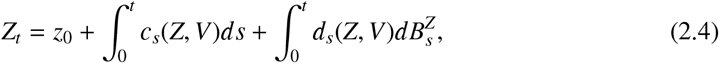

where the first integral is a Riemann integral and the second one is an Itô integral. Itô integrals are defined for integrands which allow such integrals to be martingales. For details on the martingale theory for Itô integrals, see [32].

Based on Eq. (2.4), the notion of a solution of SDE (2.3) can be constructed after establishing the conditions needed on the collection of integrands (*c*_*t*_(*Z, V*), *t* ≥ 0) and (*d*_*t*_(*Z, V*), *t* ≥ 0) to allow the two integrals to be well-defined. To this end, we first endow our probability space with a notion of time: this is done by specifying which sub-*σ*-field of information in ℱ can be gathered by time *t*. Secondly, we define an increasing family (ℱ_*t*_, *t* ≥ 0) of *σ*-fields of ℱ with ℱ_*s*_ ⊂ ℱ_*t*_, for *s, t* ∈ [0, ∞), with *s* ≤ *t*, and such that ℱ_0_ includes all ℙ-null sets in ℱ. Then, for the resulting filtered probability space (Ω, ℱ, (ℱ_*t*_, *t* ≥ 0), ℙ) over which *B*^*Z*^, *V* and *z*_0_ are defined, we have:

##### Definition 2.1

**(Strong solution [33])**. *Let c*_*t*_ *and d*_*t*_ *be non-anticipative functionals. Let* (Ω, ℱ, (ℱ_*t*_, *t* ≥ 0), ℙ) *be a filtered probability space for B*^*Z*^, *V, and z*_0_. *A stochastic process Z is said to be a strong solution of SDE* (2.3) *if the following holds:*

1. *Z is adapted to* (ℱ_*t*_, *t* ≥ 0); *that is, each Z*_*t*_ *is measurable with respect to* ℱ_*t*_;
2. ℙ*-almost surely* 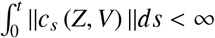 *and* 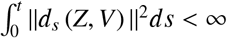, *for all t* ∈ [0, ∞) *and for an Euclidean norm of c*_*s*_ *and a Hilbert-Schmidt norm of d*_*s*_;
3. ℙ*-almost surely Z solves Eq*. (2.4).

The non-anticipativeness of *c*_*t*_ and *d*_*t*_, together with (1), implies that *c*_*t*_ (*Z, V*) and *d*_*t*_ (*Z, V*) are measurable and adapted to ℱ_*t*_+ for all *t* ≥ 0, with ℱ_*t*_+ = ∩_*s*>*t*_ ℱ_*s*_ (see, [32, Section 4.2]). Imposing (2) on the measurable ℱ_*t*_+-adapted process (*d*_*t*_ (*Z, V*), *t* ≥ 0) is sufficient to render the Itô integral in Eq. (2.4) a well-defined quantity. Because of (1), notably, a strong solution *Z* must be adapted to (ℱ_*t*_, *t* ≥ 0): for instance, if ℱ_*t*_ is the *σ*-field generated by *B*^*Z*^, *V* and *z*_0_, then the values of *Z*_*t*_ are determined by *z*_0_ and by th e values of *B*^*Z*^ and *V* up to *t*. Adaptedness to (ℱ_*t*_, *t* ≥ 0) can be too strict, and *weak solutions (*Ω, ℱ, (ℱ_*t*_, *t* ≥ 0), ℙ, *B*^*Z*^, *V, z*_0_, *Z)* which satisfy SDE (2.3) can be considered in place of only seeking the process *Z*.

Uniqueness of solutions to an SDE can arise in different forms. One such form is pathwise uniqueness, which in the context of strong solutions is defined as:

##### Definition 2.2

**(Pathwise unique strong solutions [33, Section 5.1])**. *Pathwise uniqueness of strong solutions to SDE* (2.3) *holds if for any two strong solutions Z* = (*Z*_*t*_, *t* ≥ 0) *and* 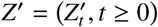 *in the sense of Definition 2.1, we have that*

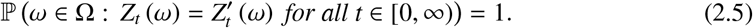

For completeness, establishing the existence and pathwise uniqueness of strong solutions to SDE (2.3) requires certain conditions on *c*_*t*_, *d*_*t*_, and *V*. We state these conditions in Appendix A for ease of reference, since they will be relaxed later in the paper.

#### 2.1.2. Exit times

In certain applications, it may be of interest to determine if, for ℙ-almost every *ω* ∈ Ω, the first time *t*_e_ when a sample path *Z* (ω) exits a certain open set *S* ⊂ S_*Z*_ is finite, or, equivalently, if ℙ-almost surely *t*_e_ = ∞. With respect to our application (Section 3), this information is relevant for establishing whether the solution to an SDE only has positive sample paths; that is, 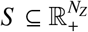.

The Khasminskii test [34] can be used to answer exit-time questions. The test is based on defining (1) an increasing sequence of open sets {*S* _*r*_}, *r* integer, whose closures are contained in *S* and such that their union equals *S*; and (2) an associated sequence of stopping times {*t*_*r*_} with *t*_*r*_ := inf{*t* ∈ [0, *t*_*e*_) : *Z*_*t*_ *S* _*r*_}, where inf ∅ = ∞. By letting *t*_∞_ := lim_*r*→∞_ *t*_*r*_, it can be shown that ℙ-almost surely *t*_∞_ ≤ *t*_e_, whereas to verify if *Z*_*t*_ ∈ *S* always, it suffices to show that no exit occurs at finite times (*t*_∞_ = ∞).

#### 2.1.3. Markovianity of the solution

In this section, the discussion is restricted to Markov processes, whose past and future are statistically independent given the present. If a solution to SDE (2.3) is Markovian, we can describe it in terms of certain transition probabilities. To formally define these notions, we consider the expectation of a stochastic process, conditional on the past information encoded in an appropriate collection of *σ*-fields. Then we have the following definition:

##### Definition 2.3

**(Markov process)**. *Let Z* = (*Z*_*t*_, *t* ≥ 0) *be a measurable process defined on the filtered probability space* (Ω, ℱ, (ℱ_*t*_, *t* ≥ 0), ℙ). *Z is a Markov process relative to* (ℱ_*t*_, *t* ≥ 0) *if for all s* ≤ *t and all bounded measurable functions ϕ, we have* E [*ϕ* (*Z*_*t*_) | ℱ_*s*_] = E [*ϕ* (*Z*_*t*_) | *Z*_*s*_*]. Equivalently, we can say that* ℙ (*Z*_*t*_ ∈ *A*_*Z*_ | ℱ_*s*_) = ℙ (*Z*_*t*_ ∈ *A*_*Z*_ | *Z*_*s*_), *for any A*_*Z*_ ∈ ℬ (S_*Z*_).

In Definition 2.3, by letting *s* = 0, the focus is on expectations that are conditional on the ℱ_0_-measurable *N*_*Z*_-dimensional random variable *Z*_0_, with an interest in the evolution in time of E [*ϕ* (*Z*_*t*_) ℱ_0_]. Without loss of generality, we discuss this evolution for a specific case of SDE (2.3) with no auxiliary process *V*:

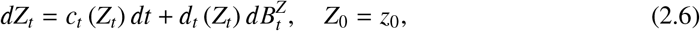

whose solution *Z* = (*Z*_*t*_, *t* ≥ 0) for a deterministic initial condition (the singleton *z*_0_ ∈ S_*Z*_) exists, and is unique and Markov relative to (ℱ_*t*_, *t* ≥ 0) [35, Theorem 17.2.3].

In the following, we make use of the infinitesimal generator *A* = (*A*_*t*_, *t* ≥ 0) of the solution *Z*, that is, the time-dependent second-order differential given by

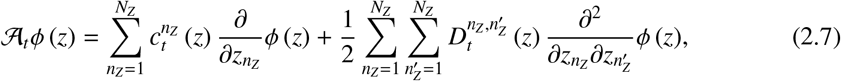

with entries 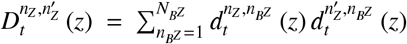 of the diffusion tensor, and where *ϕ* is a suitably regular function of the state.

Furthermore, we construct an equivalent description of *Z* in terms of a Markov transition function ℙ (·,·;·,·) of *s, z*_*s*_, *t* and *A*_*Z*_, defined for 0 ≤ *s* < *t, z*_*s*_ ∈ S_*Z*_, *A*_*Z*_ ∈ ℬ (S_*Z*_). For fixed *s, t*, and *A*_*Z*_, we have that ℙ (*s*, ·; *t, A*_*Z*_) is a measurable function of *z*_*s*_. For fixed *s, t*, and *z*_*s*_, we have that ℙ (*s, z*_*s*_; *t*,) is a probability measure in *A*_*Z*_; that is, ℙ (*s, z*_*s*_; *t, A*_*Z*_) = ℙ (*Z*_*t*_ *A*_*Z*_ *Z*_*s*_ = *z*_*s*_), the probability of *Z* being in *A*_*Z*_ at time *t*, conditional on starting from *z*_*s*_ at time *s* [36, Section 6].

Consequently, using again Definition 2.3, for all bounded measurable functions *ϕ* on S_*Z*_, we have

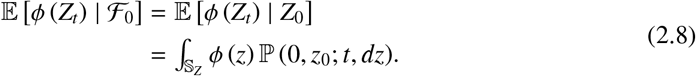

Let *p* (*t*, ·) denote the probability density of ℙ (0, *z*_0_; *t*, ·) with respect to the Lebesgue measure. Applying Itô’s formula and differentiating Eq. (2.8) with respect to time yield an equation for the evolution of E *ϕ* (*Z*_*t*_) | ℱ_0_ :

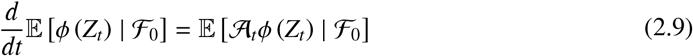

where *A*_*t*_ is the operator presented in Eq. (2.7). Notably, this evolution is deterministic and additional characterisations [37, Section 4] lead to the Kolmogorov forward equation.

#### 2.1.4. The Kolmogorov forward equation (KFE)

In this section, we express expectations of functions of the solution to an SDE in terms of partial differential equations (PDEs). In particular, we present the forward Kolmogorov equation which describes the evolution of the probability density of a process *Z* solving SDE (2.6) by assuming this solution to be a Markov process relative to (ℱ_*t*_, *t* ≥ 0).

Eq. (2.9) can be further developed as

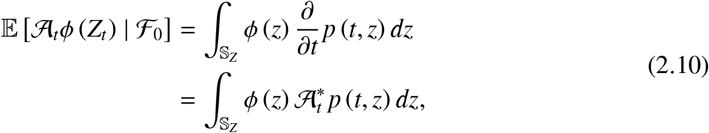

(see, e.g., [35, Section 17]). Then we have the Kolmogorov forward, or Fokker–Planck, equation

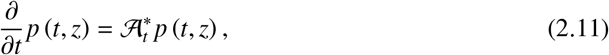

with initial condition *p* (0, *z*) = *p*_0_ (*z*), and where 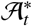 is the adjoint of *A*_*t*_ and is given by

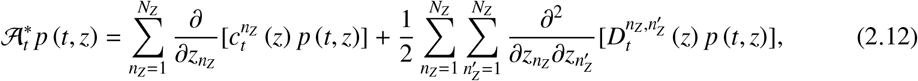

provided *p, c*_*t*_ and *D*_*t*_ are sufficiently smooth.

##### KFE: Solutions

The density *p* associated with a Markov process *Z* which solves SDE (2.6) is a *fundamental solution* to PDE (2.11). That is, with an arbitrary but fixed *T* > 0, for every *s* ∈ [0, *t*), *t* ∈ (0, *T*], *z*_*s*_ ∈ S_*Z*_, *z* ∈ S_*Z*_, and measurable non-negative functions *p*_*s*_ on S_*Z*_, we have:

1. 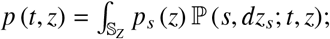
2. lim_*t*→*s*_+ *p* (*t, z*) = *p*_*s*_ (*z*);
3. *p* is continuously differentiable, once in *t* and twice in *z*;
4. *p* satisfies PDE (2.11).

Existence and pathwise uniqueness of a strong solution *Z* of SDE (2.6), along with (A.2), ensure that the fundamental solution *p*, if it exists, is unique [38, Section 5.7]. We refer the reader to [39, Section 6.4] for various regularity and ellipticity conditions on *c*_*t*_ and *D*_*t*_ sufficient to guarantee the existence of a fundamental solution.

The density *p* may also correspond to a *classical solution*. That is, *p* satisfies (2)-(4) above and there exist positive constants *α, β* such that for every *z* ∈ S_*Z*_

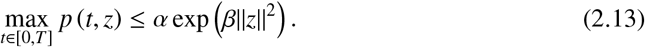

A sufficient condition is that coefficients 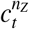 and 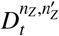 are bounded and Hölder-continuous on S_*Z*_, and the diffusion tensor *D*_*t*_ is uniformly elliptic; that is, for every pair *z, z*^′^ ∈ S_*Z*_, there exists a constant *α >* 0 such that

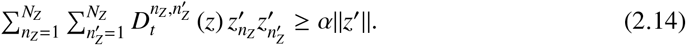

In cases for which only Eq. (2.14) is satisfied, alternative sufficient conditions can ensure the existence of a unique solution with polynomial growth instead. That is, *p* satisfies PDE (2.11) and there exist positive constants *α, β* such that for every *z* ∈ S_*Z*_

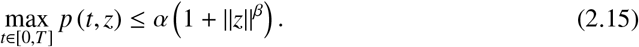

The conditions are that (1) the diffusion tensor *D*_*t*_ is uniformly elliptic, (2) the coefficients 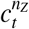 and 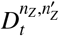 satisfy Eq. (A.2) for compact subsets of S_*Z*_, and (3) the functions 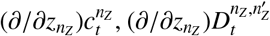 and 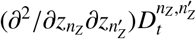 are bounded in S_*Z*_ and Hölder-continuous on compact subsets of S_*Z*_ [39, p. 147].

##### KFE: Boundary conditions

The derivation to show that the adjoint operator 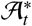 in Eq. (2.10) has coefficients associated with those in SDE (2.6) requires Itô’s formula on a continuous test function *ϕ* on S_*Z*_ with compact support [40]. This ensures that the boundary terms in subsequent integration by parts vanish. The boundary terms also vanish when selecting appropriate conditions at the boundary ∂*S* of a set *S* ⊆ S_*Z*_ inside which *z* is constrained. Boundary conditions (BCs) take on a variety of forms, but note that Eq. (2.11) can always be re-written as

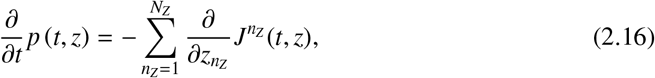

where 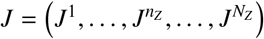 is the probability current with components

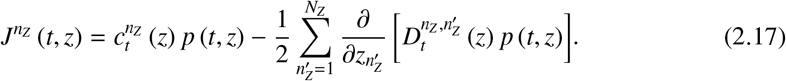

Once *J* has been defined, different BCs can be separately considered. With respect to our application (Section 3), the BCs of interest are

1. Reflecting boundary: *J* (*t, z*) · *n* = 0 for all *t* ∈ [0, ∞), *z* ∈ ∂*S*, and *n* normal to ∂*S*; that is, the net of probability current across ∂*S* is zero;
2. Natural boundary: assume the existence of a *z* ∈ ∂*S* for which *c*_*t*_(*z*) = 0 and *d*_*t*_(*z*) = 0. If process *Z* were to reach *z*, it would remain there. This scenario is called a natural boundary whenever process *Z* never reaches *z*.

### 2.2. Stochastic filtering theory

Thus far, we have only considered the prior evolution of the state process, formally embedded by 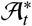 in the KFE. We now incorporate observations and formulate the stochastic filtering problem. This section begins with the introduction of the following processes on Ω: *X* = (*X*_*t*_, *t* ≥ 0), called the state process, and *Y* = (*Y*_*t*_, *t* ≥ 0), the observation process. Similarly to Section 2.1, we let *C*_*Y*_ = *C*_*Y*_ ([0, ∞), S_*Y*_) denote the space of continuous functions on [0, ∞) taking values in 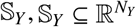, and *D*_*X*_ = *D*_*X*_ ([0, ∞), S_*X*_) denote the space of S_*X*_-valued right-continuous functions on [0, ∞) with left-hand limits, S_*X*_ being a complete separable metric space. For convenience, we let Ω = *D*_*X*_ × *C*_*Y*_. Then any *ω* ∈ Ω can be represented as a pair of sample paths *ω* = (*ω*_*X*_, *ω*_*Y*_), where *ω*_*X*_ ∈ *D*_*X*_ and *ω*_*Y*_ ∈ *C*_*Y*_. Lastly, we introduce the noise processes 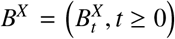 and 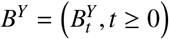. We have not taken into account the existence of *B*^*X*^ and *B*^*Y*^ when constructing Ω, but as discussed in [41], one can extend Ω to include all *ω* associated with the remaining random variables and stochastic processes.

The state process describes the dynamics of a system of interest. It is modelled as a special case of SDE (2.3):

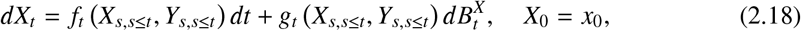

where *X* has sample paths *X* (*ω*) = *ω*_*X*_, *Y* is a stochastic process soon to be introduced, with *Y* (*ω*) = *ω*_*Y*_, process 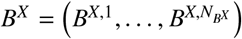 is an 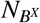-dimensional standard Brownian motion, coefficients *c*_*t*_ and *d*_*t*_ have been respectively renamed *f*_*t*_ and *g*_*t*_, with 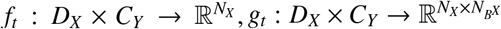, and *X*_0_ is given.

Generally, the state process cannot be directly observed (for instance, only certain components of a vector-valued process are observed). Moreover, observations are prone to error. These two aspects are represented in the following observation model:

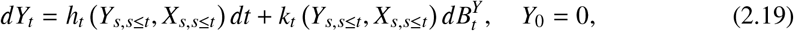

where coefficients *c*_*t*_ and *d*_*t*_ have been respectively renamed *h*_*t*_ and *k*_*t*_, with 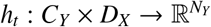 being the observation function and 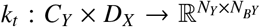 denoting the noise intensity; and the 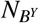-dimensional standard Brownian motion process 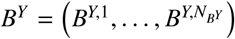 is independent of *B*^*X*^. We let 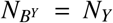. Notably, SDE (2.19) describes the evolution of an integral form of the observations: *Y* is the integral process of the actually observed process 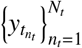, where 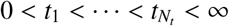 is a uniform partition of a subset of (0, ∞). Therefore, *Y*_0_ being zero does not imply zero-valued observations at *t* = 0, but rather a lack of initial information.

#### 2.2.1. Stochastic equation for the filter

In filtering, we define a stochastic process *π* = (*π*_*t*_, *t ≥* 0) for the conditional distribution of *X*_*t*_ given the observation *σ*-field 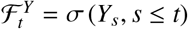:

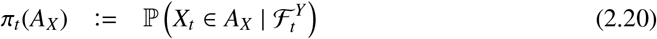

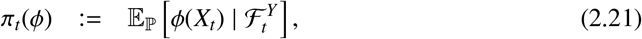

where *A*_*X*_ is an arbitrary set in ℬ (S_*X*_), *ϕ* is a measurable scalar function, and we explicitly show the measure with respect to which expectations are taken. *π*_*t*_ in Eq. (2.20) takes values from ℬ (S_*X*_) into [0, 1]. *π*_*t*_ in Eq. (2.21) gets information about the state through *ϕ*, assuming that the expectation with respect to ℙ exists.

The expectation process *π* (*ϕ*) = (*π*_*t*_ (*ϕ*), *t* ≥ 0), hereafter called filter, satisfies

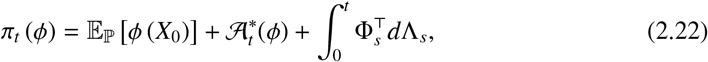

where 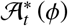 is a measurable process adapted to (ℱ_*t*_, *t* ≥ 0), Φ_*s*_ is a measurable process adapted to ( _*s*_, *s t*), the integration is with respect to (Λ_*s*_, *s ≤ t*), known as the innovation process, and 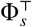 denotes transpose of Φ_*s*_. The required conditions for solvability of Eq. (2.22) for every measurable scalar *ϕ* are stated in [33, Theorem 8.4.1]. Since these conditions rely on additional technical definitions and cannot be easily expressed using the existing notation, they are not explicitly stated here. The remaining problem is to discover the particular form that 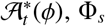 and Λ_*s*_ take in Eq. (2.22) for specific choices of SDEs (2.18) and (2.19).

#### 2.2.2. The Kushner–Stratonovich equation (KSE)

We proceed by assuming the existence of a density *p*_*t*_ = *p* (*t*, ·) of *π*_*t*_ with respect to the Lebesgue measure, for which *π*_*t*_ (*ϕ*) as defined in Eq. (2.21) is obtained via 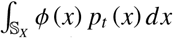. A detailed derivation of the evolution equation for this conditional density is available in [1, 41]. Since the derivation uses arguments which are standard in the theory of Markov processes, we shall state only the outline. Two common approaches for deducing this evolution equation are the change of measure and innovation process methods. We present the former method in the Supplementary Material following similar steps as in [42]. For the latter, the interested reader can refer to [33].

Let us assume that the pair (*X, Y*) solves a special case of SDEs (2.18) and (2.19):

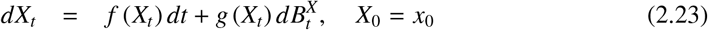

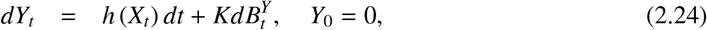

that is, the case in which *f*_*t*_ = *f, g*_*t*_ = *g* and *h*_*t*_ = *h* are autonomous functions of the state *X*_*t*_. Since the coefficients are autonomous, the infinitesimal operator and its adjoint become time-independent, denoted as A and A∗, respectively. The observation process is driven by additive noise, i.e. *k*_*t*_(*X*_*s,s*≤*t*_, *Y*_*s,s*≤*t*_) = *K*, where *K* is a matrix of size *N*_*Y*_ × *N*_*Y*_. Notably, the filtering problem whose observation model is driven by multiplicative noise can be converted into a problem whose observation model is driven by additive noise, via a suitably chosen stochastic flow mapping [43].

We define the process *L* = (*L*_*t*_, 0 ≤ *t* < ∞) by

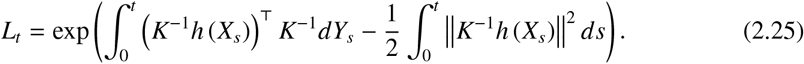

Under certain technical conditions (see Appendix B), Eq. (2.25), along with Itô’s formula, leads to the following proposition:

##### Proposition 2.1

**(Kushner [44], Stratonovich [45])**. *Let* ϕ *have continuous partial derivatives of every order up to 2, all of which are bounded. Then we can write*

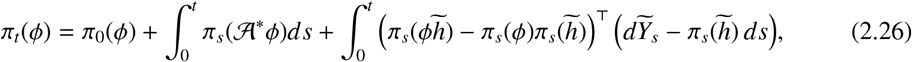

*where π*_0_ (*ϕ*) = E_ℙ_ *ϕ* (*X*_0_). *Hereafter* 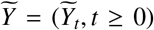 *is the rescaled observation process (i.e*. 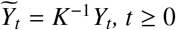*), and* 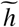 *is the rescaled observation function (i.e*. 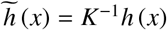).

*Furthermore, an application of integration by parts to Eq*. (2.26) *results in the so-called Kushner–Stratonovich equation (KSE)*

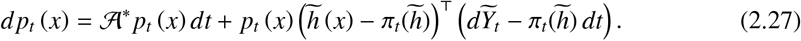

Proposition 2.1 concerns the evolutions of *π* and *p* in the special case of Eq. (2.22) for which 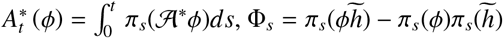, and 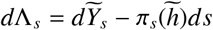. We refer the reader to [33] for the detailed derivations of Eq. (2.26) and PDE (2.27). For completeness, we outline the derivation using *L*_*t*_ from Eq. (2.25) in the Supplementary Material.

##### KSE: Pathwise filtering and solutions

So far, *π*_*t*_(*ϕ*) has been defined on an unspecified set in ℬ (*C*_*Y*_) of measure one. We have not yet shown *π*_*t*_(*ϕ*) in the case of a given rescaled observation path 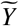 (*ω*). To this end, we present the notion of pathwise filtering from [4, 46].

*L*_*t*_ in Eq. (2.25) is written in terms of a stochastic integral with the process 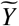 as the integrator. However, since the integrand is independent of the observations, we reverse the role of processes *X* and 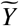: up to some finite time *t, L*_*t*_ is obtained by an integral whose integrator is (*X*_*s*_ (*ω*), *s* _≤_ *t*). By integration by parts,

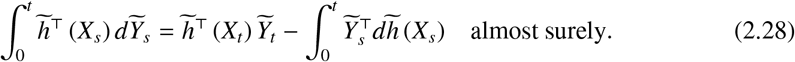

The pathwise counterpart to Eq. (2.25) is then

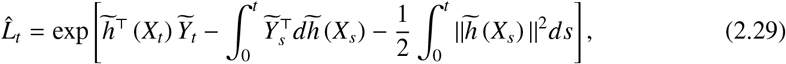

where 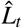 corresponds to *L*_*t*_ when computed with respect to a given observation path.

The KSE in Proposition 2.1 is based on the Radon–Nikodym derivative *L*_*t*_ from Eq. (2.25). Similarly, a pathwise KSE can be derived based on 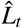 from Eq. (2.29). This reduces the stochastic PDE (2.27) to a PDE with a random coefficient:

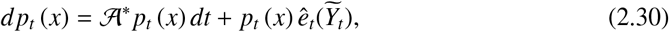

where *ê*_*t*_ : S_*Y* →_ ℝ is related to 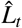 (see [5, 41]). Once 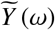 has been fixed, a solution to PDE (2.30) can be characterised as in Section 2.1.4, under additional regularity conditions on *h*, such as continuity and boundedness. Hereafter, *p* = (*p*_*t*_, *t* ≥ 0) refers to this pathwise solution.

##### KSE: Pathwise Approximations to Solutions

This section contains an overview of two classes of methods to approximate *p*. We briefly describe each class and state related results.

Let the uniform partition of the interval [0, ∞) have time step 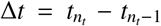. Given a set of rescaled observations 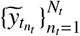, the density 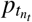 is approximated by 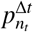 by first defining 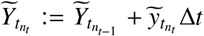. This corresponds to a continuous-discrete filtering problem, in which the observation process 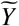 from the continuous formulation is interpolated by straight-line segments between discrete observation times. Secondly, 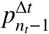 is propagated to 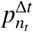 according to one of the following methods:

- Particle filtering
- Finite-difference + splitting-up approximation

##### Particle filtering

The density 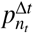 is constructed by density estimation techniques [47], such as histograms or kernel estimation, over a finite number of samples updated sequentially up to time 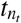. The following exposition is adapted from [1]:

Let 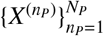 be a collection of *N*_*P*_ mutually independent stochastic processes, each solving SDE (2.23) and independent of 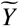. Then the pairs 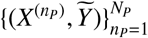 are identically distributed.

Additionally, each pair is associated with a suitably chosen weight 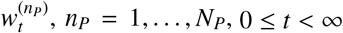, such that the triples 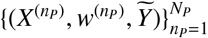 are identically distributed and have the same distribution as the triple 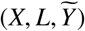 under a suitable probability measaure. We refer the reader to the Supplementary Material for the full characterisation of 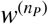.

Lastly, the *N*_*P*_ triples are used to construct the measure-valued process 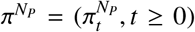 defined as 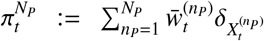, where 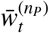 is given by 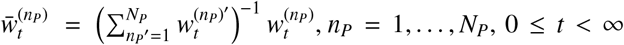, and *δ* is the Dirac delta generalised function; that is, 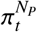 is the empirical normalised measure of *N*_*P*_ particles with positions 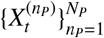. Results on the convergence to *π*_*t*_ are detailed in, for instance, [1].

##### Finite-difference + splitting-up approximation

stochastic PDEs are solved using tools inherited from solving classical PDEs. The following exposition follows the description given in [48]:

The transition from 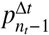 to 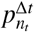 follows a splitting-up approximation [13, 14], whose first step, called the *prediction* step, consists in solving the KFE (2.11) for a single time step Δ*t*. In other words, we solve

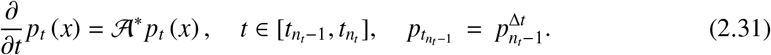

We then define 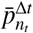 as the solution evaluated at 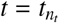, i.e. 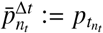.

The second step, called the *correction* step, uses Eq. (2.29) and the new rescaled observation 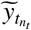 to update 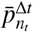. We obtain 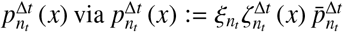, where 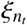 is a normalising constant chosen such that 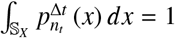, and

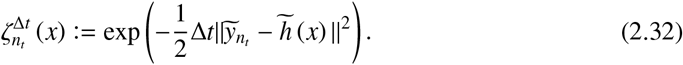

Results concerning the convergence to *p* = (*p*_*t*_, *t* ≥ 0) are detailed in, for instance, [49].

## 3. The chemostat

Let us consider a bioreactor in which one or more populations of microorganisms grow in a nutrient medium consisting of a cocktail of molecules. The chemostat (Figure 1) is a laboratory device used to represent such a bioreactor [50]. It is widely used in experimental design due to its ability to provide controlled conditions and reproducible results [26].

**Figure 1:**
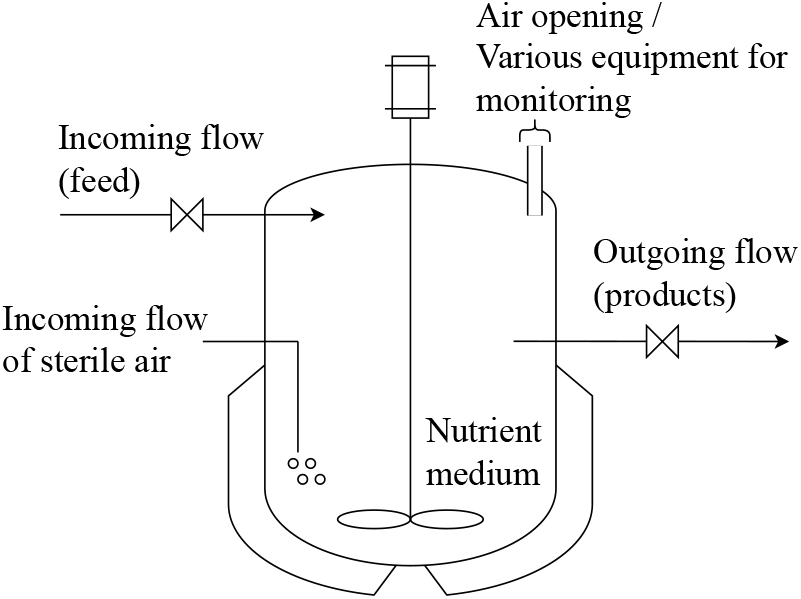
Schematic representation of a chemostat.

In this work, biomass refers to a single population of growing microorganisms and substrate to the cocktail of molecules. The production of biomass may also generate the byproducts methane and carbon dioxide, collectively referred to as biogas. Under this scenario, the following reaction takes place to produce one unit of biomass and *γ* units of biogas from *κ* units of substrate:

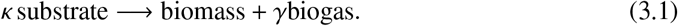

We begin with a differential description of the substrate and biomass concentrations in a chemostat, based on reaction (3.1).

Let *b* = (*b*_*t*_, *t* ≥ 0) and *s* = (*s*_*t*_, *t* ≥ 0) denote, respectively, the time evolution of the concentrations of biomass and substrate (hereafter, *s* refers to a concentration process, rather than a time variable as in Section 2). Let *x* = (*x*_*t*_, *t* ≥ 0) = (*b, s*) denote the state process. Then the differential equation for the dynamics of the chemostat is

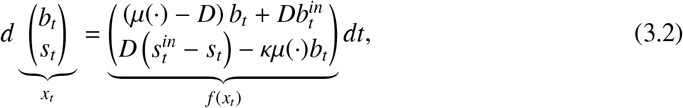

with initial concentrations 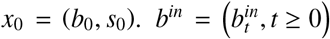 and 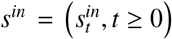 are incoming concentrations, *D* is the dilution rate (that is, the volumetric flow rate divided by the bioreactor volume), and *µ* is the growth rate, a function modelling the growth of a population and ensuring a zero reaction rate in the absence of substrate. The derivation of ODE (3.2) is provided in the Supplementary Material.

The operation of the chemostat can be summarised as follows: the inputs comprise the dilution rate *D* together with the incoming concentrations *b*^*in*^ and *s*^*in*^. Among these inputs, *D* is typically the manipulated variable. Under the perfect-mixing assumption, the outgoing flow in Figure 1 carries the same biomass and substrate concentrations, *b* and *s*, as those within the bioreactor. The online characterisation of *b* via direct observations is generally unavailable or expensive, whereas that of *s* is generally more accessible, although observations are typically available only intermittently (e.g. every few minutes). On the other hand, the biogas flow rate is routinely measured, and can therefore provide information for process supervision and feedback control [51, 52]. Combined with biogas flow-rate observations, reconstructed values of *s* may be used in feedback laws for selecting the dilution rate that stabilises the chemostat around a desired operating point [53]. Furthermore, reconstructed values of *b* can be exploited in control strategies aimed at maximising biogas production [54] or for the early detection of abnormal operating conditions.

For instance, a persistent decrease in the estimated biomass concentration may indicate the onset of biomass washout, prompting an adjustment of the dilution rate. Conversely, persistently high estimates of biomass concentration may indicate excessive biomass accumulation, triggering control actions to maintain stable bioreactor operation.

This work assumes no incoming flow of biomass; thus 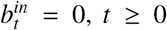. Moreover, we proceed with a simplified version of ODE (3.2) called the minimal model [25], in which (i) *µ* is a function of the substrate, with *µ*(*s*) > 0 for *s* > 0 and *µ*(0) = 0, and (ii) the yield parameter *κ* is constant. These model assumptions are to be interpreted in the following way: (i) the velocity—or kinetics—of the reaction in Eq. (3.1) does not change as the biomass concentration changes and (ii) temperature and pressure are kept constant, thus *κ* is also constant.

Under these model assumptions, we now introduce two cases for which different types of organisms are being cultivated:

- Case 1 - Monotonic growth *µ* (Monod kinetics)
- Case 2 - Non-monotonic growth *µ* (Haldane kinetics)

**Case 1:** the Monod growth law [55] assumes that the growth rate is zero when there is no substrate, and tends to an upper limit when the substrate is in excess:

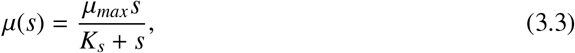

where *µ*_*max*_ is the maximum growth rate and *K*_*s*_ is the half-saturation constant.

**Case 2:** some experiments indicate that excessive substrate may inhibit the growth of a population [56]. This scenario is represented by the following growth function:

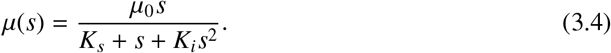

Here, *µ* depends on the growth *µ*_0_, half-saturation *K*_*s*_ and inhibitor parameter *K*_*i*_.

Figure 2 illustrates the phase portraits corresponding to these two cases and some highlighted phase paths for ODE (3.2) under the parameters from Table 1.

**Table 1:**
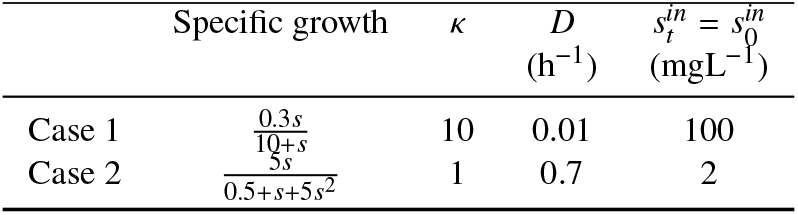
Model parameters for ODE (3.2).

**Figure 2:**
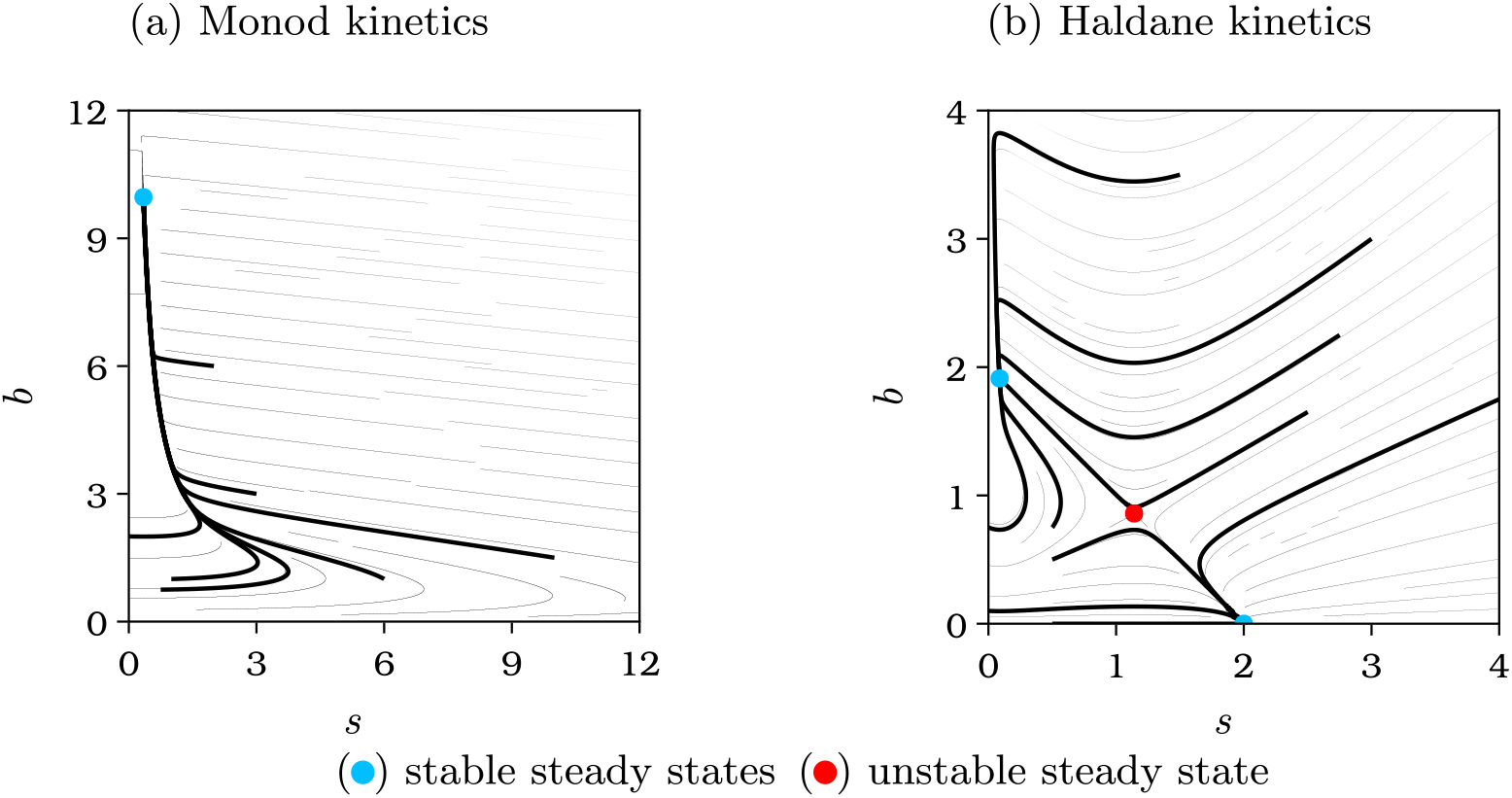
Phase portrait for ODE (3.2) with growth functions (a) (3.3) and (b) (3.4).

The coloured points in Figure 2 are equilibrium pairs (*b*_*e*_, *s*_*e*_) for which the right-hand side of ODE (3.2) is zero. Case 1 has the equilibrium point 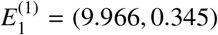, known to be locally exponentially stable [25]. Case 2 has a trivial equilibrium point 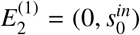. Additionally, it has interior equilibria 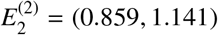 and 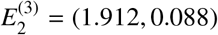. It can be shown that one interior equilibrium is a stable node and the other a saddle point [25].

Having introduced the chemostat model, we are ready to apply the concepts from Section 2 to this system. Let us assume that the concentration of an element changes within a time period by a random amount, and that this random amount is a function of the current concentration itself. This model assumption has been discussed in [27], for which the equation governing the dynamics, in the form of SDE (2.23), is

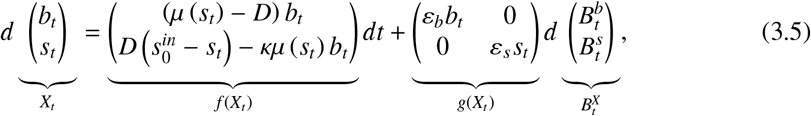

where 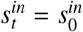 was held constant, and *X*_0_ = (*b*_0_, *s*_0_).

Figure 3 presents numerical solutions of SDE (3.5) for Cases 1 and 2. The noise intensities (*ε*_*b*_, *ε*_*s*_) are (0.05, 0.05) for Case 1 and (0.075, 0.075) for Case 2.

**Figure 3:**
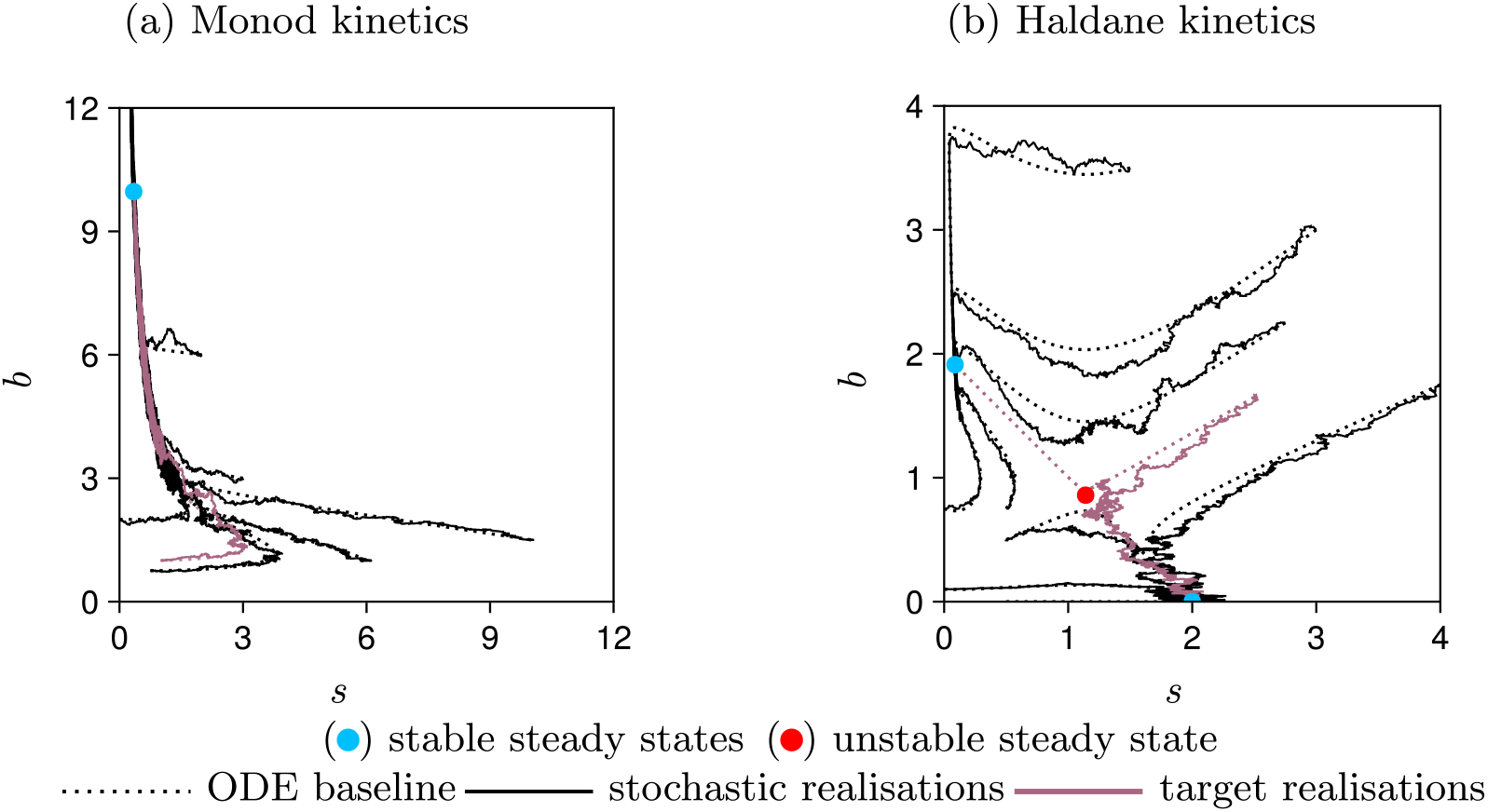
Solutions of SDE (3.5) with growth functions (a) (3.3) and (b) (3.4). The dashed lines show the solutions of ODE (3.2), as in Figure 2. A target realisation refers to a trajectory to be estimated from observations later in this section.

Figure 3(a) suggests that the processes *X* move around the trajectories of the deterministic chemostat model, with small fluctuations. Here, 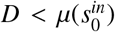. Under the deterministic model, this condition implies that the microbial population forming the biomass survives regardless of the initial concentrations of substrate and biomass [25]. In Figure 3(b), the processes *X* also move around the trajectories of the deterministic chemostat model, but in order to avoid “near” extinction of the population around 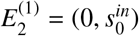, we would have to appropriately choose *X*_0_.

Our main contributions are presented in the following two sections. Section 3.1 establishes some theoretical guarantees for the stochastic chemostat. In particular, we prove in Section 3.1.1 the existence, uniqueness and non-explosion of the solution *X*, as introduced in Sections 2.1.1 and 2.1.2. Section 3.1.2 revists the Markovianity of the solution and the KFE from Sections 2.1.3 and 2.1.4. Section 3.2 particularises the filtering equations to the case in which the pair (*X, Y*) is Markovian, with *Y* corresponding to possible observation processes for a chemostat. Finally, we present numerical approximations of the solution to the KSE of Section 2.2.2 based on the computational methods previously outlined.

### 3.1. Theoretical guarantees

#### 3.1.1. Existence, uniqueness and non-explosion of the solution

While numerical simulations are valuable for exploring the behaviour of the chemostat, proofs of existence and uniqueness of solutions of SDE (3.5) are required to ensure the validity of any conclusions drawn from Figure 3. Moreover, as described in Section 2.1.2, we question whether a sample path of *X* almost surely exits an open set *S* in finite time. By letting 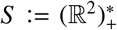, i.e. *S* = {(*b, s*) ℝ^2^ *b* > 0, *s* > 0}, we investigate whether *X* explodes or reaches zero/negative values in finite time. This checks for unrealistic scenarios in a chemostat, such as negative concentrations.

Since Condition A does not hold for SDE (3.5), Theorem A has to be modified to prove the existence and uniqueness of solutions. By following [30], we present the necessary modifications and show that a unique solution *X* only exits *S* at infinite time.

##### Theorem 3.1.

*For any initial condition X*_0_ ∈ *S, there exists a unique strong solution X of SDE* (3.5) *with growth functions* (3.3) *or* (3.4). *Futhermore, X only has positive sample paths*.

##### Proof 3.1.

*We begin by showing that the coefficients f and g of SDE* (3.5) *satisfy the local Lipschitz continuous conditions. The diffusion coefficient g is affine and therefore globally Lipschitz. Furthermore, both the Monod and Haldane growth functions are continuously differentiable on* (0, ∞). *Consequently, the drift coefficient f, being composed of sums and products of continuously differentiable functions, is continuously differentiable and is therefore locally Lipschitz on S (by the mean value theorem). By the standard local existence and uniqueness theorem for SDEs with locally Lipschitz coefficients (see, e.g*., *[34, 57]), there exists a unique solution X* = (*X*_*t*_, *t* ≥ 0) *of SDE* (3.5) *on the stochastic interval* [0, *t*_*e*_), *where X*_0_ ∈ *S and*

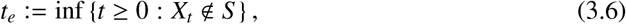

*again with the convention that* inf ∅ = ∞.

*To prove that the solution is global, it suffices to show that t*_*e*_ = ∞ *almost surely. Let r*_0_ *be a sufficiently large integer such that* 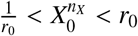, *n*_*X*_ ∈ {1, 2}. *For each integer r* ≥ *r*_0_, *we define an increasing sequence of open sets* 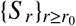, *where S* _*r*_ := (1/*r, r*) × (1/*r, r*), *so that S* _*r*_ ⊂ *S* _*r*+1_ *and* 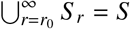. *We further define the stopping times*

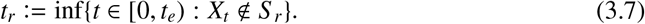

*Since* 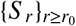 *is an increasing sequence*, 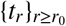 *is non-decreasing. Hence*,

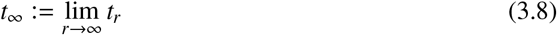

*is well-defined and satisfies t*_∞_ ≤ *t*_*e*_ *almost surely*.

*We proceed by contradiction. Suppose that*

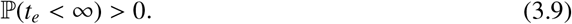

*Since t*_*r*_↑ *t*_∞_, *it follows that* ℙ(*t*_∞_ < ∞) > 0. *Therefore, there exist positive constants T, β, and an integer r*_0_ *such that*

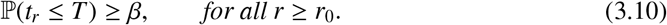

*We will show that this assumption leads to contradiction. To this end, we construct a Lyapunov function that diverges whenever either state variable approaches the boundary of S*.

*Let us define*

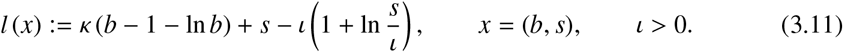

*The non-negativity of l follows from the inequality α ≥* 1 + ln(*α*) *for all α >* 0. *It is sufficient to consider the Haldane growth function, since the Monod growth function is recovered by setting K*_*i*_ = 0. *Let X*_0_ = *x*_0_, *with l*(*x*_0_) *<* ∞. *Applying Itô’s formula to the process l* (*X*) *yields*

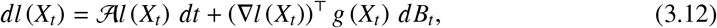

*where A denotes the infinitesimal generator presented in Eq*. (2.7) *and associated with SDE* (3.5). *Therefore, for the first term on the RHS*,

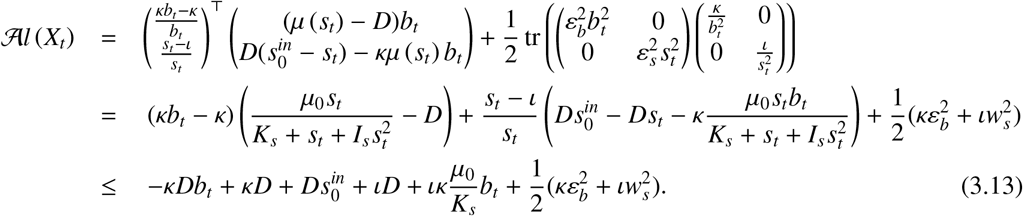

*If we let* 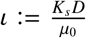, *the terms depending on b*_*t*_ *cancel. Consequently, there exists a constant α >* 0 *such that* 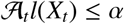, *and therefore*

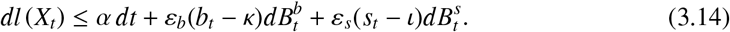

*Let t*_*r*_ ∧ *t* = min{*t*_*r*_, *t*}. *Integrating inequality* (3.14) *over the interval* [0, *t*_*r*_ ∧ *T*] *gives*

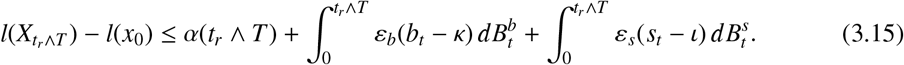

*Taking expectations and using that both stochastic integrals are martingales with zero expectation yields*

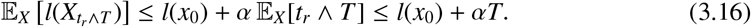

*Also, for every t*_*r*_ ≤ *T, there exists some n*_*X*_ ∈ {1, 2} *such that either* 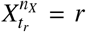 *or* 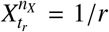. *Hence*, 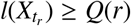, *where*

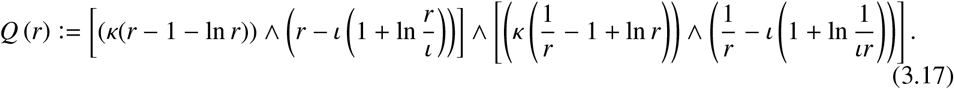

*Therefore*,

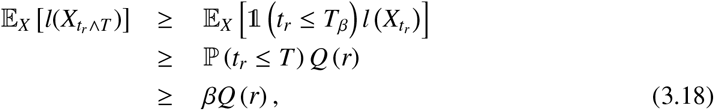

*Combining* (3.16) *and* (3.18) *yields*

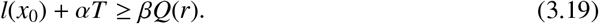

*Since Q*(*r*) → ∞ *as r* → ∞, *we obtain l*(*x*_0_) + *αT* = ∞, *which contradicts the finiteness of both l*(*x*_0_) *and T. Therefore*, ℙ(*t*_*e*_ *<* ∞) = 0, *so that t*_*e*_ = ∞ *almost surely. Hence the local solution extends to a global solution whose sample paths remain in S*.

##### Remark 3.1.

*[58] proposes alternative driving processes to randomise the deterministic dynamics of the chemostat: for instance, a Brownian motion term* 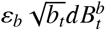 *instead of* 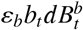, *and* 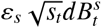 *instead of* 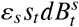. *As detailed in their paper, this induces minimal change in the case of large population sizes. However, these driving processes differ drastically in the washout regime, i.e. near the boundary b*_*t*_ = 0, *as the two diffusion processes exhibit different behaviour. Theorem 3.1 assures that X remains within S, thus the washout regime is unattainable in SDE* (3.5). *If X is driven by* 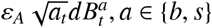, *then not only is b*_*t*_ = 0 *attainable, but so is s*_*t*_ = 0, *for certain regions of the parameter space. Care needs to be taken to avoid the unrealistic scenario of a vanishing substrate concentration, despite the continuous substrate inflow s*^*in*^.

#### 3.1.2. Markovianity and the KFE

Due to the Markovianity of the unique strong solution *X* of SDE (3.5), the evolution of *X* can be studied through its associated KFE, as in [23]. By solving the KFE for a given initial distribution, we examine the probability that the state of the chemostat lies within regions of interest, e.g. around 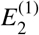 or 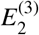.

Notably, the results from Theorem 3.1 alone are insufficient to ensure that the fundamental and classical solutions to the KFE discussed in Section 2.1.4 exist and are unique. For instance, the coefficients of SDE (3.5) do not satisfy Condition (A.2), and hence the sufficient condition for the uniqueness of a fundamental solution according to [38, Section 5.7] is not satisfied. Moreover, the diffusion tensor associated with SDE (3.5)

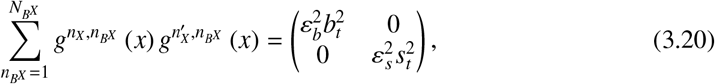

herein denoted by *G*, does not satisfy the uniform ellipticity condition (2.14). Consequently, we are unable to show the existence of a unique solution *p* satisfying the associated KFE with exponential (2.13) or polynomial (2.15) growth.

However, the existence of a unique solution *p* can be established under broader assumptions on *f* and *G*, along with certain properties of a Lyapunov function *l*. We present these assumptions and verify their validity for the stochastic chemostat. With the uniqueness of the solution to the KFE ensured, we present its numerical solution.

First, let us consider a local boundedness condition on coefficient *f*_*t*_:

***Condition A***. For all positive integer *r, f* ∈ ℒ^∞^ (*S* _*r*_). That is, *f* is an essentially bounded measurable function on the open set *S* _*r*_.

Secondly, let us consider a relaxed version of inequality (A.1) for coefficient *G*:

***Condition B***. For all positive integer *r*, there exists a constant *γ*_*r*_ *>* 0 such that

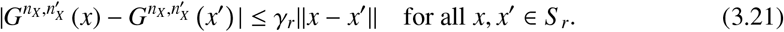

Lastly, let us consider a relaxed version of inequality (2.14), again for *G*:

***Condition C***. For every *S* _*r*_ ⊂ *S*, there exist positive constants *α*_*r*_ and *β*_*r*_ such that

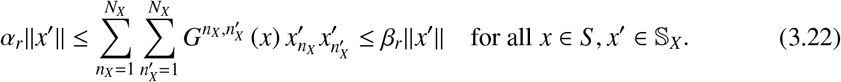

These conditions yield the following result for existence and uniqueness:

##### Proposition 3.1.

*There exists exactly one probability measure whose density with respect to the Lebesgue measure satisfies the KFE associated with SDE* (3.5).

##### Proof 3.2.

*Let us note that the coefficients of SDE* (3.5) *satisfy conditions A, B, and C (by letting* 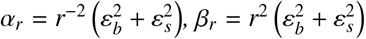, *and* 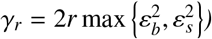.

*According to [59, Theorem 3.6], such conditions are sufficient to ensure the existence of a unique solution, given that there exists a Lyapunov function l such that*

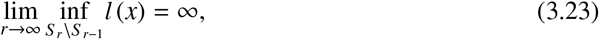

*and for all t* ∈ [0, ∞), *x* ∈ *S, and some α >* 0, *the following inequality holds:*

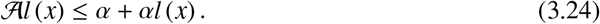

*If we define* 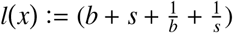, *then* 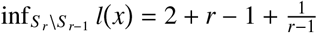 *and nd Al (x) ≤ α + αl ( x), where* 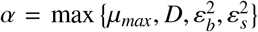 *is an upper bound for the growth function µ. Hence, Eq*. (3.23) *and inequality* (3.24) *hold, and the proof is complete*.

To conclude this section, we present numerical solutions of the KFE for Cases 1 and 2 in Figure 4. These simulations complement the theoretical results established above by illustrating the evolution of the state density. They are obtained using tools similar to those in [23]: an upwind differencing scheme for the space discretisation, an implicit scheme for the time discretisation, reflecting boundaries along the truncated discretisation of the positive quadrant, and natural boundaries along the coordinate axes at which one of the state variables vanishes. The upwind differencing scheme is chosen to ensure the non-negativity of probabilistic terms, as discussed in [60]. Our approach differs from that of [23] by replacing the implicit Euler scheme with the Crank-Nicolson scheme as it led to improved stability.

**Figure 4:**
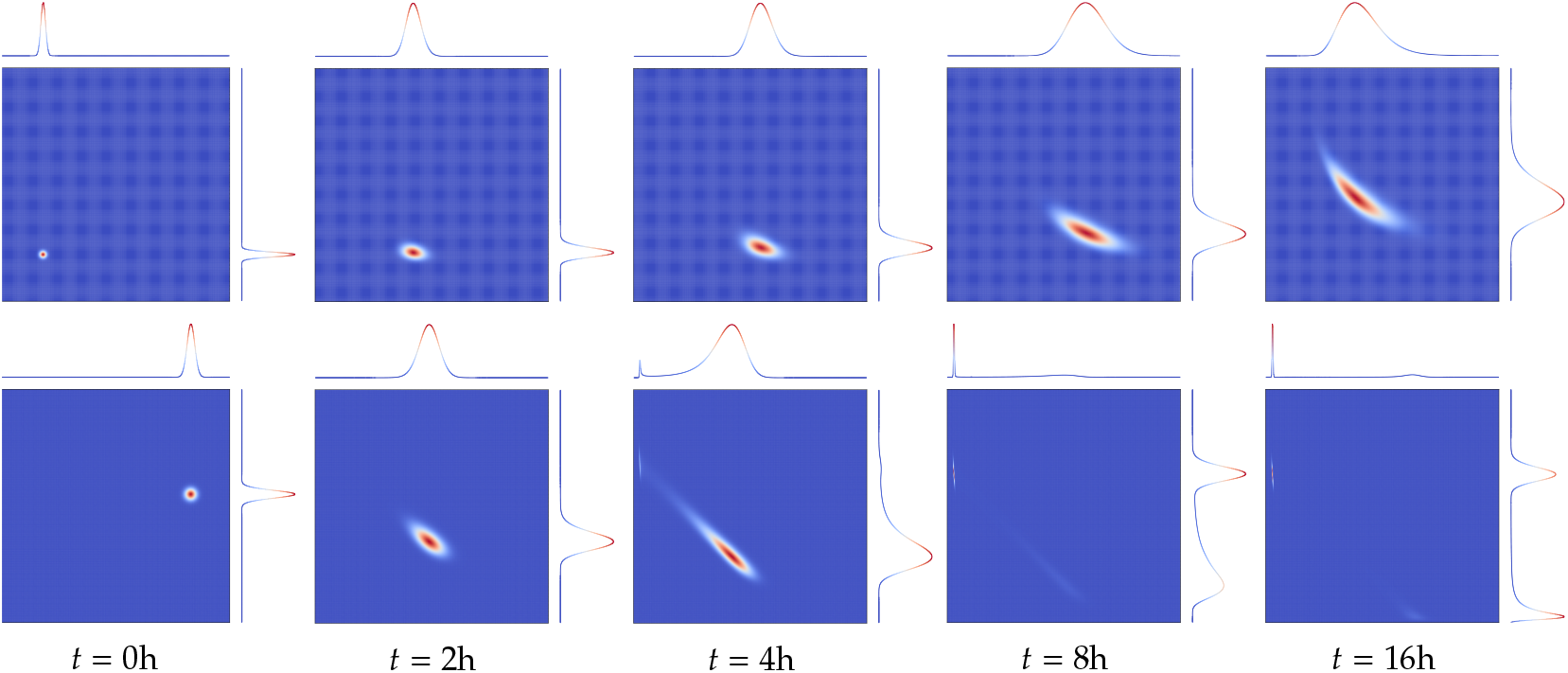
KFE solutions for Monod growth (Case 1, top) and Haldane growth (Case 2, bottom). Case 1: *X*_0_ ~ *N*([1, 1]^⊤^, 0.05^2^*I*_2×2_), the spatial discretisation is performed over the domain (0, 5] × (0, 5], using 256^2^ finite volumes. Case 2: *X*_0_ ~ *N* ([1.65, 2.5]^⊤^, 0.05^2^*I*_2×2_); the spatial discretisation is performed over the domain (0, 3] × (0, 3], using 512^2^ finite volumes. The marginal density functions are displayed as marginal curves, normalised by their maximum value for the sake of illustration.

In Figure 4 (top), the density remains unimodal, while exhibiting a relatively high spread over time due to diffusion. Notably, the state distribution departs from Gaussianity, as can be verified by computing its higher-order moments. In Figure 4 (bottom), the probability mass gradually splits into two regions centred around the stable equilibria. In both cases, effective control of the chemostat may therefore require reducing the uncertainty in the state location.

### 3.2. Filtering problem for the stochastic chemostat

As discussed in Section 2.2, the state process cannot be directly observed. We introduce an observation model in which a sensor transforms the state variables in a univariate signal prone to error. Specifically, we assume noisy observations of the biogas flow rate produced by the biomass after substrate consumption. This flow, proportional to the microbial activity [61, 62], is modelled by the mapping (*s, b*) → *γµ* (*s*) *b*, where *γ* is the yield parameter for biogas in Eq. (3.1). This setting is denoted by Scenario A.

Written in the form of SDE (2.24), the observation process evolves according to

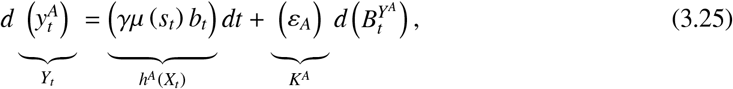

where the superscript *A* refers to Scenario *A*, and *Y*_0_ = 0.

We combine Cases 1 and 2 with SDE (3.25) to design the experiments in Table 2. Over a time interval of interest [0, *T*], we simulate trajectories of *X* and *Y* using the time discretisations Δ*t*^*x*^ and Δ*t*^*y*^, respectively. The trajectories of *Y* are illustrated in the Supplementary Material.

**Table 2:** Additional parameters for differential equations of *X* and *Y*.

|  | Model parameters |  | Simulation parameters |  |
| --- | --- | --- | --- | --- |
| | Observation function | $\varepsilon_A$ | $\Delta t^x$<br>(hours) | $\Delta t^y$<br>(hours) |
| 1A - high-frequency | $27.5\mu(s)b$ | 0.01 | 0.01 | 0.01 |
| 1A - low-frequency | $27.5\mu(s)b$ | $0.01 \left(\frac{\Delta t^y}{\Delta t^x}\right)^{\frac{1}{2}}$ | 0.01 | 2 |
| 2A - high-frequency | $2.75\mu(s)b$ | 0.01 | 0.005 | 0.005 |
| 2A - low-frequency | $2.75\mu(s)b$ | $0.01 \left(\frac{\Delta t^y}{\Delta t^x}\right)^{\frac{1}{2}}$ | 0.005 | 2 |

The observations 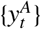 are assumed to be collected either at a high frequency (Δ*t*^*y*^ = Δ*t*^*x*^) or at a low frequency (Δ*t*^*y*^ ≫ Δ*t*^*x*^). The initial pair (*b*_0_, *s*_0_) is (1 mgL^−1^, 1 mgL^−1^) for Case 1 and (1.65 mgL^−1^, 2.5 mgL^−1^) for Case 2.

In addition to Scenario A, the Supplementary Material presents the filtering problem for Cases 1 and 2 under three additional scenarios: Scenario B, with noisy observations of the substrate concentration at a low frequency; Scenario C, with noisy observations of the biomass concentration at a low frequency; and Scenario D, with noisy observations of both biomass and substrate concentrations at a low frequency.

Without the observations, the probability density of the hidden state evolved according to KFE (2.11) as illustrated in Figure 4. In light of the observations, we can now approximate the solution of the pathwise version of KSE (2.27) for all the experiments in Table 2. Figures 5 and 6 show the pathwise solution 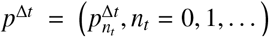 for low-frequency observations Δ*t* = Δ*t* ≫ Δ*t*) via the splitting-up approximation described in Section 2.2.2.

**Figure 5:**
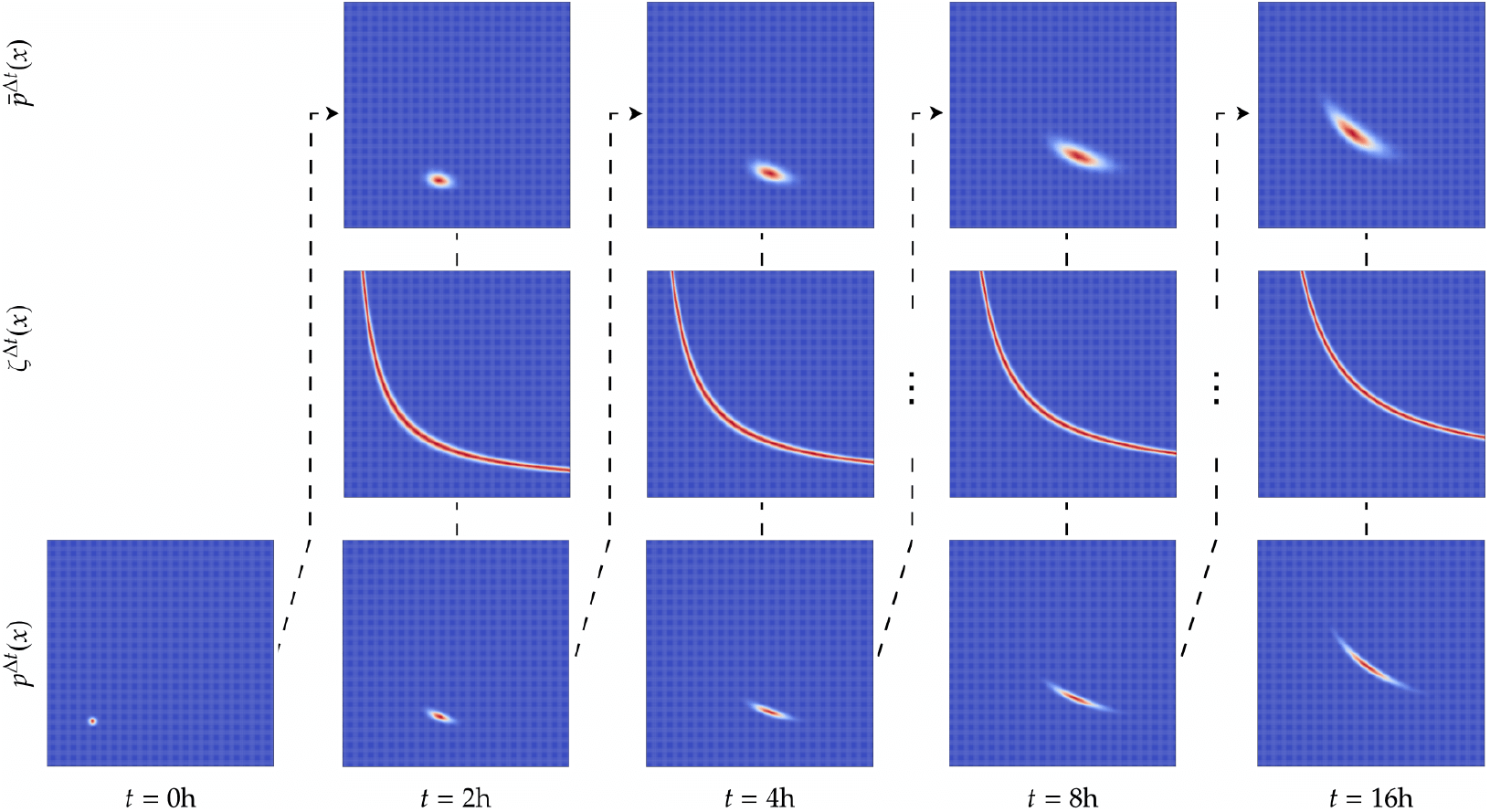
Splitting-up approximation for Case 1A - low-frequency observations.

**Figure 6:**
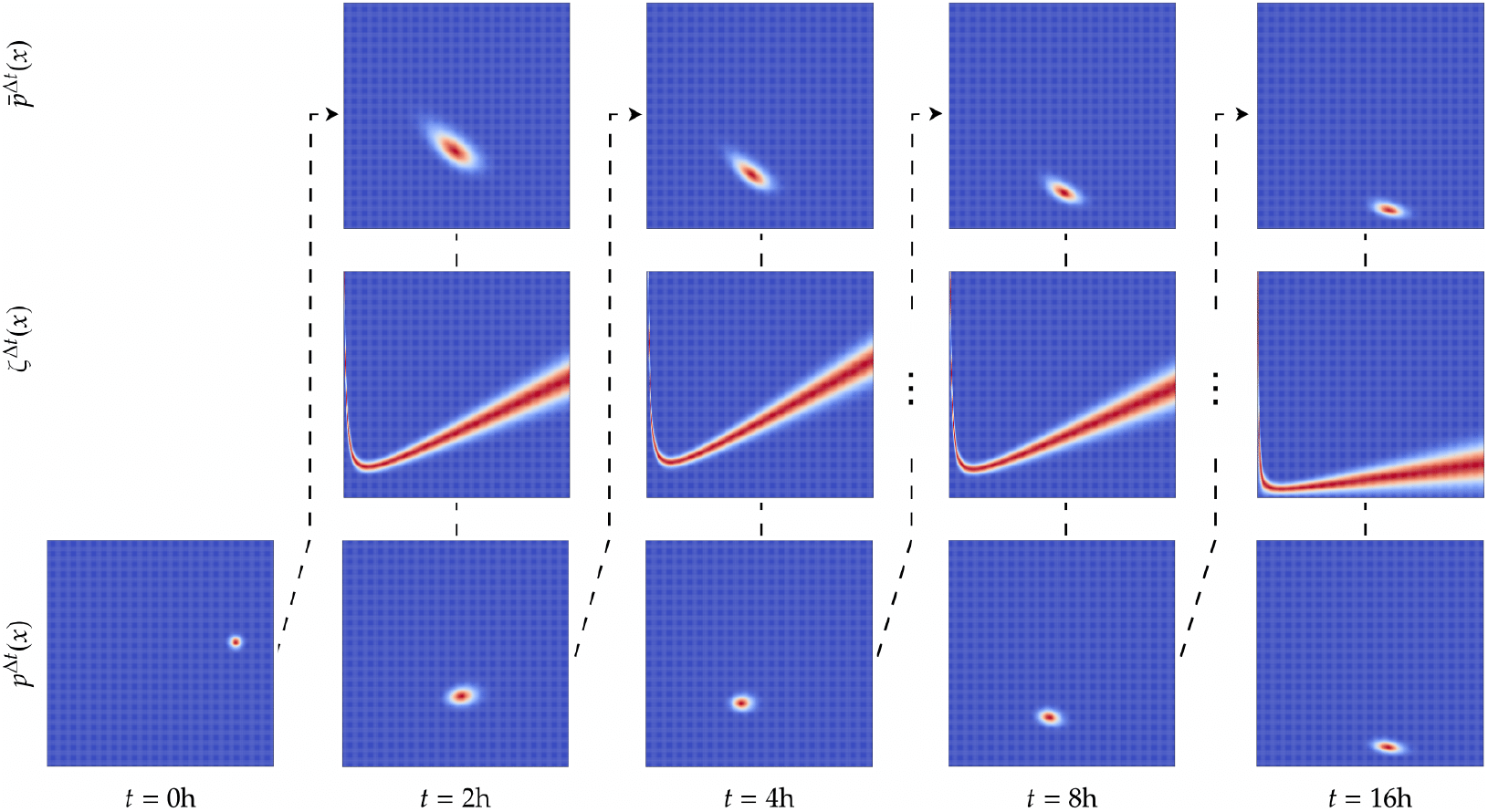
Splitting-up approximation for Case 2A - low-frequency observations.

Each column in Figures 5 and 6 illustrates the iterative steps of the splitting-up approximation. The top row shows the result from the prediction step based on the KFE (2.31), the middle row depicts the likelihood ratio from Eq. (2.32), and the bottom row presents the normalised product of these two, i.e. the corrected density.

We approximate these solutions via two other methods: particle filtering, discussed in Section 2.2.2, and hereafter denoted by bootstrap particle filter (BPF), and the linearisation method known as extended Kalman filter (EKF). The experiments were run in a Precision 5820 workstation (Intel Xeon W-2245 (8C) @ 3.9GHz, 64GB of RAM).

To quantify the filtering accuracy, we compare the estimated conditional mean, *π*_*t*_(*X*_*t*_), with the true simulated state, *X*_*t*_, using the Integral of Time-weighted Absolute Error (ITAE, [63]), which provides a single summary measure of the estimation error over each simulation. Smaller values of ITAE correspond to more accurate reconstructions of the state.

Table 3 compares the filtering methods in terms of the ITAE metric and the average runtime per iteration. The PDE-based approximation generally benefits from mesh refinement, at the expense of higher computational cost. Likewise, the BPF exhibits improved accuracy as the number of particles increases. For the Monod case, the EKF provides the lowest runtime but with slightly lower estimation accuracy than the best-performing PDE and BPF configurations. Conversely, for the Haldane case, the finest PDE discretisation considered here (128^2^) still underperforms both the BPF and the EKF. This indicates that additional mesh refinement is required, a point that is confirmed by the subsequent experiments, in which a 512^2^ grid is shown to outperform the EKF.

**Table 3:** Comparison of the filtering methods in terms of the ITAE metric and average runtime in milliseconds per iteration for the Monod and Haldane kinetics over 20 independent trajectories with high-frequency observations.

| Method | Case 1A (Monod) |  | Case 2A (Haldane) |  |
| --- | --- | --- | --- | --- |
| | ITAE<br>$T = 16$ h | Runtime<br>ms/iter | ITAE<br>$T = 16$ h | Runtime<br>ms/iter |
| KSE-PDE ($32^2$) | 176.37 | 9.67 | 384.42 | 10.61 |
| **KSE-PDE ($128^2$)** | **37.91** | **174.38** | 15.37 | 2096.35 |
| KSE-BPF ($N_p = 128$) | 47.85 | 12.31 | 11.30 | 12.30 |
| **KSE-BPF ($N_p = 1024$)** | **36.80** | **74.93** | **9.44** | **80.86** |
| **KSE-EKF** | 42.05 | 4.07 | **9.33** | **2.60** |

Although the ITAE provides a summary measure of the state estimation accuracy, it does not assess the agreement between the underlying filtering distributions. We therefore analyse the similarity among the approximated solutions by computing the Hellinger distance ℋ between two probability distributions, say *F*_1_ and *F*_2_. With respect to the Lebesgue measure, let *f*_1_ and *f*_2_ denote their density functions. The square of the Hellinger distance is then defined as 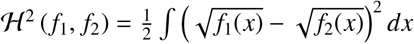.

Figures 7 and 8 show comparative results at a fixed time *t*. For the BPF method, we let *N*_*P*_ = 2.5 × 10^6^. The algorithms are provided in the Supplementary Material.

**Figure 7:**
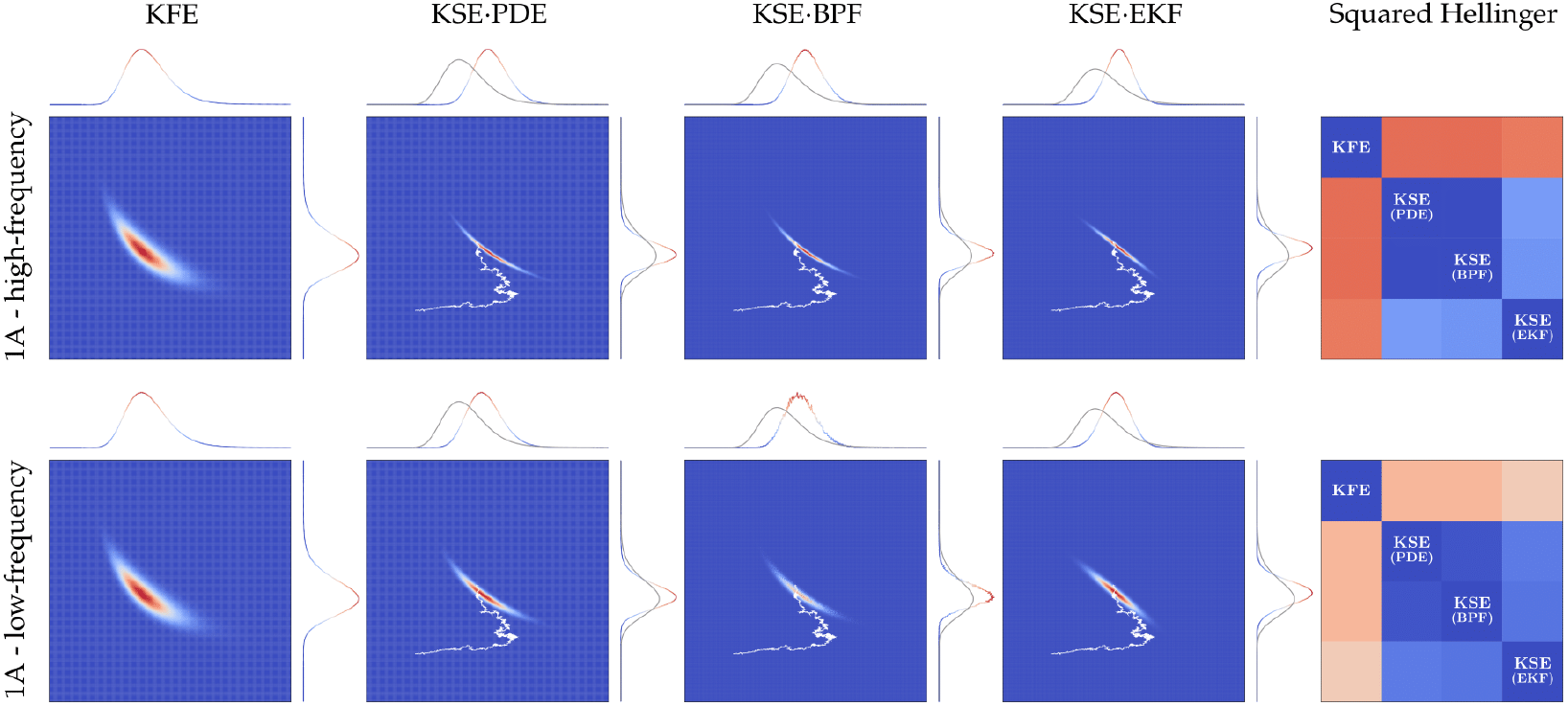
Case 1A - approximated solution of the KSE at *t* = 16h with the finite differences and splitting-up approximation for PDEs (KSE PDE), the BPF (KSE BPF) and the EKF (KSE EKF). The trajectory of *X* from which observations were simulated is shown in white and can also be found in Figure 3.

**Figure 8:**
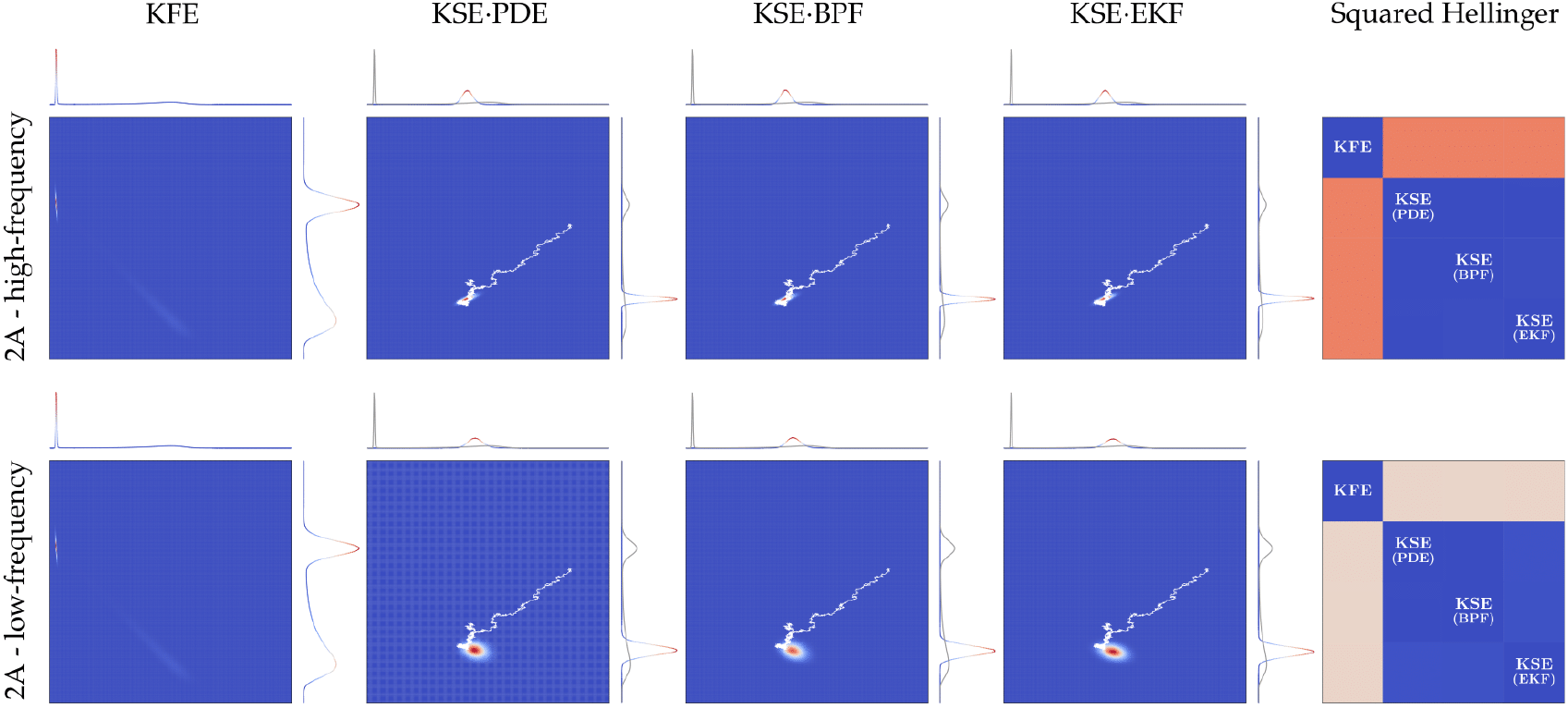
Case 2A - KSE PDE, KSE BPF, and KSE EKF approximations at *t* = 8h. Differently from the solution of the KFE, more probability mass moves towards the washout on account of the observations.

Figure 9 shows the square of the Hellinger distance for Case 1 between *π* approximated by the splitting-up method and by the BPF. It is clear that the discretisation error decreases with a finer grid refinement. The distance between *π* approximated by the EKF and by the BPF is also shown. The EKF fails to capture higher moments of the distribution, hence its divergence from the BPF result. This has already been suggested by Figure 7, for which we can observe non-zero skewness and excess kurtosis resulting from having both a non-linear dynamic process and a non-linear observation process. The shape of the conditional density for different times can be found in the Supplementary Material.

**Figure 9:**
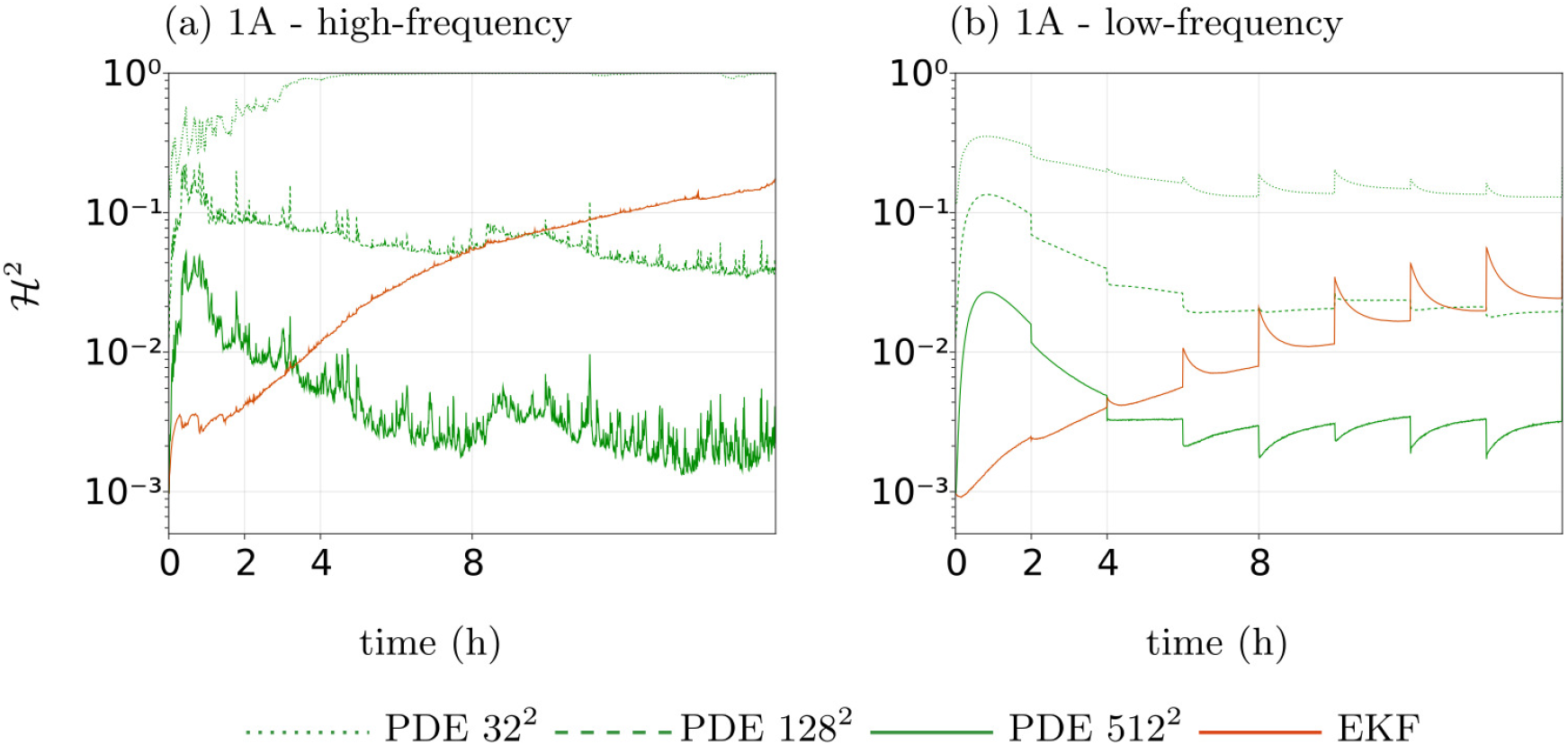
Case 1A - Squared Hellinger distances between *π* approximated by different grid refinements and the BPF (green), and between *π* approximated by the EKF and the BPF (orange). We consider three scenarios for the grid refinement, each with a total count of 32^2^, 128^2^, and 512^2^ finite volumes.

Figure 10 shows the square of the Hellinger distance between *π* approximated by the splittingup method and by the BPF, now for Case 2A. We again include the square of the Hellinger distance between *π* approximated by the EKF and by the BPF. Here, the EKF suffers in the highly non-linear region of the state space, namely when the state is near the saddle point, *t ≥* 4h.

**Figure 10:**
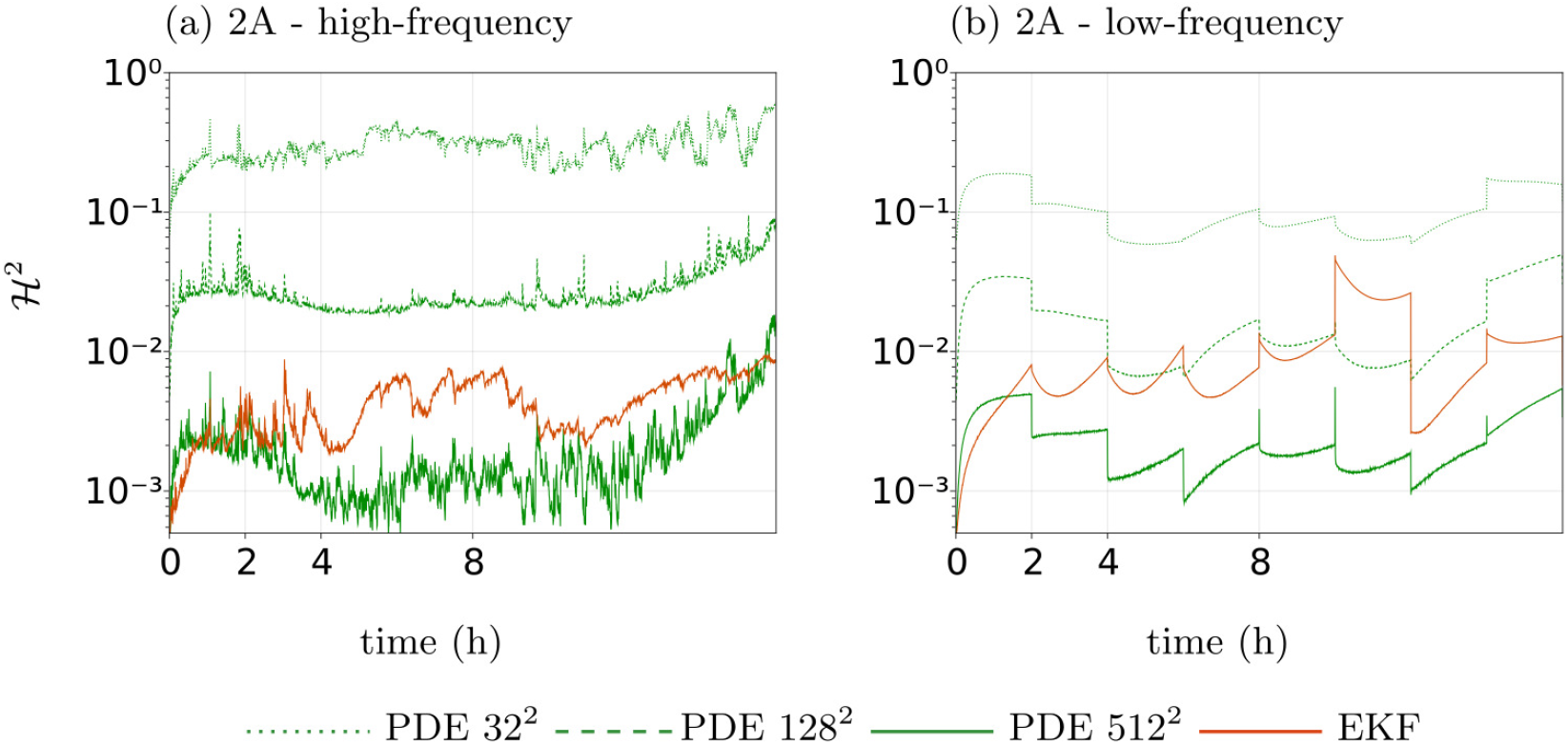
Case 2A - Squared Hellinger distances between *π* approximated by different grid refinements and the BPF (green), and between *π* approximated by the EKF and the BPF (orange). Same algorithmic settings as in Figure 9.

In Figure 10, the approximation with the splitting-up method combined with finite-differences on a 512^2^ grid mismatches the approximation with the BPF at observation times. At *t* = 8*h*, for instance, the Hellinger distance between both distributions increases. The densities approximated by grid-based schemes or by the EKF assume that the solution is uniformly distributed within each finite volume. The density approximated by the BPF results from counting the number of particles within each finite volume. Here, we encounter the problem of degenerate volumes with no particles. This issue is similar to the so-called sample impoverishment, which is a reduction in the number of distinct sample values when the model noise variance for process *X* is very low. As a result, the particles will hardly diversify and may collapse to a single point in state space after a few iterations. There are different solutions for this problem, the simplest being an increase in the number of particles. This, however, leads to an unreasonable computational load. Alternatively, a density kernel estimate can smooth the particle-based approximation and extrapolate the density values for finite volumes without any particles. Other solutions are more complex and often problem-specific. See [64] and [65] for more methods to prevent sample impoverishment.

## 4. Discussion and concluding remarks

In the first part of this work, we established the mathematical guarantees for the inverse problem of state-estimating a stochastic chemostat. Specifically, we proved the existence and uniqueness of the solution to the SDE governing the evolution of biomass and substrate concentrations, ensured trajectory positivity, and discussed the well-posedness of the KFE. Coupled with an observation model, we presented the filtering problem and established conditions for the well-posedness of the KSE. Consequently, the numerical pathwise approximations inherited a theoretical guarantee of reliability.

As a complement to the first part, our work advances existing methodologies to numerically realise the filtering solutions for stochastic chemostats using a PDE-based approach. We presented numerical solutions to the governing SDE, the corresponding KFE, and the pathwise KSE, with the latter two realised through a grid-based scheme. These PDE-based approximations impose no restrictions on the distribution, allowing for greater accuracy compared to linearisation methods. Furthermore, they allowed us to visualise the shape of the entire density across the state space, providing a more comprehensive understanding of system dynamics and associated uncertainty.

This dual focus on theoretical and numerical methodologies contributes to the management of microbial populations, particularly in industrial settings such as bioreactors. In these environments, precise control of microbial interactions and substrate use is essential for optimising productivity and maintaining system stability. Our results provide insights into the dynamics of microbial populations with a mathematical guarantee of reliability. This allows operators to accurately predict and adjust conditions to optimise yield, efficiency, or other desired operational targets.

## Appendix A. Existence and uniqueness of the solution to an SDE

### Condition A.

*For all t* ∈ [0, ∞), *v* ∈ *D*_*V*_, *and pairs z, z*^′^ ∈ *C*_*Z*_, *there exists a constant α >* 0 *independent of t, v, z and z*^′^, *such that*

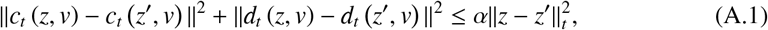

*and*

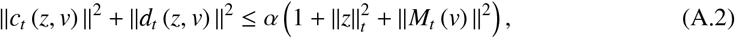

*in which* ∥*a*∥_*t*_ := sup_*s*∈[0,*t*]_ ∥*a*_*s*_∥, *for a* = *z* − *z*^′^ *(inequality A.1) and a* = *z (inequality A.2), and M*_*t*_ *is a* ℬ *D-measurable functional with* 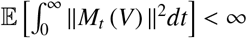.

Then, we have a theorem [33, Theorem 5.1.1], which we restate here without proof.

### Theorem A

***(Existence and pathwise uniqueness of strong solutions)***. *Suppose that c*_*t*_, *d*_*t*_, (Ω, ℱ, (ℱ_*t*_, *≥ t*, ℙ), *V, B*^*Z*^, *and z*_0_ *are given, with c*_*t*_ *and d*_*t*_ *non-anticipative. If Condition A holds, SDE* (2.3) *has a pathwise unique strong solution*.

## Appendix B. Additional conditions for the KSE

1. *K* is invertible;
2. Condition A holds for SDE (2.23), such that this SDE has a unique strong solution *X* = (*X*_*t*_, *t* ≥ 0) by Theorem A;
3. Condition A also holds for SDE (2.24) when *M*_*t*_(*X*) = *h*(*X*_*t*_).

## Data availability statement

The code used in this work is at BayesianFilteringLab (DOI: 10.5281/zenodo.21377771).

