## Supplementary Material for "Pathwise approximations to solving the filtering problem for the stochastic chemostat"

2.3.2 Solving the Kushner-Stratonovich equation with sequential Monte Carlo . 13
2.3.3 Solving the Kushner-Stratonovich equation with linearisation methods . . 15

**3 Chemostat equations** **20**

**4 Numerical experiments** **22**

### 1. Notation

Table 1: Notation table for the main text.

| Symbol | Definition / Usage |
|---|---|
| $(b_e, s_e)$ | Equilibrium pair |
| $(b_t, t \geq 0)$ | Biomass concentration process |
| $(s_t, t \geq 0)$ | Substrate concentration process |
| $(\omega_X, \omega_Y)$ | Pair composing $\omega \in \Omega$ |
| $(\mathbb{R}^2)^*_+$ | Positive quadrant |
| $A_X$ | Element of $\mathcal{B}(\mathbb{S}_X)$ |
| $A_Z$ | Element of $\mathcal{B}(\mathbb{S}_Z)$ |
| $A_{Z,V}$ | Element of $\mathcal{B}(\mathbb{R})$ |
| $B = (B_t, t \geq 0)$ | Brownian motion process |
| $B^X$, $B^Y$ | Brownian motion processes for $X$ and $Y$ |
| $B^Z$ | Brownian motion process for $Z$ |
| $C_X, C_Y$ | Space of continuous function w/ values in $\mathbb{S}_X$ and $\mathbb{S}_Y$, respectively |
| $C_Z$ | Space of continuous functions w/ values in $\mathbb{S}_Z$ |
| $D$ | Dilution rate |
| $D_V$ | Space of right-continuous functions w/ values in $\mathbb{S}_V$ w/ LH limits |
| $D_t$ | Diffusion tensor |
| $G$ | Diffusion tensor associated to $X$ |
| $I_{N_B \times N_B}$ | Covariance of standard $N_B$-dimensional $B$ |
| $J = (J^1, \ldots, J^{n_Z}, \ldots J^{N_Z})$ | Probability current |
| $K$ | Matrix of size $N_Y \times N_Y$ |
| $K_i$ | Inhibitor parameter |
| $K_s$ | Half-saturation parameter |
| $L = (L_t, t \geq 0)$ | Radon-Nikodym derivative process |
| $N_B$ | Dimension of $B$ |
| $N_P$ | Number of Monte Carlo particles |
| $N_X$, $N_Y$ | Dimensions of $X$ and $Y$ |
| $N_Z$ | Dimension of $Z$ |
| $N_{B^X}$, $N_{B^Y}$ | Dimensions of $B^X$ and $B^Y$ |
| $N_{B^Z}$ | Dimension of $B^Z$ |
| $N_t$ | Number of time indices |
| $Q$ | Compact form for Lyapunov bound |

| | |
|---|---|
| $S, \partial S$ | Open set of $\mathbb{S}_Z$ and its boundary |
| $S_r$ | Indexed set for set sequence |
| $T$ | Fixed time instant greater than 0 |
| $T_\beta$ | Constant |
| $V = (V_t, t \geq 0)$ | Auxiliary stochastic process |
| $X = (X_t, t \geq 0)$, $Y = (Y_t, t \geq 0)$ | State and observation processes |
| $X_0$ | Initial condition for $X$ |
| $Z = (Z_t, t \geq 0)$ | General stochastic process |
| $X^{(n_P)}$ | $n_P$th stochastic process $X$ |
| $a$ | Auxiliary variable for norms |
| $a_t$ | Functional $a_t : C_Z \times D_V \to \mathbb{R}$ |
| $\alpha, \beta$ | Constants |
| $\alpha_r, \beta_r, \gamma_r$ | Constants |
| $b^{in} = \left(b^{in}_t, t \geq 0\right)$ | Incoming biomass concentration process |
| $c_t$ | Functional for the dynamics |
| $d_t$ | Functional $d_t : D_V \times C_Z \to \mathbb{R}^{N_Z \times N_{B^Z}}$ |
| $f, g, h$ | Autonomous versions of the SDE coefficients |
| $f_t, g_t$ | SDE functionals replacing $c_t$ and $d_t$, respectively |
| $h_t, k_t$ | SDE functionals replacing $c_t$ and $d_t$, respectively |
| $l$ | Lyapunov function |
| $n_X, n_Y, n_{B^X}, n_{B^Y}, n'_X, \ldots$ | Iterators |
| $n_Z$ | Iterator from 1 to $N_Z$ |
| $n_{B^Z}, n'_{B^Z}$ | Iterators from 1 to $N_{B^Z}$ |
| $p\,(t, \cdot)$ | Probability density function of $\mathbb{P}(0, z_0; t, \cdot)$ |
| $p^{\Delta t}_{n_t}$ | Approximated posterior density at time $t_{n_t}$ |
| $p_s$ | Measurable non-negative functions on $\mathbb{S}_Z$ |
| $r$ | Integer for indexing sequences |
| $r'$ | Integer pair to $r$ |
| $s^{in} = \left(s^{in}_t, t \geq 0\right)$ | Incoming substrate concentration process |
| $t, s, s'$ | Time variables in Section 2 |
| $t_e$ | Exit time |
| $t_\infty$ | Limit of indexed stopping time $t_r$ |
| $t_r$ | Indexed time for stopping time sequence |
| $u = (u_t, t \geq 0)$ | State-feedback control process |
| $v = (v_t, t \geq 0)$ | Auxiliary process |
| $z = (z_t, t \geq 0)$ | General process |

| | |
|---|---|
| $z_0$ | Initial condition for $z$ |
| $z' = (z'_t, t \geq 0)$ | Pair to $z$ |
| $y_t$ | Observation at time $t$ |
| $\Lambda = (\Lambda_t, t \geq 0)$ | Innovation process |

### 2. Stochastic filtering

#### 2.1. Preliminaries

The standard Brownian motion process $B$ is defined as follows:

**Definition 2.1 (Standard Brownian motion process).** *A stochastic process $B = (B_t, t \geq 0)$ on $(\Omega, \mathcal{F}, \mathbb{P})$ is said to be an $N_B$-dimensional standard Brownian motion process if the following conditions are satisfied:*

i. *$\mathbb{P}$-almost surely $B_0 = 0$; that is $\mathbb{P}(B_0 = 0) = 1$;*
ii. *$B$ has independent increments; that is, $B_0, B_{t_1} - B_0, \ldots, B_{t_{N_t}} - B_{t_{N_t-1}}$ are independent, for every integer $N_t > 1$ and indices $0 < t_1 < \cdots < t_{N_t} < \infty$;*
iii. *$B$ has increments which are normally distributed with mean $0$ and variance $(t-s) I_{N_B \times N_B}$, for all $s \in [0, t)$, where $I_{N_B \times N_B}$ is the identity matrix of size $N_B$; that is, $B_t - B_s \sim \mathcal{N}(0, (t-s) I_{N_B \times N_B})$;*
iv. *$B(\omega) = (B_t(\omega), t \in [0, \infty))$ is continuous for $\mathbb{P}$-almost every $\omega \in \Omega$, where $B(\omega)$ is the sample path, or trajectory, of $B$ associated with $\omega$.*

A stochastic process which only satisfies conditions (1)-(3) is called a standard pre-Brownian motion process. It can be shown that such process admits a version, a modification, $\tilde{B}$ whose sample paths are $\mathbb{P}$-almost surely locally Hölder-continuous for every exponent $\alpha \in (0, \frac{1}{2})$; that is, for all finite times $T > 0$, there exists a constant $\beta > 0$ such that

$$\mathbb{P}\left[\omega \in \Omega : \sup_{0 \leq s < t < T} \frac{\|\widetilde{B}_t(\omega) - \widetilde{B}_s(\omega)\|}{(t-s)^{\alpha}} \leq \beta\right] = 1, \alpha \in \left(0, \frac{1}{2}\right), \|\cdot\| = \text{Euclidean norm.}$$

In this work, we choose such version to work with by setting $B = \tilde{B}$. If $B$ has locally Hölder-continuous sample paths, it satisfies condition (4), thereby satisfying all the conditions to classify $B$ as the standard Brownian motion process. Proof of the existence of such $B$ can be found in, for instance, [1, Section 2].

Alongside the aforementioned definition, we provide two examples of processes driven by $B$. These examples introduce the SDE forms used in the main manuscript.

**Example 2.1 (SDE with additive noise).** *Consider a $\mathbb{R}$-valued process $Z$, a univariate standard Brownian motion process $B^Z$, and non-anticipative coefficients $c_t$ and $d_t(Z, V) = \varepsilon$ with $\varepsilon \in \mathbb{R}$. Let the following SDE relate, additively, $B^Z$ and $Z$:*

$$dZ_t = c_t(Z, V)\, dt + \varepsilon dB_t^Z, \quad Z_0 = z_0.$$

*For $c_t \equiv 0$, Condition A holds: the resulting SDE possesses a pathwise unique strong solution by Theorem 2.1, and such solution takes the common form $Z_t = z_0 + \varepsilon B_t^Z$.*

The example below illustrates the notion of positivity of the solution of an SDE.

**Example 2.2 (SDE with multiplicative noise).** *Consider a process $Z$, a univariate standard Brownian motion process $B^Z$, and non-anticipative $c_t \equiv 0$ and $d_t(Z, V) = \varepsilon Z_t$ with $\varepsilon \in \mathbb{R}$. Let the following SDE relate, multiplicatively, $B^Z$ and $Z$:*

$$dZ_t = \varepsilon Z_t dB_t^Z, \quad Z_0 = z_0. \tag{1}$$

*It can be shown that a pathwise unique strong solution to SDE* (1) *exists, and that for any positive-valued random variable* $z_0$*, a sample path* $Z(\omega) = (Z_t(\omega), t \geq 0)$ *has positive real values for* $\mathbb{P}$*-almost every* $\omega \in \Omega$.

*Itô's formula [2, Theorem 6] can be used on functions from* $\mathbb{S}_Z$ *with continuous first and second derivatives. For the logarithm function, SDE* (1) *yields*

$$d(\ln(Z_t)) = \varepsilon dB_t^Z - 1/2\varepsilon^2 t, \quad \ln(Z_0) = \ln(z_0).$$

*Therefore,* $Z_t = z_0 \exp\left(\varepsilon B_t^Z - 1/2\varepsilon^2 t\right)$*, assuring that* $Z$ *is constrained to* $S \subseteq \mathbb{R}_+$.

We conclude this section by defining non-anticipative functionals. For $\mathcal{B}_s(C_Z)$ - respectively, $\mathcal{B}_s(D_V)$ - denoting the smallest $\sigma$-field which makes $(Z_{s'}, s' \leq s)$ - respectively, $(V_{s'}, s' \leq s)$ - measurable and for $\mathcal{B}_{t^+} = \cap_{s>t}\mathcal{B}_s$ denoting the smallest $\sigma$-field containing $\mathcal{B}_s$ for all $s > t$, we define non-anticipativeness as follows:

**Definition 2.2 (Non-anticipative functionals).** *A* $\mathcal{B}(C_Z) \times \mathcal{B}(D_V)$*-measurable functional* $a_t : C_Z \times D_V \to \mathbb{R}$ *is said to be non-anticipative if* $a_t$ *is* $\mathcal{B}_{t^+}(C_Z) \times \mathcal{B}_{t^+}(D_V)$*-measurable. That is,* $a_t^{-1}(A_{Z,V}) \in \mathcal{B}_{t^+}(C_Z) \times \mathcal{B}_{t^+}(D_V)$ *for every* $A_{Z,V} \in \mathcal{B}(\mathbb{R})$.

If each component functional $c_t^{n_Z}$ is non-anticipative in the sense of Definition 2.2, then the functional $c_t = \left(c_t^1, \ldots, c_t^{n_Z}, \ldots, c_t^{N_Z}\right)$ is said to be non-anticipative. Similarly, if each entry $d_t^{n_Z, n_{B^Z}}$ is non-anticipative, then the matrix-valued functional $d_t = \left(d_t^{n_Z, n_{B^Z}}\right)$, with $n_Z = 1, \ldots, N_Z$ and $n_{B^Z} = 1, \ldots, N_{B^Z}$, is said to be non-anticipative.

#### 2.2. Proofs

Let us consider two probability measures $\mathbb{P}$ and $\mathbb{Q}$ on $\mathcal{F}$. We recall that $\mathbb{P}$ is called *absolutely continuous* with respect to $\mathbb{Q}$, written as $\mathbb{P} \ll \mathbb{Q}$, if every null-set of $\mathbb{Q}$ is a null-set of $\mathbb{P}$. We restrict the definition of the measure $\mathbb{P}$ to $\mathcal{F}_t$, written as $\mathbb{P}_t$; that is, $\mathbb{P}$ is restricted to all the events that can be described in terms of the behaviour of $(X_s, s \leq t)$ and $(Y_s, s \leq t)$. Let us assume the existence of a $\mathbb{Q}$ such that $\mathbb{P}_t \ll \mathbb{Q}_t$ for all finite $t$. Then the Radon–Nikodym theorem [3] guarantees the existence of a random variable $L_t$, unique for $\mathbb{Q}$-almost every $\omega \in \Omega$, satisfying

$$\mathbb{E}_{\mathbb{P}}[\phi(X_t)] = \mathbb{E}_{\mathbb{Q}}[\phi(X_t) L_t], \tag{2}$$

where $L_t := \frac{d\mathbb{P}_t}{d\mathbb{Q}_t}$ is called the Radon–Nikodym derivative of $\mathbb{P}_t$ with respect to $\mathbb{Q}_t$. sNotably, $\mathbb{E}_{\mathbb{Q}}[L_t] = 1$ as a direct consequence of the definition.

The Radon–Nikodym derivative $L_t$ plays the role of a likelihood ratio between the probability measures $\mathbb{P}_t$ and $\mathbb{Q}_t$. It allows the conditional expectation of $\phi$, i.e. the filter $\pi_t(\phi)$, to be expressed in terms of expectations under the reference measure $\mathbb{Q}$:

$$\begin{aligned} \pi_t(\phi) &= \frac{\mathbb{E}_{\mathbb{Q}}\left[\phi(X_t)\ L_t \mid \mathcal{F}_t^Y\right]}{\mathbb{E}_{\mathbb{Q}}\left[L_t \mid \mathcal{F}_t^Y\right]} \\ &=: \frac{\rho_t(\phi)}{\rho_t(\mathbb{1})}, \end{aligned} \tag{3}$$

Eq. (3) is known as the Bayes formula for stochastic processes or *Kallianpur-Striebel formula* [4, 5]. A concise modern derivation is given in Proposition 19 of [6]. The denominator $\rho_t(\mathbb{1})$ acts as a normalising factor that converts the unnormalised conditional expectation in the numerator into a probability measure. The hope is that $L_t$ is simple enough to make $\mathbb{E}_{\mathbb{P}}\left[\phi(X_t)|\mathcal{F}_t^Y\right]$ more tractable to compute via the right-hand side of Eq. (3) than via an integration over $\mathbb{P}$. For instance, some simplification might be achieved by switching from a model with restriction $\mathbb{P}_t$ in which $X$ and $Y$ are coupled to a model with restriction $\mathbb{Q}_t$ in which they are independent, while preserving the distribution of $X_t$. For our problem, we will assume the existence of a $\mathbb{Q}$ under which $(Y_s, s \leq t)$ is a standard Brownian motion process independent of $(X_s, s \leq t)$ for all finite $t$.

The following result will be useful for proving Proposition 2.1.

**Lemma 2.1 (SDE for the Radon-Nikodym derivative).** *The derivative $L_t$ can be written as*

$$L_t = \exp\left[\int_0^t (K^{-1}h(X_s))^\top K^{-1}dY_s - \frac{1}{2}\int_0^t \left\|K^{-1}h(X_s)\right\|^2 ds\right],$$

*and it evolves according to the following SDE:*

$$dL_t = L_t\tilde{h}^\top(X_t)\,d\tilde{Y}_t, \tag{4}$$

*where $\tilde{Y} = (\tilde{Y}_t, t \geq 0)$ is the rescaled observation process (i.e. $\tilde{Y}_t = K^{-1}Y_t, t \geq 0$), and $\tilde{h}$ is the rescaled observation function $\tilde{h}(X_t) = K^{-1}h(X_t)$.*

**Proof 2.1.** *Let $q(t, dY_t)$ be the density of $\mathcal{N}(dY_t; 0, KK^\top dt)$. $L_t$ can be written as*

$$\begin{aligned}
L_t &= \frac{p(t, dY_{0:t}|X_{0:t})}{q(t, dY_{0:t})} \\
&= \frac{\prod_{s=0}^t p(t, dY_s|X_s)}{\prod_{s=0}^t q(t, dY_s)} \\
&= \prod_{s=0}^t \frac{\text{density of } \mathcal{N}(dY_s; h(X_s)\,ds, KK^\top ds)}{\text{density of } \mathcal{N}(dY_s; 0, KK^\top ds)} \\
&= \prod_{s=0}^t \exp\left[(K^{-1}h(X_s))^\top K^{-1}dY_s - \frac{1}{2}\left\|K^{-1}h(X_s)\right\|^2 ds\right] \\
&\overset{\lim_{dt\to 0}}{=} \exp\left[\int_0^t (K^{-1}h(X_s))^\top K^{-1}dY_s - \frac{1}{2}\int_0^t \left\|K^{-1}h(X_s)\right\|^2 ds\right],
\end{aligned}$$

*where in the last step we took the continuum limit $\lim_{dt\to 0}$.*

*If we define*

$$\begin{aligned}
\Lambda_t &:= \log L_t = \left[\int_0^t (K^{-1}h(X_s))^\top K^{-1}dY_s - \frac{1}{2}\int_0^t \left\|K^{-1}h(X_s)\right\|^2 ds\right], \\
d\Lambda_t &= \left[(K^{-1}h(X_t))^\top K^{-1}dY_t - \frac{1}{2}\left\|K^{-1}h(X_t)\right\|^2 dt\right].
\end{aligned}$$

*Then $dL_t = d(\exp \Lambda_t)$ and Itô's formula applied to the process* $\exp \Lambda$ *leads to*

$$\begin{aligned} dL_t &= d(\exp \Lambda_t) \\ &= \exp \Lambda_t d\Lambda_t + \frac{1}{2}\exp \Lambda_t (d\Lambda_t)^2 \\ &= \exp \Lambda_t d\Lambda_t + \frac{1}{2}\exp \Lambda_t \left\|K^{-1}h(X_t)\right\|^2 dt \\ &= L_t\left[(K^{-1}h(X_t))^\top K^{-1}dY_t - \frac{1}{2}\left\|K^{-1}h(X_t)\right\|^2 dt\right] + \frac{1}{2}L_t\left\|K^{-1}h(X_t)\right\|^2 dt \\ &= L_t\tilde{h}^\top(X_t)\, d\tilde{Y}_t. \end{aligned}$$

**Proposition 1 (The KSE in Proposition 2.1 (revisited)).** *Let $\phi$ have continuous partial derivatives of every order up to 2, all of which are bounded. Then we can write*

$$\pi_t(\phi) = \pi_0(\phi) + \int_0^t \pi_s(\mathcal{A}^*\phi)ds + \int_0^t \left(\pi_s(\phi\tilde{h}) - \pi_s(\phi)\pi_s(\tilde{h})\right)^\top \left(d\tilde{Y}_s - \pi_s(\tilde{h})\,ds\right), \tag{5}$$

*where $\pi_0(\phi) = \mathbb{E}_{\mathbb{P}}[\phi(X_0)] = \mathbb{E}_{\mathbb{Q}}\left[\phi(X_0)L_0|\mathcal{F}_0^Y\right]$.*

*Moreover, an application of integration by parts to Eq. (5) results in the so-called Kushner-Stratonovich equation (KSE)*

$$dp_t(x) = \mathcal{A}^* p_t(x)\,dt + p_t(x)\left(\tilde{h}(x) - \pi_t(\tilde{h})\right)^\top \left(d\tilde{Y}_t - \pi_t(\tilde{h})\,dt\right).$$

**Proof 2.2.** *Our problem is to compute the expectation under the posterior distribution $\pi_t$ of some function $\phi$. This is a conditional expectation, rewritten in terms of a reference probability measure $\mathbb{Q}$ as*

$$\mathbb{E}_{\mathbb{P}}\left[\phi(X_t)|\mathcal{F}_t^Y\right] = \frac{\mathbb{E}_{\mathbb{Q}}\left[\phi(X_t)\,L_t|\mathcal{F}_t^Y\right]}{\mathbb{E}_{\mathbb{Q}}\left[L_t|\mathcal{F}_t^Y\right]} =: \frac{\rho_t(\phi)}{\rho_t(\mathbb{1})}, \tag{6}$$

*where we have defined the unnormalised estimate $\rho_t(\phi) := \mathbb{E}_{\mathbb{Q}}\left[\phi(X_t)\,L_t|\mathcal{F}_t^Y\right]$.*

*Notably, $d\mathbb{E}_{\mathbb{P}}\left[\phi(X_t)|\mathcal{F}_t^Y\right] = d\left(\frac{\rho_t(\phi)}{\rho_t(\mathbb{1})}\right)$. By using Itô's formula*[1]*, we obtain*

$$\begin{aligned} d\mathbb{E}_{\mathbb{P}}\left[\phi(X_t)|\mathcal{F}_t^Y\right] &= \frac{1}{\rho_t(\mathbb{1})}d\rho_t(\phi) + \rho_t(\phi)d\left(\frac{1}{\rho_t(\mathbb{1})}\right) + d\rho_t(\phi)d\left(\frac{1}{\rho_t(\mathbb{1})}\right) \\ &= \frac{1}{\rho_t(\mathbb{1})}\mathbb{E}_{\mathbb{Q}}\left[d(\phi(X_t)L_t)|\mathcal{F}_t^Y\right] + \mathbb{E}_{\mathbb{Q}}\left[\phi(X_t)L_t|\mathcal{F}_t^Y\right]d\left(\frac{1}{\rho_t(\mathbb{1})}\right) \\ &\quad + \mathbb{E}_{\mathbb{Q}}\left[d(\phi(X_t)L_t)|\mathcal{F}_t^Y\right]d\left(\frac{1}{\rho_t(\mathbb{1})}\right), \end{aligned} \tag{7}$$

*where the second equality results from taking the stochastic differential inside the expectation under $\mathbb{Q}$ (see [7, Section 7, Lemma 7.2.7]).*

[1]Recall that we have to consider the product of differentials, since the Itô's formula corresponds to a Taylor expansion up to second order for diffusion processes.

*Let us evaluate these terms separately:*

*The first term can be expressed as*

$$\begin{aligned}\frac{1}{\rho_t(\mathbb{1})}\mathbb{E}_{\mathbb{Q}}\left[d(\phi(X_t)\ L_t)|\mathcal{F}_t^Y\right] &= \frac{1}{\rho_t(\mathbb{1})}\mathbb{E}_{\mathbb{Q}}\left[L_t d\phi_t + \phi_t dL_t + \underbrace{d\phi_t dL_t}_{=0}\right] \\ &= \frac{1}{\rho_t(\mathbb{1})}\left(\mathbb{E}_{\mathbb{Q}}\left[L_t\mathcal{A}^*\phi_t(X_t)|\mathcal{F}_t^Y\right]dt + \mathbb{E}_{\mathbb{Q}}\left[\phi_t(X_t)L_t\tilde{h}^\top(X_t)|\mathcal{F}_t^Y\right]d\tilde{Y}_t\right) \\ &= \mathbb{E}_{\mathbb{P}}\left[\mathcal{A}^*\phi_t(X_t)|\mathcal{F}_t^Y\right]dt + \mathbb{E}_{\mathbb{P}}\left[\phi_t(X_t)\tilde{h}^\top(X_t)|\mathcal{F}_t^Y\right]d\tilde{Y}_t,\end{aligned}$$

*where we used the Itô formula for the products. The term $d\phi_t(X_t)\,dL_t$ equals zero because the noise components are independent.*

*For the second term, we first need to obtain $d\left(\frac{1}{\rho_t(\mathbb{1})}\right)$. For this, we apply Itô formula to the inverse of the process $\rho(1) = (\rho_t(\mathbb{1}), t \geq 0)$, where $L_t$ solves SDE (4) from Lemma 2.1.*

$$\begin{aligned}d\rho(1) &= d\left(\mathbb{E}_{\mathbb{Q}}\left[L_t|\mathcal{F}_t^Y\right]\right) \\ &= \mathbb{E}_{\mathbb{Q}}\left[L_t\tilde{h}^\top(X_t)\,d\tilde{Y}_t|\mathcal{F}_t^Y\right].\end{aligned}$$

*Using Itô formula, we obtain:*

$$\begin{aligned}d\left(\frac{1}{\rho_t(\mathbb{1})}\right) &= -\rho_t(\mathbb{1})^{-2}\,d\rho_t(\mathbb{1}) + \rho_t(\mathbb{1})^{-3}\,(d\rho_t(\mathbb{1}))^2 \\ &= -\rho_t(\mathbb{1})^{-2}\,\mathbb{E}_{\mathbb{Q}}\left[L_t\tilde{h}^\top(X_t)|\mathcal{F}_t^Y\right]d\tilde{Y}_t + \rho_t(\mathbb{1})^{-3}\left(\mathbb{E}_{\mathbb{Q}}\left[L_t\left\|\tilde{h}(X_t)\right\||\mathcal{F}_t^Y\right]\right)^2 dt \\ &= -\rho_t(\mathbb{1})^{-1}\,\mathbb{E}_{\mathbb{P}}\left[(\tilde{h}^\top(X_t)|\mathcal{F}_t^Y\right]d\tilde{Y}_t + \rho_t(\mathbb{1})^{-1}\left(\mathbb{E}_{\mathbb{P}}\left[\left\|\tilde{h}(X_t)\right\||\mathcal{F}_t^Y\right]\right)^2 dt \\ &= -\rho_t(\mathbb{1})^{-1}\left(\mathbb{E}_{\mathbb{P}}\left[\tilde{h}(X_t)|\mathcal{F}_t^Y\right]\right)^\top\left(d\tilde{Y} - \mathbb{E}_{\mathbb{P}}\left[\tilde{h}(X_t)|\mathcal{F}_t^Y\right]dt\right).\end{aligned} \tag{8}$$

*Thus, the second term in Eq. (7) reads:*

$$\mathbb{E}_{\mathbb{Q}}\left[\phi(X_t)L_t|\mathcal{F}_t^Y\right]d\left(\frac{1}{\rho_t(\mathbb{1})}\right) = -\mathbb{E}_{\mathbb{P}}\left[\phi(X_t)|\mathcal{F}_t^Y\right]\left(\mathbb{E}_{\mathbb{P}}\left[\tilde{h}(X_t)|\mathcal{F}_t^Y\right]\right)^\top\left(d\tilde{Y} - \mathbb{E}_{\mathbb{P}}\left[\tilde{h}(X_t)|\mathcal{F}_t^Y\right]dt\right).$$

*Finally, the third term uses the result in Eq. (8) together with the following unnormalised posterior expectation:*

$$\begin{aligned}\mathbb{E}_{\mathbb{Q}}\left[d(\phi(X_t)\ L_t)|\mathcal{F}_t^Y\right] &= \mathbb{E}_{\mathbb{Q}}\left[d(\phi(X_t))L_t + \phi(X_t)\,d(L_t) + d(\phi(X_t))d(L_t)\,|\mathcal{F}_t^Y\right] \\ &= \mathbb{E}_{\mathbb{Q}}\left[L_t\mathcal{A}^*\phi(X_t)|\mathcal{F}_t^Y\right]dt + \mathbb{E}_{\mathbb{Q}}\left[\phi(X_t)L_t\tilde{h}^\top(X_t)\,d\tilde{Y}_t|\mathcal{F}_t^Y\right].\end{aligned}$$

*Here, we again used $\mathbb{E}_{\mathbb{Q}}\left[d(\phi_t(X_t))d(L_t)|\mathcal{F}_t^Y\right] = 0$ because the noise components are independent.*

*Keeping only terms up to $O(dt)$, the third term in Eq. (7) reads:*

$$\mathbb{E}_{\mathbb{Q}}\left[d(\phi(X_t)\ L_t)|\mathcal{F}_t^Y\right]d\left(\frac{1}{\rho_t(\mathbb{1})}\right) = -\mathbb{E}_{\mathbb{P}}\left[\phi_t(X_t)\tilde{h}^\top(X_t)|\mathcal{F}_t^Y\right]\mathbb{E}_{\mathbb{P}}\left[\tilde{h}(X_t)|\mathcal{F}_t^Y\right]dt.$$

*Adding up and rearranging the terms, we obtain:*

$$\begin{aligned}
d\mathbb{E}_{\mathbb{P}}\left[\phi\left(X_t\right)|\mathcal{F}_t^Y\right] &= \mathbb{E}_{\mathbb{P}}\left[\mathcal{A}^*\phi_t\left(X_t\right)|\mathcal{F}_t^Y\right]dt + \mathbb{E}_{\mathbb{P}}\left[\phi_t\left(X_t\right)\tilde{h}^\top\left(X_t\right)|\mathcal{F}_t^Y\right]d\tilde{Y}_t \\
&\quad -\mathbb{E}_{\mathbb{P}}\left[\phi\left(X_t\right)|\mathcal{F}_t^Y\right]\left(\mathbb{E}_{\mathbb{P}}\left[\tilde{h}\left(X_t\right)|\mathcal{F}_t^Y\right]\right)^\top\left(d\tilde{Y} - \mathbb{E}_{\mathbb{P}}\left[\tilde{h}\left(X_t\right)|\mathcal{F}_t^Y\right]dt\right) \\
&\quad -\mathbb{E}_{\mathbb{P}}\left[\phi_t\left(X_t\right)\tilde{h}^\top\left(X_t\right)|\mathcal{F}_t^Y\right]\mathbb{E}_{\mathbb{P}}\left[\tilde{h}\left(X_t\right)|\mathcal{F}_t^Y\right]dt \\
&= \mathbb{E}_{\mathbb{P}}\left[\mathcal{A}^*\phi_t\left(X_t\right)|\mathcal{F}_t^Y\right]dt + \\
&\quad \left(\mathbb{E}_{\mathbb{P}}\left[\phi\left(X_t\right)\tilde{h}^\top\left(X_t\right) - \mathbb{E}_{\mathbb{P}}\left[\phi\left(X_t\right)|\,\mathcal{F}_t^Y\right]\mathbb{E}_{\mathbb{P}}[\tilde{h}^\top\left(X_t\right)|\mathcal{F}_t^Y]\middle|\mathcal{F}_t^Y\right]\right)\times \\
&\quad \left(d\tilde{Y} - \mathbb{E}_{\mathbb{P}}\left[\tilde{h}\left(X_t\right)|\mathcal{F}_t^Y\right]dt\right). \qquad (9)
\end{aligned}$$

*Notably, Eq. (9) can be written in the following integrated form:*

$$\pi_t(\phi) = \pi_0(\phi) + \int_0^t \pi_s(\mathcal{A}^*\phi)\,ds + \int_0^t \left(\pi_s(\tilde{h}\phi) - \pi_s(\phi)\pi_s(\tilde{h})\right)^\top (d\tilde{Y}_s - \pi_s(\tilde{h})\,ds),$$

*where* $\pi_0(\phi) = \mathbb{E}_{\mathbb{P}}\left[\phi(X_0)\,L_0|\mathcal{F}_0^Y\right] = \mathbb{E}_{\mathbb{P}}\left[\phi(X_0)\right]$.

#### 2.3. Algorithms

- Algorithm 1: Splitting algorithm for the KSE with methods for PDEs
- Algorithm 2: Continuous-Discrete Bootstrap Particle filter
- Algorithm 3: Continuous-Discrete Kalman-Bucy filter
- Algorithm 4: Squared-root Extended Kalman filter

##### 2.3.1. Solving the Kushner-Stratonovich equation with methods for PDEs

Consider a function $p_t : \mathbb{S}_X \to \mathbb{R}_+$. Assume $p_t$ to be twice differentiable and have evolution law

$$dp(t,x) = \mathcal{A}^* p(t,x)\,dt, \quad p(0,x) = p_0(x) \tag{10}$$

where the initial condition $p_0(x)$ is given, and the operator $\mathcal{A}^*$ is

$$\mathcal{A}^* p(t,x) = -\sum_{n_X=1}^{N_X} \frac{\partial}{\partial x_{n_X}} \left[ f^{n_X}(x)\, p(t,x) \right] + \frac{1}{2} \sum_{n_X=1}^{N_X} \sum_{n'_X=1}^{N_X} \frac{\partial^2}{\partial x_{n_X} \partial x_{n'_X}} \left[ G^{n_X,n'_X}(x)\, p(t,x) \right].$$

We approximate the solution to equations in the form of Eq. (10) by using a finite difference scheme on a given $N_X$-dimensional regular grid $\Omega^j$ with mesh $j = (j_1, \ldots, j_{n_X}, \ldots, j_{N_X})$ in order to approximate the differential operator $\mathcal{A}^*$. Let $\epsilon_{n_X}$ be a unit vector in the $n_X$th coordinate. The scheme approximates first-order derivatives of functions $\varphi$ on $\mathbb{S}_X$ evaluated at $x$ according to the following up-wind scheme:

$$\left.\frac{\partial \varphi}{\partial x_{n_X}}\right|_x \approx \frac{\varphi(x + \epsilon_{n_X} j_{n_X}) - \varphi(x)}{j_{n_X}}, \quad \text{if } f_{n_X} \geq 0, \text{ i.e. forward difference}$$

$$\left.\frac{\partial \varphi}{\partial x_{n_X}}\right|_x \approx \frac{\varphi(x) - \varphi(x - \epsilon_{n_X} j_{n_X})}{j_{n_X}}, \quad \text{if } f_{n_X} < 0, \text{ i.e. backward difference,}$$

and the second-order derivates as

$$\left.\frac{\partial^2 \varphi}{\partial x_{n_X}^2}\right|_x \simeq \frac{\varphi(x + \epsilon_{n_X} j_{n_X}) - 2\varphi(x) + \varphi(x - \epsilon_{n_X} j_{n_X})}{j_{n_X}^2}.$$

The cross term $\left.\frac{\partial^2 \varphi}{\partial x_{n_X} \partial x_{n'_X}}\right|_x$ is zero-valued for the dynamics discussed in this work. For general cases, see [8, Section 8].

Let $0 = t_0 < t_1 < \cdots < t_{n_t} \ldots$ be a uniform partition of the interval $[0, \infty)$ with time step $\Delta t = t_{n_t} - t_{n_t-1}$. If the set of observations $\{y_{n'_t}, 1 \leq n'_t \leq n_t\}$ is available, the density $p_{t_{n_t}}$ will be approximated by $p^{\Delta t}_{n_t}$, with the transition from $p^{\Delta t}_{n_t-1}$ to $p^{\Delta t}_{n_t}$ divided into two steps, as outlined in the main text.

**Algorithm 1:** Splitting algorithm for the KSE with methods for PDE

**Input:** Initial density $p_0^{\Delta t}$, operator $\mathcal{A}^*$, rescaled observation function $\tilde{h}$ and rescaled observations $\{\tilde{y}_{t_{n_t}}\}_{n_t=1}^{N_t}$

**Output:** $p^{\Delta t} = (p_t^{\Delta t}, t \geq 0)$

**for** $n_t = 1, 2, \ldots$ **do**

For $t \in [t_{n_t-1}, t_{n_t}]$, solve the following partial differential equation

$$\frac{\partial p_t^{n_t}(x)}{\partial t} = \mathcal{A}^* p_t^{n_t}(x), \qquad p_{t_{n-1}}^{n_t} = p_{n_t-1}^{\Delta t};$$

Define $\overline{p}_{n_t}^{\Delta t} := p_{t_{n_t}}^{n_t}(x)$

**if** $t_{n_t}$ *is an observation time* **then**

Update the density with information from observation $\tilde{y}_{t_{n_t}}$, that is, compute $p_{n_t}^{\Delta t}(x)$ for $x \in \mathbb{S}_X$ via

$$p_{n_t}^{\Delta t}(x) := \xi_{n_t} \zeta_{n_t}^{\Delta t}(x)\, \bar{p}_{n_t}^{\Delta t}(x)$$

where

$$\zeta_{n_t}^{\Delta t}(x) := \exp\left(-\frac{1}{2}\Delta t \|\tilde{y}_{n_t} - \tilde{h}(x)\|^2\right),$$

$\tilde{y}_{n_t}$, $\tilde{h}$ are rescaled versions of $y_{n_t}$ and $h$, respectively, and $\xi_{n_t}$ is a normalisation constant chosen such that

$$\int_{\mathbb{S}_X} p_{n_t}^{\Delta t}(x)\, dx = 1$$

**else**

Set $p_{n_t}^{\Delta t} \leftarrow p_{t_{n_t}}^{n_t}$

*2.3.2. Solving the Kushner-Stratonovich equation with sequential Monte Carlo*

Let $\{X^{(n_P)}\}_{n_P=1}^{N_P}$ be a collection of $N_P$ mutually independent stochastic processes, all independent of $\widetilde{Y}$ and each being a solution to SDE (2.23). Then the pairs $\{(X^{(n_P)}, \widetilde{Y})\}_{n_P=1}^{N_P}$ are identically distributed and have the same distribution as the pair $(X, \widetilde{Y})$ under $\mathbb{Q}$. Additionally, for $n_P = 1, \ldots, N_P$, $0 \leq t < \infty$, define $w_t^{(n_P)}$ as

$$w_t^{(n_P)} = \exp\left(\widetilde{h}^\top \left(X_t^{(n_P)}\right) \widetilde{Y}_t - \int_0^t \widetilde{Y}_s^\top d\widetilde{h}\left(X_s^{(n_P)}\right) - \frac{1}{2} \int_0^t \|\widetilde{h}\left(X_s^{(n_P)}\right)\|^2 ds\right).$$

Then the triples $\{(X^{(n_P)}, w^{(n_P)}, \widetilde{Y})\}_{n_P=1}^{N_P}$ are identically distributed and have the same distribution as the triple $(X, L, \widetilde{Y})$ under $\mathbb{Q}$.

For $n_P = 1, \ldots, N_P$, $w^{(n_P)}$ can be computed recursively according to

$$w_{t_{n_t}}^{(n_P)} = w_{t_{n_t-1}}^{(n_P)} \times \exp\left(\widetilde{h}^\top \left(X_s^{(n_P)}\right) \widetilde{Y}_s\Big|_{t_{n_t-1}}^{t_{n_t}} - \int_{t_{n_t-1}}^{t_{n_t}} \widetilde{Y}_s^\top d\widetilde{h}\left(X_s^{(n_P)}\right) - \frac{1}{2} \int_{t_{n_t-1}}^{t_{n_t}} \|\widetilde{h}\left(X_s^{(n_P)}\right)\|^2 ds\right)$$

whenever a new rescaled observation $\widetilde{y}_{t_{n_t}}$ becomes available.

Lastly, the $N_P$ triples may also be used to construct the measure-valued processes $\rho^{N_P} = (\rho_t^{N_P}, t \geq 0)$ and $\pi^{N_P} = (\pi_t^{N_P}, t \geq 0)$:

$$\rho_t^{N_P} := \frac{1}{N_P} \sum_{n_P=1}^{N_P} w_t^{(n_P)} \delta_{X_t^{(n_P)}}, \qquad \pi_t^{N_P} := \frac{\rho_t^{N_P}}{\rho_t^{N_P}(\mathbb{1})} = \sum_{n_P=1}^{N_P} \bar{w}_t^{(n_P)} \delta_{X_t^{(n_P)}},$$

where $\bar{w}_t^{(n_P)}$ is given by $\bar{w}_t^{(n_P)} = \left(\sum_{n_P'=1}^{N_P} w_t^{(n_P)'}\right)^{-1} w_t^{(n_P)}$, $n_P = 1, \ldots, N_P$, $0 \leq t < \infty$, and $\delta$ is the Dirac's delta generalised function; that is, $\rho_t^{N_P}$ is the empirical measure of $N_P$ particles with positions $\{X_t^{(n_P)}\}_{n_P=1}^{N_P}$, and $\pi_t^{N_P}$ is its normalised version.

**Algorithm 2:** Continuous-Discrete Bootstrap Particle filter

**Input:** Initial density $p_0^{\Delta t}$, coefficients $f$ and $g$, rescaled observation function $\tilde{h}$, rescaled observations $\{\tilde{y}_{t_{n_t}}\}_{n_t=1}^{N_t}$, number of particles $N_P$, and resampling method

**Output:** $p^{\Delta t} = (p_t^{\Delta t}, t \geq 0)$

Sample $X_0^{(n_P)}, n_P = 1, \ldots, N_P$ mutually independent random variables from $p_0^{\Delta t}$, each with initial weight $1/N_P$

**for** $n_t = 1, 2, \ldots$ **do**

```
/* Use a resampling step to eliminate particles with low weights
   and multiply those with high weights                          */
```

Resample $\{X_{t_{n_t-1}}^{(n_P)}, w_{t_{n_t-1}}^{(n_P)}\}_{n_P=1}^{N_P}$, resulting in equally weighted particles $\{\tilde{X}_{t_{n-1}}^{(n_P)}, 1/N_P\}_{n_P=1}^{N_P}$

**for** $n_P = 1, 2, \ldots, N_P$ **do**

/* Evolve the particles from $t = t_{n_t-1}$ to $t = t_{n_t}$ */

$X_{t_n}^{(n_P)} = \tilde{X}_{t_{n-1}}^{(n_P)} + \int_{t_{n_t-1}}^{t_{n_t}} f(\tilde{X}_t^{(n_P)})\, dt + \int_{t_{n_t-1}}^{t_{n_t}} g(\tilde{X}_t^{(n_P)})\, dB_t^X$

**if** $t_n$ *is an observation time* **then**

Update the density with information from observation $\tilde{y}_{t_{n_t}}$, that is, update the weights $w_{t_{n_t}}^{(n_P)}$ via $w_{t_{n_t}}^{(n_P)} \leftarrow w_{t_{n_t-1}}^{(n_P)} \times$

$\exp\left(\tilde{h}^\top(X_s^{(n_P)})\tilde{Y}_s\Big|_{t_{n_t-1}}^{t_{n_t}} - \int_{t_{n_t-1}}^{t_{n_t}} \tilde{Y}_s^\top d\tilde{h}(X_s^{(n_P)}) - \frac{1}{2}\int_{t_{n_t-1}}^{t_{n_t}} \|\tilde{h}(X_s^{(n_P)})\|^2 ds\right)$ with the pathwise approximation $\tilde{Y}_{t_{n_t}} - \tilde{Y}_{t_{n_t-1}} = \Delta t \cdot \tilde{y}_{t_{n_t}}$

Normalise the weights with $\bar{w}_{t_{n_t}}^{(n_P)} = \dfrac{w_{t_{n_t}}^{(n_P)}}{\sum_{n_P'=1}^{N_P} w_{t_{n_t}}^{(n_P')}}$

Set $w_{t_{n_t}}^{(n_P)} \leftarrow \bar{w}_{t_{n_t}}^{(n_P)}$

Construct $p_{n_t}^{\Delta t}$ by density estimation techniques

**The Kalman-Bucy filter**

Before presenting another approximation method, let us first examine a case for which the Kushner-Stratonovich equation has a closed-form solution. Consider the signal process with evolution in time as described in the main text, that is

$$\begin{aligned} dX_t &= f_t(X_t)\,dt + g_t(X_t)\,dB_t^X, \quad X_0 = x_0, \\ dY_t &= h_t(X_t)\,dt + k_t(X_t)\,dB_t^Y, \quad Y_0 = 0. \end{aligned}$$

As a starting point, let us assume that $f_t$ and $h_t$ are linear functions of the state, the value of $g_t$ does not vary with the state variable, and $k_t(X_t) = K$, where $K$ is a matrix of size $N_Y \times N_Y$. It is possible to rewrite our state-space model as:

$$\begin{aligned} dX_t &= f(t)X_t + g(t)dB_t^X, \quad X_0 = x_0, \\ dY_t &= h(t)X_t + KdB_t^Y, \quad Y_0 = 0. \end{aligned}$$

Now suppose that

- $x_0$ is a Gaussian random variable with mean $m_0$ and covariance $P_0$;
- $K$ is invertible;
- $f(t)$, $g(t)$, and $h(t)$ are continuous.

**Theorem 1 (Kalman-Bucy filter).** *The estimates for the conditional mean $m_t := \mathbb{E}\left[X_t|\mathcal{F}_t^Y\right]$ and error covariance $P := \mathbb{E}\left[(X_t - m_t)(X_t - m_t)^\top\right]$ form the solution to*

$$\begin{aligned} dm_t &= f(t)m_t\,dt + P_t\left(K^{-1}h(t)\right)^\top dV_t, \\ \frac{dP_t}{dt} &= f(t)P_t + P_t f^\top(t) - P_t h^\top(t)\left(KK^\top\right)^{-1} h(t)P_t + KK^\top, \end{aligned}$$

*with initial conditions $m_0$ and $P_0$.*

It is known that the solution $p_t$ to this filtering problem refers to the density of a Gaussian distribution with mean $m_t$ and covariance matrix $P_t$. One can verify this claim by noting that

$$q_t(x) \propto \exp\left(-\frac{1}{2}(x - m_t)^\top P_t^{-1}(x - m_t)\right)$$

is the density function of the unnormalised conditional distribution $\rho_t$. $q_t$ being a density of a Gaussian distribution automatically implies that $p_t$ has the same property.

As presented in the main manuscript, once we fix the initial condition $x_0 \sim \mathcal{N}(m_0, P_0)$, the exact solution to the linear Gaussian filtering problem is given by the Kalman-Bucy filter [9]. Starting at iteration $n_1 = 1$, we propagate the estimates via the procedure below.

- The *prediction* step consists of finding the estimate for the predictive mean $m_t^-$ satisfying

  $$\frac{dm_t^-}{dt} = f(t)m_t^-, \quad m_{t_{n_t-1}}^- = m_{t_{n_t-1}},$$

  and the estimate for the predictive covariance $P_t^-$ satisfies the equation

  $$\frac{dP_t^-}{dt} = f(t)P_t^- + P_t^- f^\top(t) + gg^\top(t), \quad P_{t_{n_t-1}}^- = P_{t_{n_t-1}}.$$

- The *correction* step consists of using the new observation $y_{n_t}$ at time step $t_{n_t}$. The state mean $m_{t_{n_t}}$ and covariance $P_{t_{n_t}}$ are updated according to

$$
\begin{aligned}
U_{n_t} &= P^-_{t_{n_t}} h^\top(t_{n_t}) \left( h(t_{n_t}) P^-_{t_{n_t}} h^\top(t_{n_t}) + KK^\top \right)^{-1}, \\
m_{t_{n_t}} &= m_{t_{n_t}} + U_{n_t} \left( y_{n_t} - h(t_{n_t}) m^-_{t_{n_t}} \right), \\
P_{t_{n_t}} &= \left( I_{N_X \times N_X} - U_{n_t} h(t_{n_t}) \right) P^-_{t_{n_t}} \left( I_{N_X \times N_X} - U_{n_t} h(t_{n_t}) \right) + U_{n_t} K (U_{n_t} K)^\top.
\end{aligned}
$$

**Algorithm 3:** Continuous-Discrete Kalman-Bucy filter

**Input:** Initial state estimates $m_0$, $P_0$ and observations $\{y_{t_{n_t}}\}_{n_t=1}^{N_t}$

**Output:** State variable estimates $m = (m_t, t \geq 0)$ , $P = (P_t, t \geq 0)$

```
for n_t = 1, 2, ... do
    /* Compute predicted state mean and covariance                */
    Compute the predicted state mean and covariance (m^-(t), P^-(t)) as solutions to

        dm_t^-/dt = f(t) m_t^-,     dP_t^-/dt = f(t) P_t^- + P_t^- f^T(t) + g g^T(t)

    for t ∈ [t_{n_t - 1}, t_{n_t}], with initial conditions m^-_{n_t - 1} = m_{n_t - 1} and P^-_{n_t - 1} = P_{n_t - 1}
    if t_{n_t} is an observation time then
        /* Update state estimates                                 */
        Compute the residual e_{n_t} = y_{n_t} - h(t_{n_t}) m^-_{n_t}
        Compute the innovation covariance Λ_{n_t} = h(t_{n_t}) P^-_{t_{n_t}} h^T(t_{n_t}) + K K^T
        Compute the Kalman gain U_{n_t} = P^-_{t_{n_t}} (h(t_{n_t}) m^-_{n_t})^T Λ^{-1}_{n_t}
        Update the mean estimate m_{n_t} = m^-_{n_t} + U_{n_t} e_{n_t}
        Update the covariance estimate P_{n_t} = P^-_{t_{n_t}} - U_{n_t} S^{-1}_{n_t} U^T_{n_t}
    else
        Set m_{n_t} ← m^-_{n_t} Set P_{n_t} ← P^-_{n_t}
```

**The extended Kalman filter**

We proceed to detail a linearisation method to cope with non-linear models, for which $f$ and/or $h$ are non-linear functions on $\mathbb{S}_X$. We present the most common approach, the extended Kalman filter. The method assumes that the conditional density is nearly Gaussian, so that third and higher-order odd central moments are essentially zero, and the fourth and higher-order even central moments can be expressed in terms of the covariance.
We apply the assumption above to truncate high-order terms in a Taylor series expansion to functions $f$ and $h$:

Let $\overline{x}_t$ be a solution of the ODE $\frac{d\overline{x}}{dt} = f(\overline{x}_t), \overline{x}_0 = m_0$. Consider a small-time frame for which the contribution of the stochastic terms $g(X_t)\,dB_t^X$ and $k(t)dB_t^Y$ remains small, such that a trajectory $t \mapsto X_t$ may be viewed as a perturbation from the deterministic trajectory $t \mapsto \overline{x}_t$. Instead of solving the filtering problem for the original system of differential equations, we work with the following Taylor-like expansion:

$$
\begin{aligned}
dX_t \simeq & F\left((\overline{x}_t)\left(X_t - \overline{x}_t\right) + f\left(\overline{x}_t\right)\right)dt + g\left(\overline{x}_t\right)dB_t^X, \\
dY_t \simeq & H\left((\overline{x}_t)\left(X_t - \overline{x}_t\right) + h\left(\overline{x}_t\right)\right)dt + k(t)dB_t^Y,
\end{aligned}
$$

where, $\simeq$ means approximately equal without a rigorous mathematical meaning to it, and $F$ and $H$ are the derivatives of $f$ and $h$, that is

$$
F\left(\overline{x}_t\right) := \left.\frac{\partial f}{\partial x}\right|_{x=\overline{x}_t}, \qquad H\left(\overline{x}_t\right) := \left.\frac{\partial h}{\partial x}\right|_{x=\overline{x}_t}.
$$

If we let $\overline{x}_t \equiv m_t$, the resulting equations form the extended Kalman filter.

- For the time interval $[t_{n_t-1}, t_{n_t}]$, the *prediction* step consists in finding the predictive estimate $m_t^-$ for the mean satisfying

$$
\frac{dm_t^-}{dt} = f\left(m_t^-\right), \quad m_{t_{n_t-1}}^- = m_{t_{n_t-1}},
$$

  and the predictive estimate $P_t^-$ for the covariance satisfying

$$
\begin{aligned}
\frac{dP_t^-}{dt} =& F\left(m_t^-\right)P_t^- + P_t^- F^\top\left(m_t^-\right) + gg^\top\left(m_t\right), \\
P_{t_{n_t-1}}^- =& P_{t_{n_t-1}}, \\
F(m_t^-) :=& \left.\frac{\partial f}{\partial x}\right|_{x=m_t^-}.
\end{aligned}
$$

- The *correction* step consists of using the new observation $y_{n_t}$ at time step $t_{n_t}$. The state mean $m_{t_{n_t}}$ and covariance $P_{t_{n_t}}$ are updated according to

$$
\begin{aligned}
H\left(m_{t_{n_t}}\right) &:= \left.\frac{\partial h}{\partial x}\right|_{x=m_{t_{n_t}}}, \\
U_{n_t} &= P_{t_{n_t}}^- H^\top\left(m_{n_t}\right)\left(H\left(m_{t_{n_t}}\right)P_{t_{n_t}}^- H^\top\left(m_{t_{n_t}}\right) + KK^\top\right)^{-1}, \\
m_{t_{n_t}} &= m_{t_{n_t}}^- + U_{n_t}\left(y_{n_t} - h\left(m_{t_{n_t}}\right)\right), \\
P_{t_{n_t}} &= \left(I_{N_X\times N_X} - K_{n_t}H^\top\left(m_{t_{n_t}}\right)\right)P_{t_{n_t}}^-\left(I_{N_X\times N_X} - U_{n_t}H^\top\left(m_{t_{n_t}}\right)\right) + U_{n_t}K\left(U_{n_t}K\right)^\top.
\end{aligned}
$$

To cope with numerical errors, Kalman filter equations are often implemented such that the matrix square roots of covariance matrices are used in computations instead of their actual values. Following the derivation in [10] and [11], we can rewrite the extended Kalman filter equations in terms of generalised or modified Cholesky factors. Let $\Lambda_t \Lambda_t^\top = P_t$, then the differential equation for either $\frac{d\Lambda_t}{dt}$ or $\frac{d\Lambda_t^\top}{dt}$ is solved. For the standard implementation, the problem requires ODE solvers with variable step size for their accurate and efficient numerical integration [12, 13, 14]. Lastly, it may be beneficial to maintain symmetry of $P_t$ by evaluating the expression $P_t = 0.5 * (P_t + P_t^\top)$ at every iteration [15].

**Algorithm 4:** Squared-root Extended Kalman filter

**Input:** Initial state estimates $m_0$, $\Lambda_0$ such that $\Lambda_0\Lambda_0^\top = P_0$, and observations $\{y_{t_{n_t}}\}_{n_t=1}^{N_t}$

**Output:** State variable estimates $m = (m_t, t \geq 0)\,, \Lambda = (\Lambda_t, t \geq 0)\,,$

Define the matrix-valued function $\Phi(\cdot)$ that returns the lower triangular part of an argument matrix $M$ as follows:

$$
\begin{aligned}
\Phi_{ij}(M) &= M_{ij} \quad \text{if} \quad i > j \\
\Phi_{ii}(M) &= 0.5M_{ii} \\
\Phi_{ij}(M) &= 0 \qquad \text{if} \quad i < j
\end{aligned}
$$

**for** $n_t = 1, 2, \ldots$ **do**

```
/* Compute predicted state estimators */
```

Compute the predicted state estimators $(m_t^-, \Lambda_t^-)$ as solutions to

$$
\frac{dm_t^-}{dt} = f(m_t^-)dt, \qquad \frac{d\Lambda_t^-}{dt} = \Lambda_t\Phi(A_t + A_t^\top + C_t)
$$

where

$$
A_t = (\Lambda_t)^{-1}F_tS_t, \quad C_t = (\Lambda_t)^{-1}gg^\top(m_t^-)((\Lambda_t)^{-1})^\top, \quad F_t = \left.\frac{\partial f}{\partial x}\right|_{x=m_t^-}
$$

for $t \in [t_{n_t-1}, t_{n_t}]$, with initial conditions $m_{t_{n-1}}^- = m_{t_{n-1}}$ and $\Lambda_{t_{n-1}}^- = \Lambda_{t_{n-1}}$

**if** $t_{n_t}$ *is an observation time* **then**

```
/* Update state estimates using QR decomposition */
```

Start with a QR decomposition of the matrix on the LHS of the following equation

$$
\begin{pmatrix} K(t_{n_t} - t_{n_t-1})^{1/2} & H_x\left(m_{t_{n_t}}^-\right)\Lambda_{t_{n_t}}^- \\ 0^\top & \Lambda_{t_{n_t}}^- \end{pmatrix}^\top = Q_{n_t}\begin{pmatrix} R_{n_t}^{1/2} & 0 \\ \tilde{U}_{n_t} & \Lambda_{t_{n_t}} \end{pmatrix}^\top
$$

where $H_x(m) \equiv \left.\frac{\partial h}{\partial x}\right|_{x=m}$

Extract $Q_{n_t}$ as the $Q$-factor and let the last matrix on the RHS be the $R$-factor. Directly extract $\Lambda_{t_{n_t}}$ from the latter

For the mean estimate, let

$$
e_{n_t} = y_{n_t} - h\left(m_{t_{n_t}}^-\right),\ U_{n_t} = \tilde{U}_{n_t}((R_{n_t}^{1/2})^{-1})^\top,\ m_{t_{n_t}} = m_{t_{n_t}}^- + U_{n_t}e_{n_t}
$$

**else**

Set $m_{t_{n_t}} \leftarrow m_{t_{n_t}}^-$

Set $\Lambda_{t_{n_t}} \leftarrow \Lambda_{t_{n_t}}^-$

### 3. Chemostat equations

Let $b$ denote the concentration of biomass and $s$ that of substrate. Consider the following reaction to produce one unit of biomass and $\gamma$ units of methane (biogas) from $\kappa$ units of substrate:

$$\kappa\text{substrate} \xrightarrow{\lambda(\cdot)} \text{biomass} + \gamma\text{biogas},$$

where $\lambda(\cdot)$ is the reaction rate. Since biomass here is not a chemically defined compound, we make use of the so-called yield coefficients $\kappa$ and $\gamma$ under the assumptions that the biomass composition is constant and that the surface-to-volume ratio of microbial cells is constant at a population level [16]. These are mild assumptions when one is interested not in the full details of intracellular reactions but rather in the macroscopic behaviour of the system. Additionally, let us assume that $\lambda(\cdot) = \mu(\cdot)b$, where $\mu$ is called "specific growth velocity". With this assumption, we guarantee that the reaction rate is zero in the absence of biomass.

At a specific time $t$, let $F_t^{in}$ express how much medium flows into the vessel per hour. The incoming dilution rate is calculated by dividing the incoming flow rate by the culture volume $V_t$, that is $D_t^{in} = F_t^{in}/V_t$. Analogously, $F_t^{out}$ and $D_t^{out}$ denote the outgoing flow rate and its corresponding dilution rate, respectively.

To establish the equations of the chemostat, we apply a mass balance in which, over a period of time $dt$, the variation in the mass of an element is the net result of four possible terms: the quantity of that element that has been brought into the system, the produced quantity in the vessel, the consumed quantity inside the vessel and the extracted quantity.

We start with the equation for the change in volume:

$$\frac{dV_t}{dt} = F_t^{in} - F_t^{out},$$

and the chain rule on the derivative of the quantity of substrate with respect to time:

$$\frac{d(s_tV_t)}{dt} = s_t(F_t^{in} - F_t^{out}) + \frac{ds_t}{dt}V_t. \tag{11}$$

The variation in the substrate mass $d(s_tV_t)/dt$ is the balance between the four possible terms outlined above. After rearranging the terms in Eq. (11), we get

$$\begin{aligned} V_t\frac{ds_t}{dt} =& \frac{d(s_tV_t)}{dt} - s_t(F_t^{in} - F_t^{out}) \\ =& \left(s_t^{in}F_t^{in} - s_tF_t^{out} - \kappa V_t\lambda(\cdot)\right) - s_t(F_t^{in} - F_t^{out}) \\ =& F_t^{in}(s_t^{in} - s_t) - \kappa V_t\lambda(\cdot). \end{aligned}$$

Analogously, we apply the chain rule for the biomass with respect to time:

$$\begin{aligned} V_t\frac{db_t}{dt} =& \frac{d(b_tV_t)}{dt} - b_t(F_t^{in} - F_t^{out}) \\ =& \left(b_t^{in}F_t^{in} - b_tF_t^{out} + V_t\lambda(\cdot)\right) - b_t(F_t^{in} - F_t^{out}) \\ =& V_t\lambda(\cdot) - F_t^{in}(b_t - b_t^{in}). \end{aligned}$$

The following are the resulting balance equations:

$$\frac{dV_t}{dt} = F_t^{in} - F_t^{out}, \quad \frac{db_t}{dt} = \lambda(\cdot) - D_t^{in}(b_t - b_t^{in}), \quad \frac{ds_t}{dt} = D_{in}(s_t^{in} - s_t) - \kappa\lambda(\cdot).$$

The focus is on bioreactor processes performed at continuous mode, where $F_t^{in} = F_t^{out} = F$, $V_t = V$ and $D_t^{in} = D_t^{out} = D$. Under the previous assumption $\lambda(\cdot) = \mu(\cdot)b$, the ordinary differential model for the growth of a single species in a chemostat is

$$\frac{db_t}{dt} = (\mu(\cdot) - D)b_t + Db_t^{in}, \qquad \frac{ds_t}{dt} = D(s_t^{in} - s_t) - \kappa\mu(\cdot)b_t,$$

with initial concentrations $b_0$ and $s_0$.

### 4. Numerical experiments

Table 2: Additional parameters for differential equations of $X$ and $Y$ (Extension of Table 2 of the main material).

| | Model parameters | | Simulation parameters | |
|---|---|---|---|---|
| | Obs. function | $K$ | $\Delta t^x$ (hours) | $\Delta t^y$ (hours) |
| 1A - high-freq. obs. | $27.5\mu(s)b$ | 0.01 | 0.01 | 0.01 |
| 1A - low-freq. obs. | $27.5\mu(s)b$ | $0.01\left(\frac{\Delta t^y}{\Delta t^x}\right)^{0.5}$ | 0.01 | 2 |
| 1B - low-freq. obs. | $s$ | $0.02\left(\frac{\Delta t^y}{\Delta t^x}\right)^{0.5}$ | 0.01 | 2 |
| 1C - low-freq. obs. | $b$ | $0.02\left(\frac{\Delta t^y}{\Delta t^x}\right)^{0.5}$ | 0.01 | 2 |
| 1D - low-freq. obs. | $(b, s)$ | $0.02\left(\frac{\Delta t^y}{\Delta t^x}\right)^{0.5} I_{2\times2}$ | 0.01 | 2 |
| 2A - high-freq. obs. | $2.75\mu(s)b$ | 0.01 | 0.005 | 0.005 |
| 2A - low-freq. obs. | $2.75\mu(s)b$ | $0.01\left(\frac{\Delta t^y}{\Delta t^x}\right)^{0.5}$ | 0.005 | 2 |
| 2B - low-freq. obs. | $s$ | $0.02\left(\frac{\Delta t^y}{\Delta t^x}\right)^{0.5}$ | 0.005 | 2 |
| 2C - low-freq. obs. | $b$ | $0.02\left(\frac{\Delta t^y}{\Delta t^x}\right)^{0.5}$ | 0.005 | 2 |
| 2D - low-freq. obs. | $(b, s)$ | $0.02\left(\frac{\Delta t^y}{\Delta t^x}\right)^{0.5} I_{2\times2}$ | 0.005 | 2 |

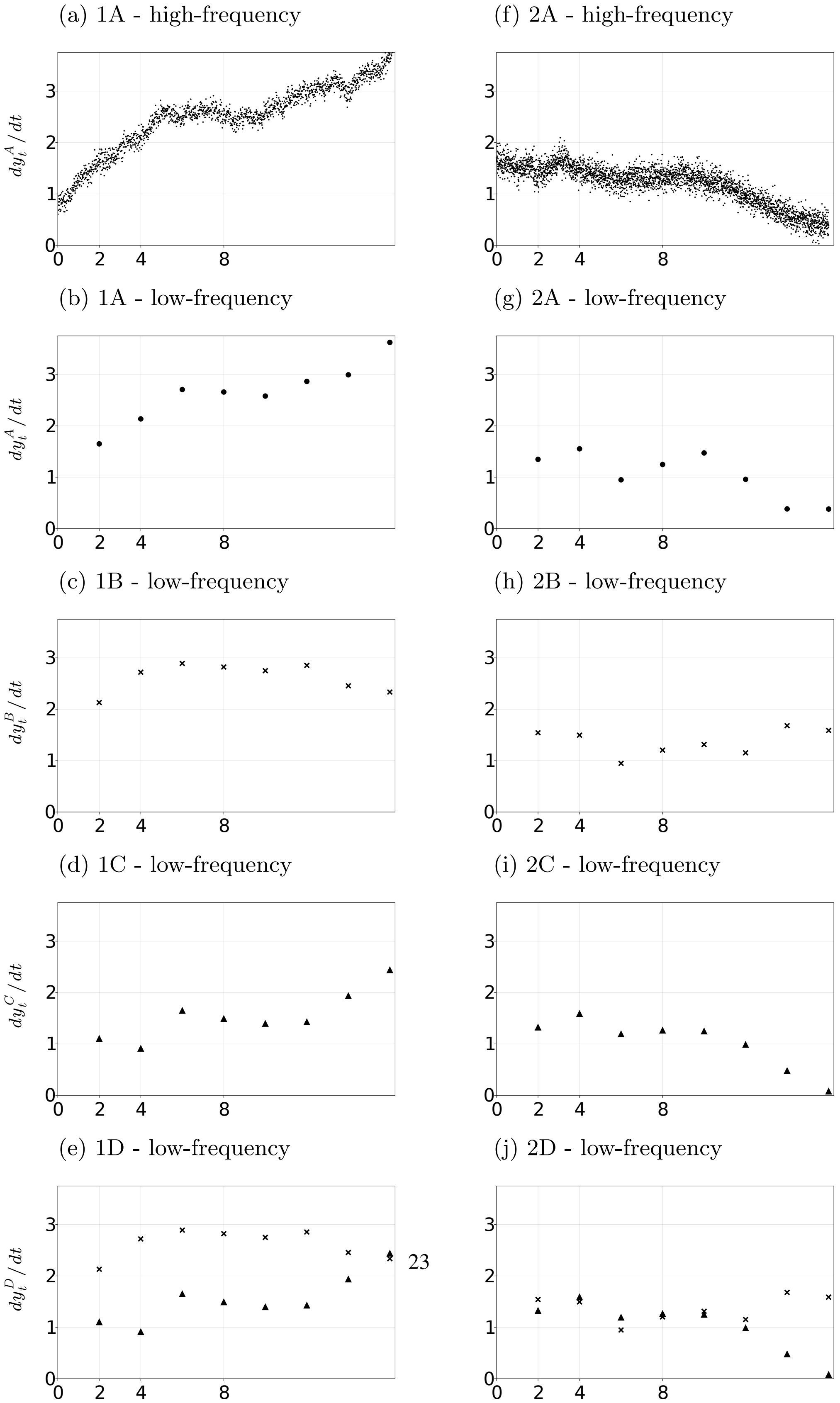

Figure 1: Observations from the trajectories highlighted in Figure 3 of the main material.

We now present the pathwise approximation to the solution of the Kushner-Stratonovich equation for the growth function of the Monod type with different observation scenarios. $X_0 \sim \mathcal{N}([1,\ 1]^\top,\ 0.05^2 I_{2\times 2})$.

For each scenario, we first show the squared Hellinger distances between the densities obtained by approximating the solution to the Kushner-Stratonovich equation with particle filters and with (i) methods for PDEs or (ii) the EKF. For the PDE methods, the discretisation in space is in the domain $(0, 5.0] \times (0, 5.0]$, and we consider three scenarios for the refinement of the grid, each containing a total count of $32^2, 128^2, 512^2$ finite volumes. The squared Hellinger distance is computed with respect to the approximation from the BPF with $N_P = 2.5 \times 10^6$. Secondly, we present snapshots of the approximations at selected times $t$. The prior knowledge from the Kolmogorov forward equation is represented by marginal density functions in gray. The updated marginal density function is represented by coloured curves. Lastly, for scenarios with sparse observations, we present the splitting-up approximation.

**1A - high-frequency observations (Monod kinetics, observations of biogas flow rate)**

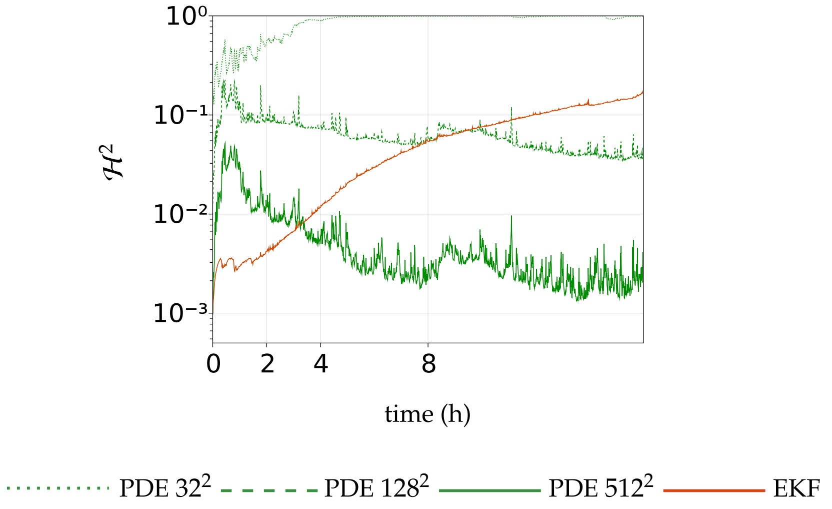

Figure 2: Squared Hellinger distance for 1A - high-frequency observations. BPF: $N_P = 2.5 \times 10^6$.

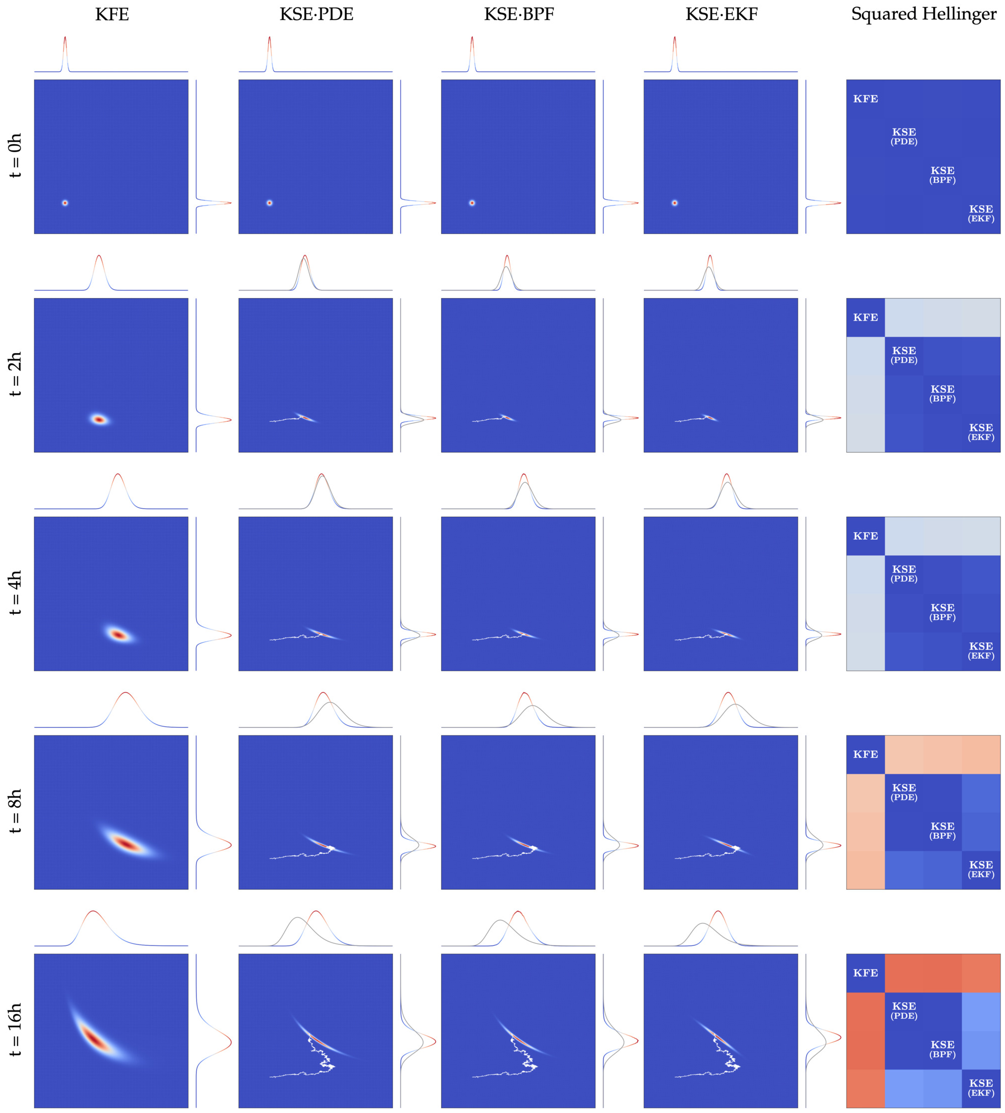

Figure 3: Approximation to the solution of the Kushner-Stratonovich equation for the growth function of the Monod type with continuous stream of observations of biogas flow rate. $256^2$ grid. $N_P = 2.5 \times 10^6$.

### 1A - low-frequency observations (Monod kinetics, sparse observations of biogas flow rate)

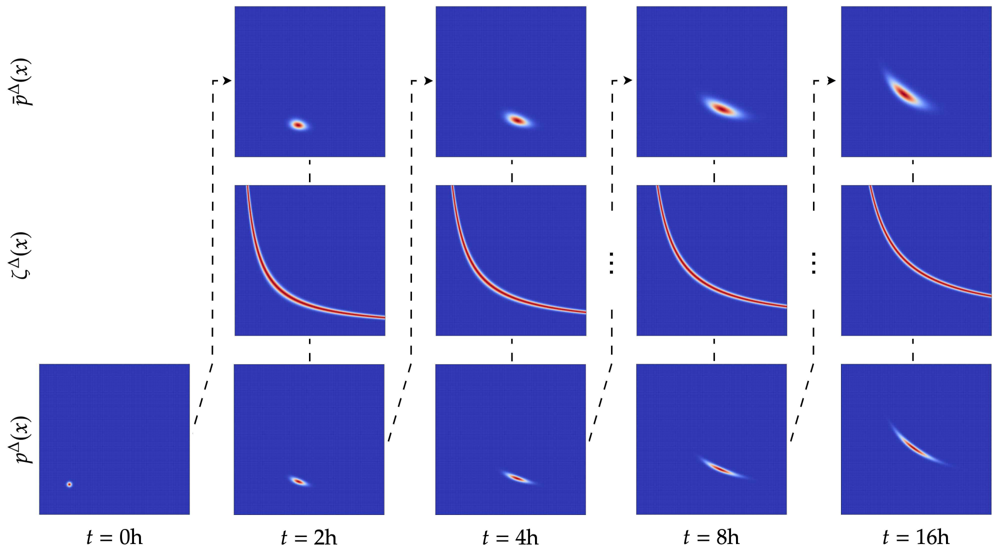

Figure 4: Splitting-up approximation for the KSE using a $256^2$ grid.

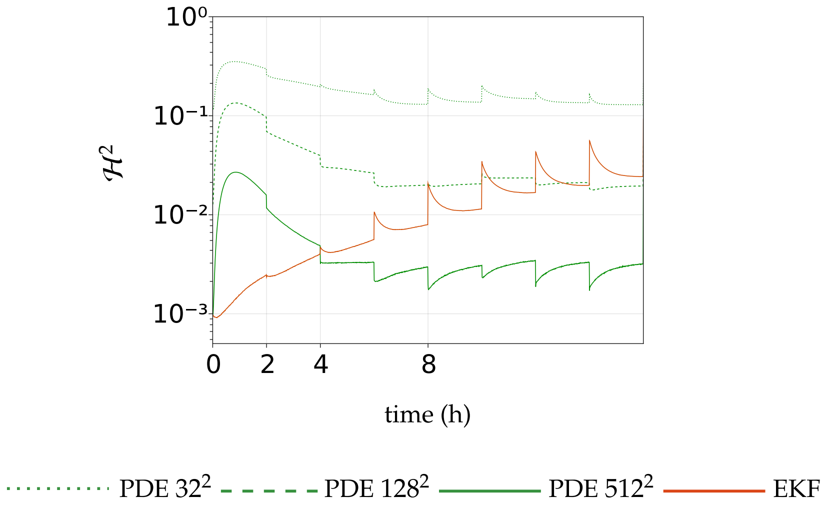

Figure 5: Squared Hellinger distance for 1A - low-frequency observations. BPF: $N_P = 2.5 \times 10^6$.

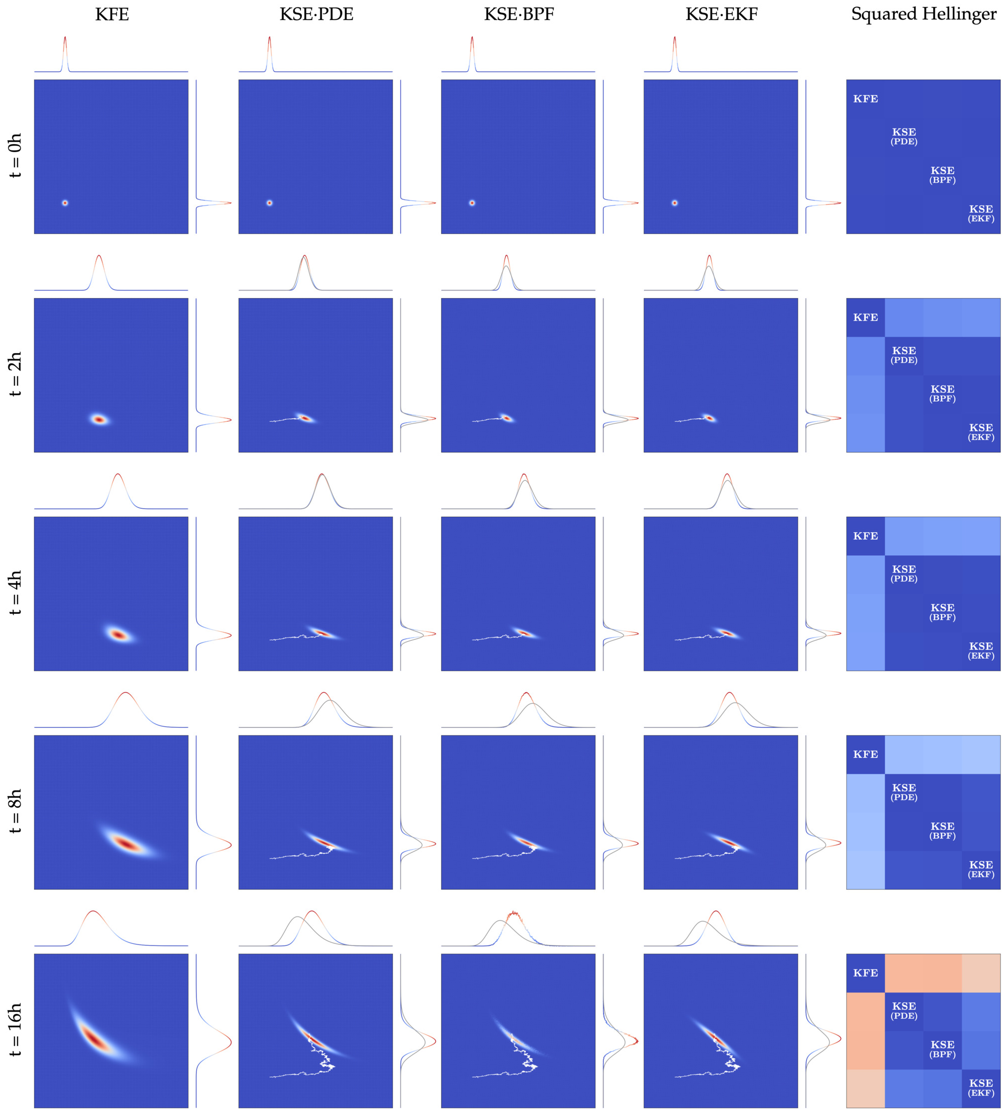

Figure 6: Approximation to the solution of the Kushner-Stratonovich equation for the growth function of the Monod type with sparse observations of biogas flow rate. $256^2$ grid. $N_P = 2.5 \times 10^6$.

**1B - low-frequency observations (Monod kinetics, sparse observations of substrate concentration)**

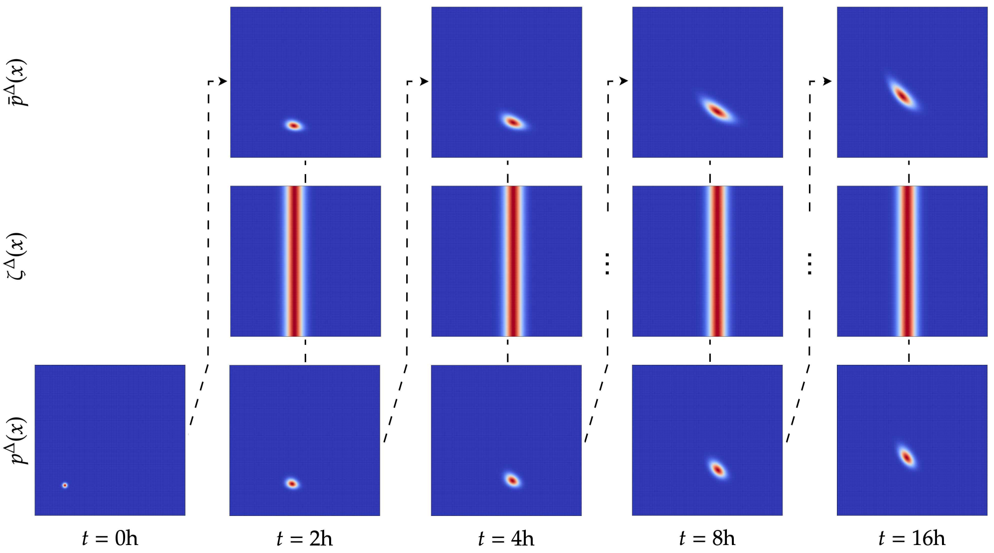

Figure 7: Splitting-up approximation for the KSE using a $256^2$ grid.

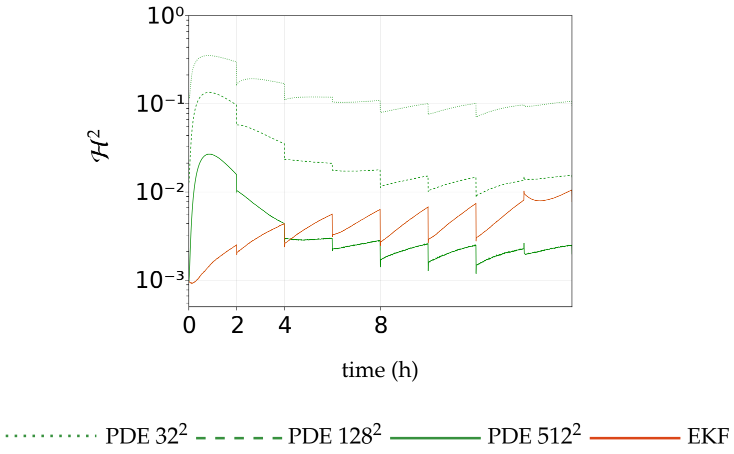

Figure 8: Squared Hellinger distance for 1B - low-frequency observations. BPF: $N_P = 2.5 \times 10^6$.

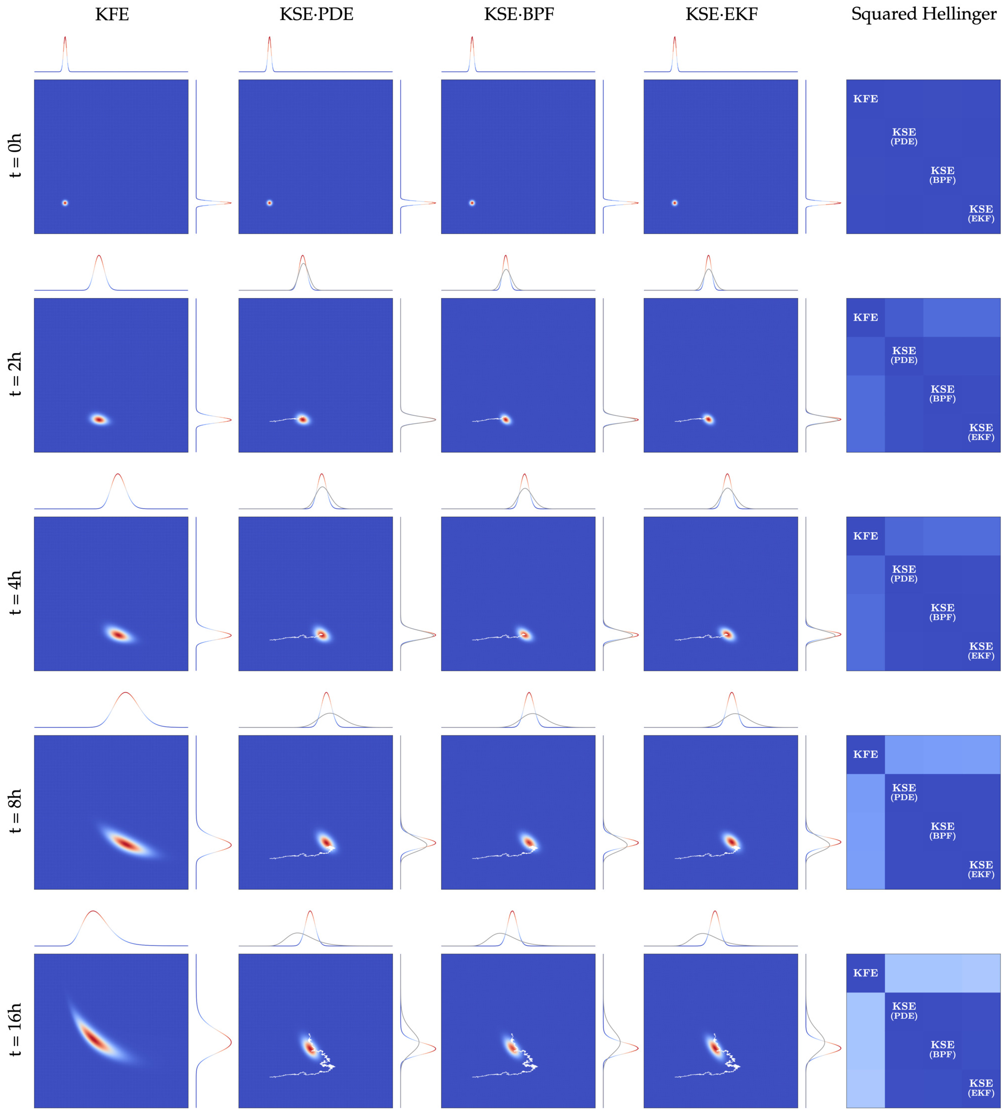

Figure 9: Approximation to the solution of the Kushner-Stratonovich equation for the growth function of the Monod type with sparse observations of substrate concentration. $256^2$ grid. $N_P = 2.5 \times 10^6$.

**1C - low-frequency observations (Monod kinetics, sparse observations of biomass concentration)**

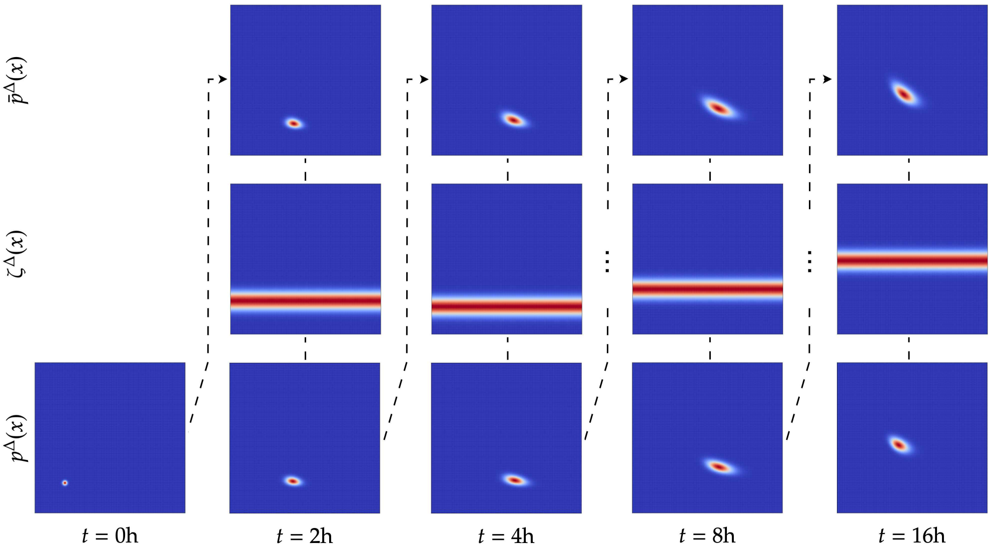

Figure 10: Splitting-up approximation for the KSE using a $256^2$ grid.

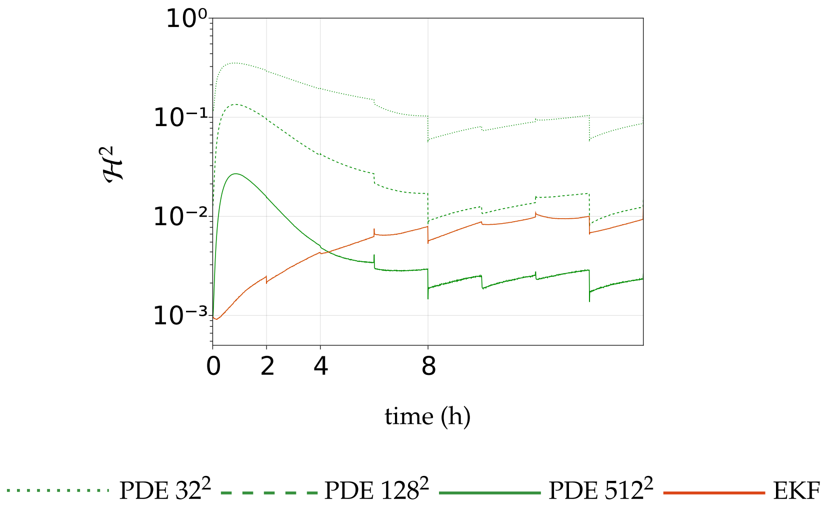

Figure 11: Squared Hellinger distance for 1C - low-frequency observations. BPF: $N_P = 2.5 \times 10^6$.

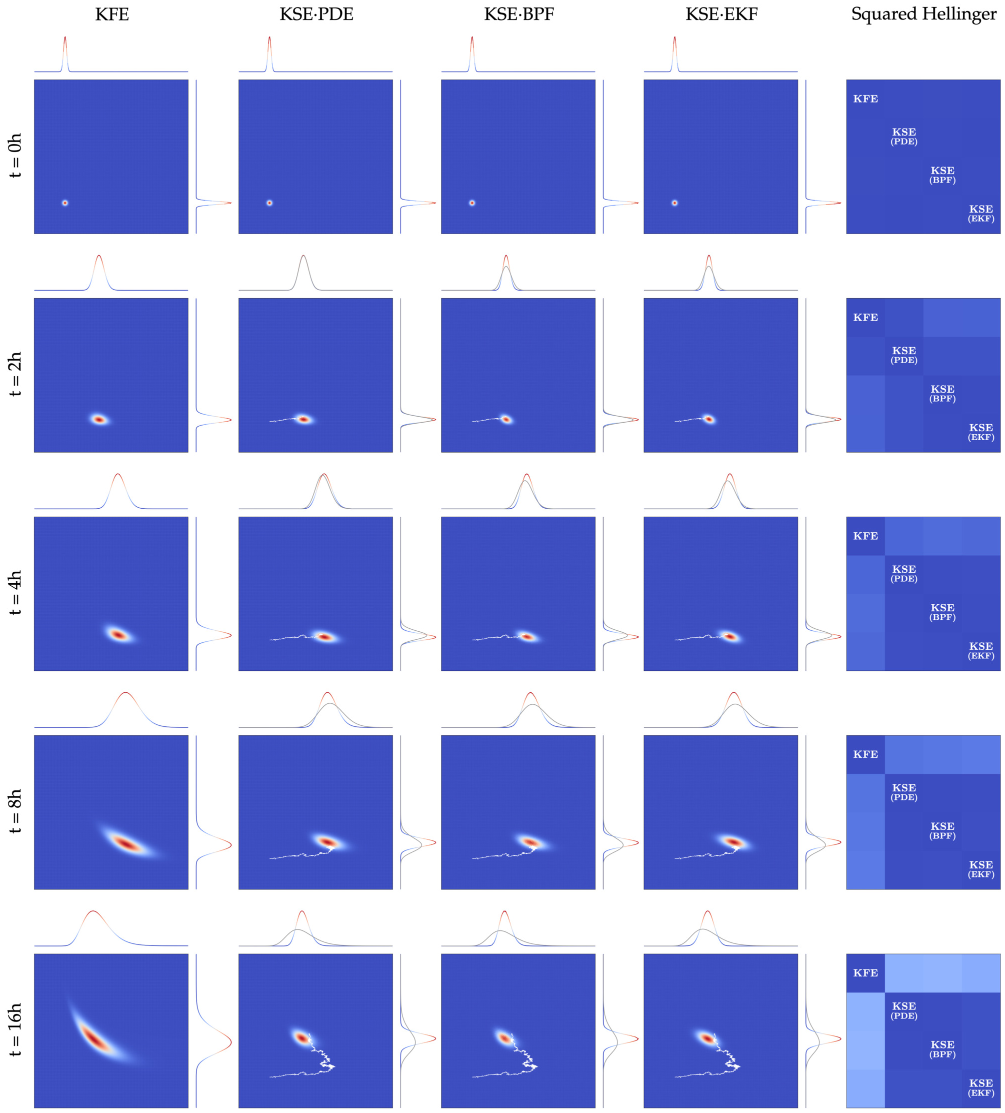

Figure 12: Approximation to the solution of the Kushner-Stratonovich equation for the growth function of the Monod type with sparse observations of biomass concentration. $256^2$ grid. $N_P = 2.5 \times 10^6$.

**1D - low-frequency observations (Monod kinetics, sparse observations of biomass and substrate concentrations)**

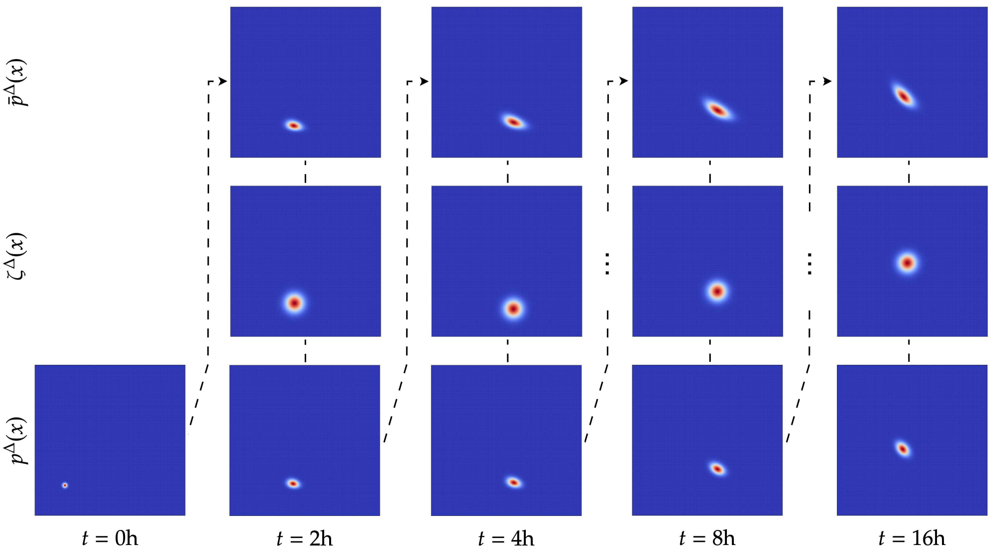

Figure 13: Splitting-up approximation for the KSE using a $256^2$ grid.

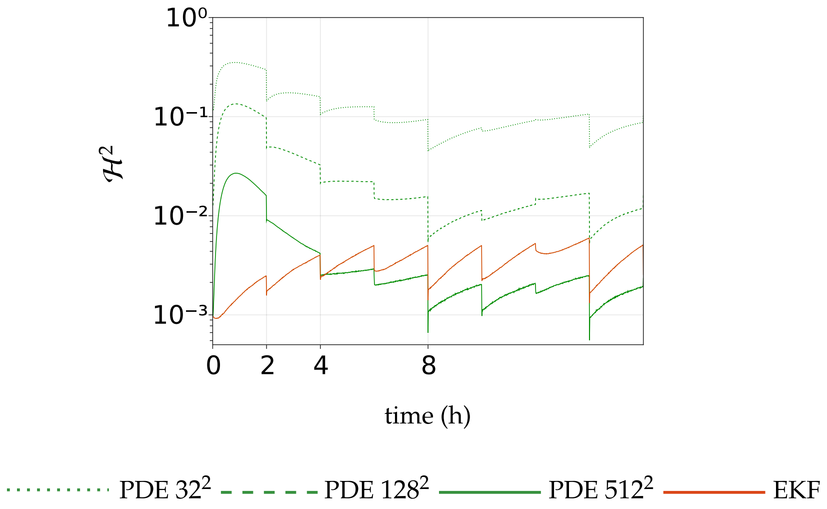

Figure 14: Squared Hellinger distance for 1D - low-frequency observations. BPF: $N_P = 2.5 \times 10^6$.

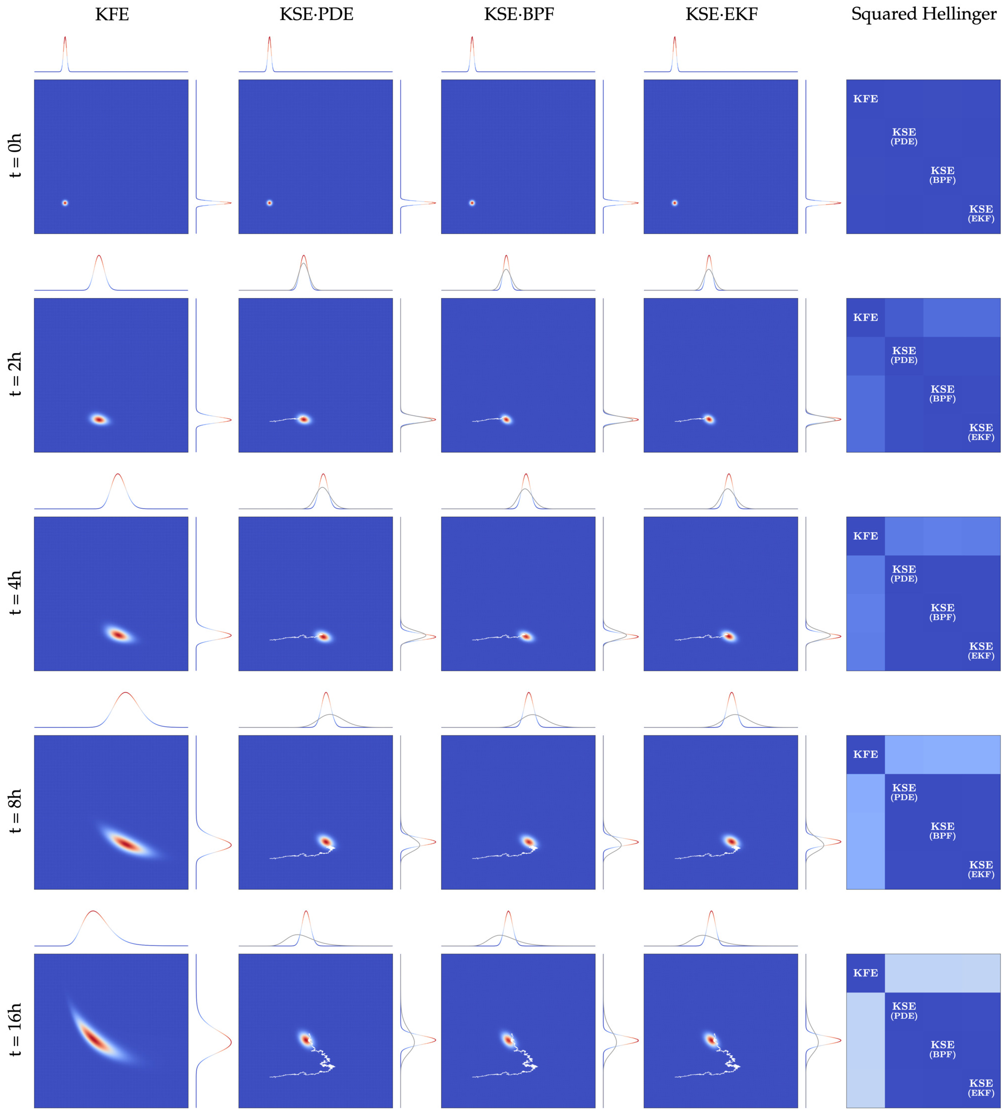

Figure 15: Approximation to the solution of the Kushner-Stratonovich equation for the growth function of the Monod type with sparse observations of biomass and substrate concentrations. $256^2$ grid. $N_P = 2.5 \times 10^6$.

We now present the pathwise approximation to the solution of the Kushner-Stratonovich equation for the growth function of the Haldane type with different observation scenarios. $X_0 \sim \mathcal{N}([1.65, 2.5]^\top, 0.05^2 I_{2\times 2})$. The prior knowledge from the Kolmogorov forward equation is represented by marginal density functions in gray. The updated marginal density function is represented by coloured curves.

For each scenario, we first show the squared Hellinger distances between the densities obtained by approximating the solution to the Kushner-Stratonovich equation with particle filters and with (i) methods for partial differential equations or (ii) the extended Kalman filter. For the PDE methods, the discretisation in space is in the domain $(0, 3.0] \times (0, 3.0]$, and we consider three scenarios for the refinement of the grid, each containing a total count of $32^2, 128^2, 512^2$ finite volumes. The squared Hellinger distance is computed with respect to the approximation from the bootstrap particle filter with $N_P = 2.5\times 10^6$. Secondly, we present snapshots of the approximations at selected times $t$. The prior knowledge from the Kolmogorov forward equation is represented by marginal density functions in gray. The updated marginal density function is represented by coloured curves. Lastly, for scenarios with sparse observations, we present the splitting-up approximation.

**2A - high-frequency observations (Haldane kinetics, observations of biogas flow rate)**

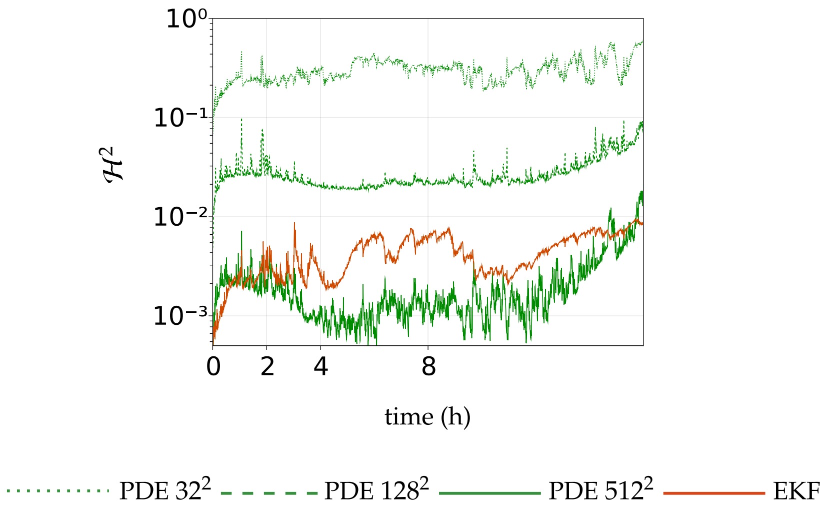

Figure 16: Squared Hellinger distance for 2A - high-frequency observations. BPF: $N_P = 2.5 \times 10^6$.

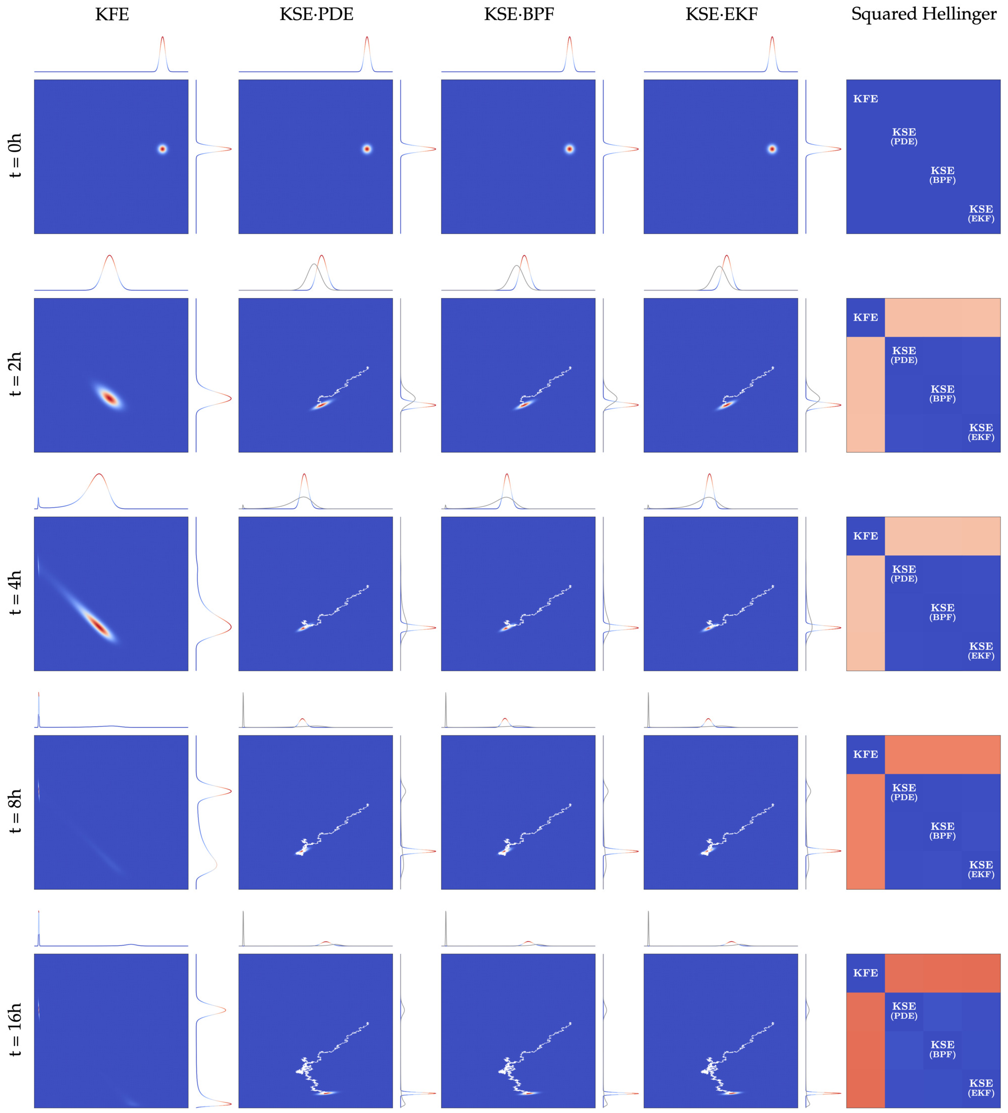

Figure 17: Approximation to the solution of the Kushner-Stratonovich equation for the growth function of the Haldane type with continuous stream of observations of biogas flow rate. $512^2$ grid. $N_P = 2.5 \times 10^6$.

**2A - low-frequency observations (Haldane kinetics, sparse observations of biogas flow rate)**

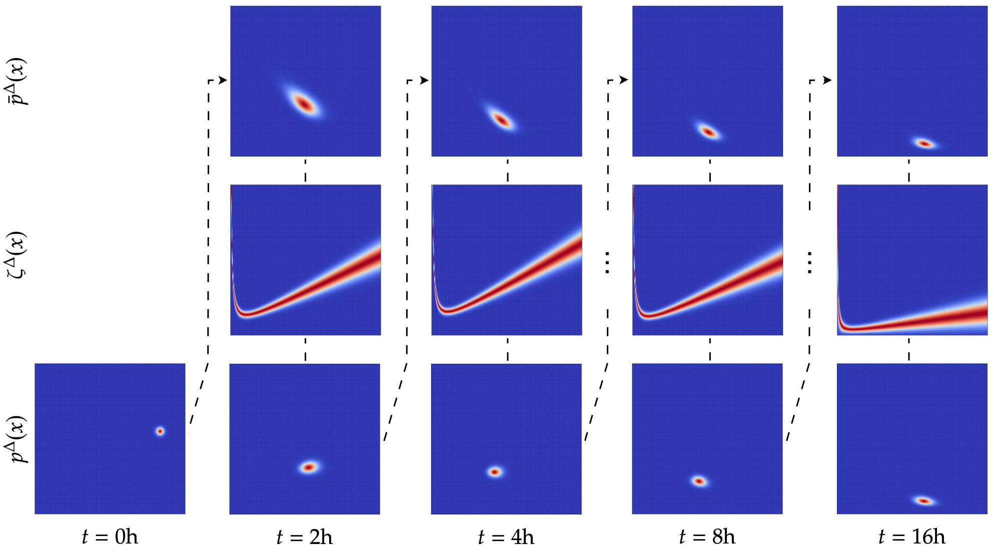

Figure 18: Splitting-up approximation for the KSE using a $256^2$ grid.

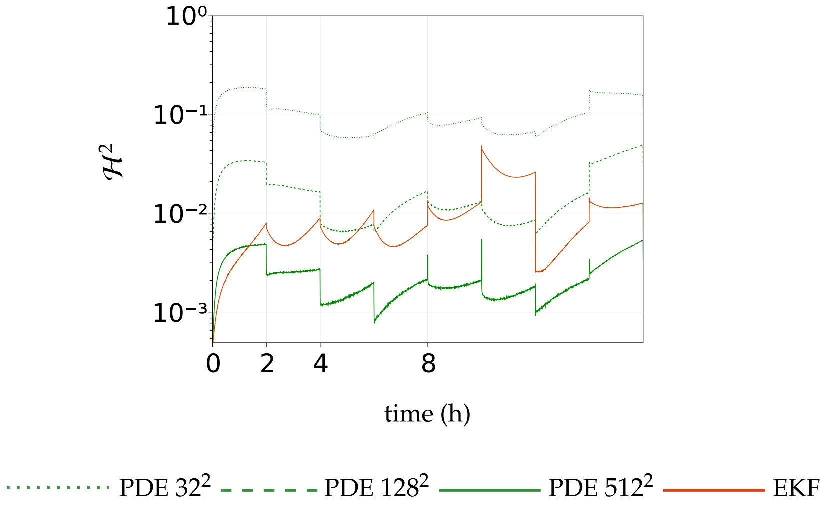

Figure 19: Squared Hellinger distance for 2A - low-frequency observations. BPF: $N_P = 2.5 \times 10^6$.

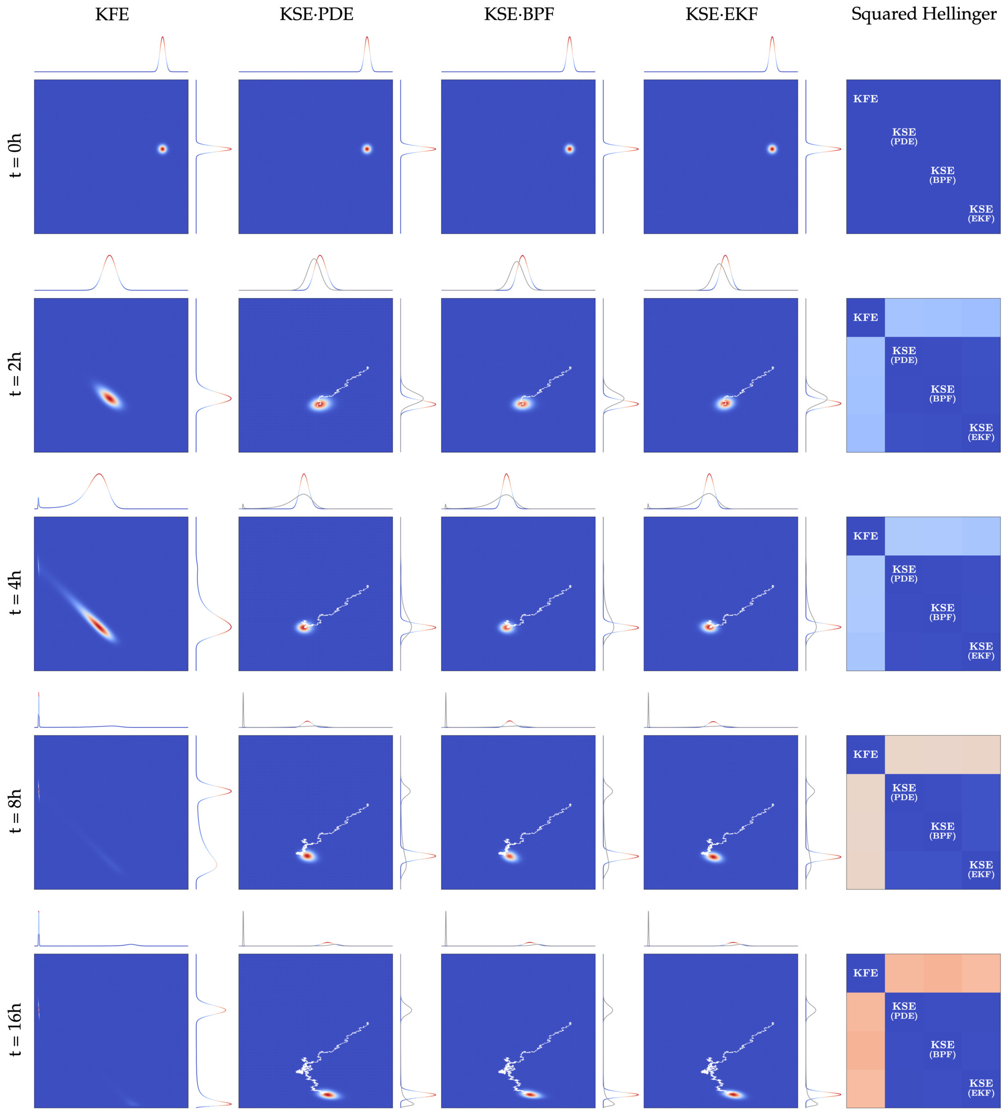

Figure 20: Approximation to the solution of the Kushner-Stratonovich equation for the growth function of the Haldane type with sparse observations of biogas flow rate. $512^2$ grid. $N_P = 2.5 \times 10^6$.

**2B - low-frequency observations (Haldane kinetics, sparse observations of substrate concentration)**

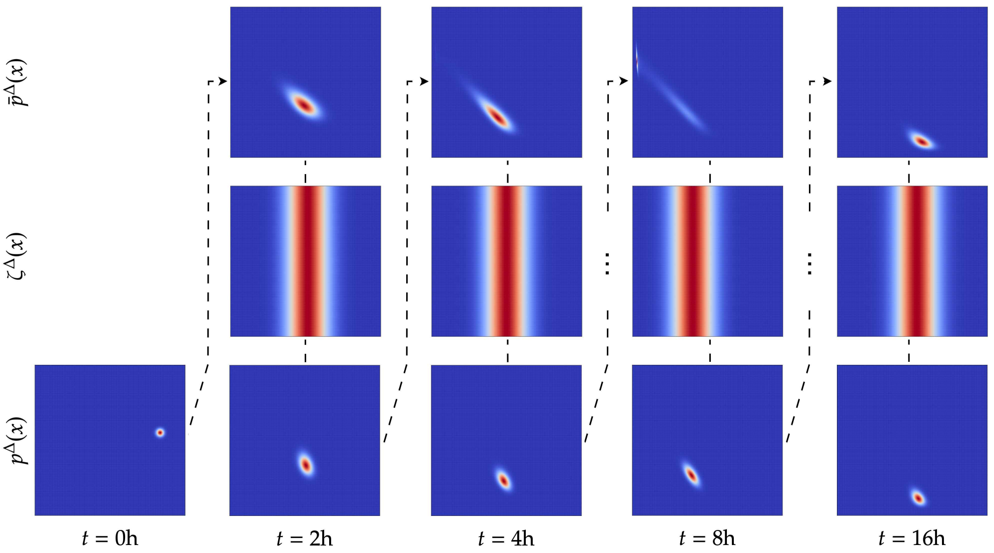

Figure 21: Splitting-up approximation for the Kushner-Stratonovich Equation with PDE using a $256^2$ grid.

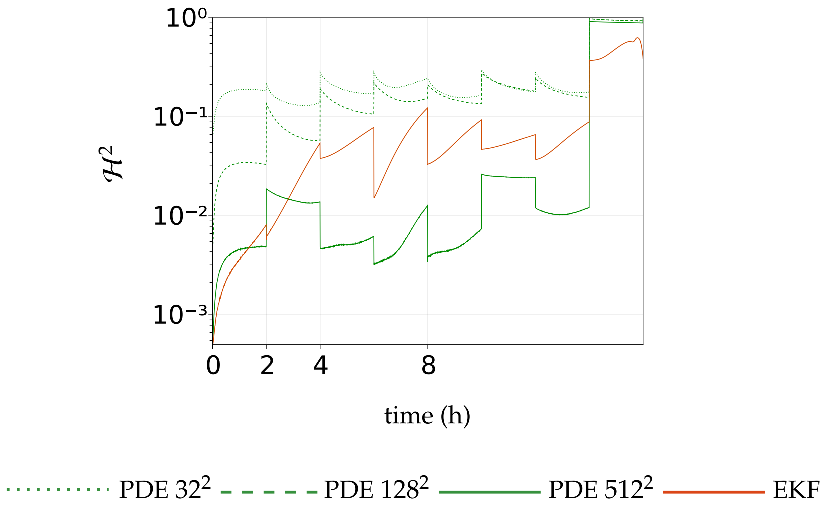

Figure 22: Squared Hellinger distance for 2B - low-frequency observations. BPF: $N_P = 2.5 \times 10^6$.

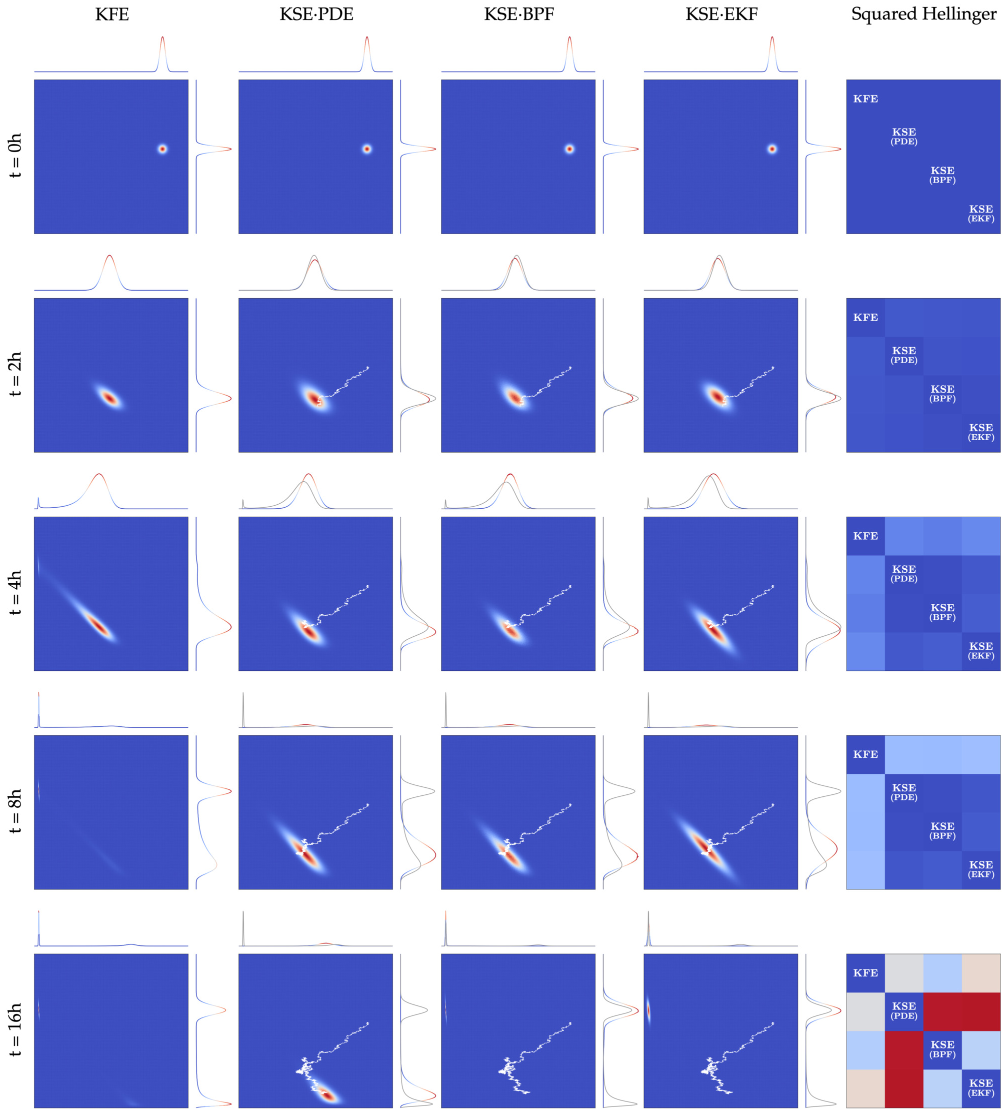

Figure 23: Approximation to the solution of the Kushner-Stratonovich equation for the growth function of the Haldane type with sparse observations of substrate concentration. $512^2$ grid. $N_P = 2.5 \times 10^6$.

The distribution of the particles $\{X_t^{n_P}\}_{n_P=1}^{N_P}$ given approximately by the evolution step is "far" from $\pi_{t_n}^{N_P}$ in the sense that the ratio (i.e., the Radon-Nykodym derivative) of these two distributions generates importance weights with a high variance. One of the reasons for why this happens comes from our sparse observations setup: the dynamics allow particles to disperse far away between observation times; if $N_P$ is not large enough, we will not be able to get a good approximation to Eq. (6) (the Kallianpur-Striebel formula).

To show that a larger number of particles can improve the approximation, we now compute the Hellinger distance against the BPF with $N_P = 3.5 \times 10^6$.

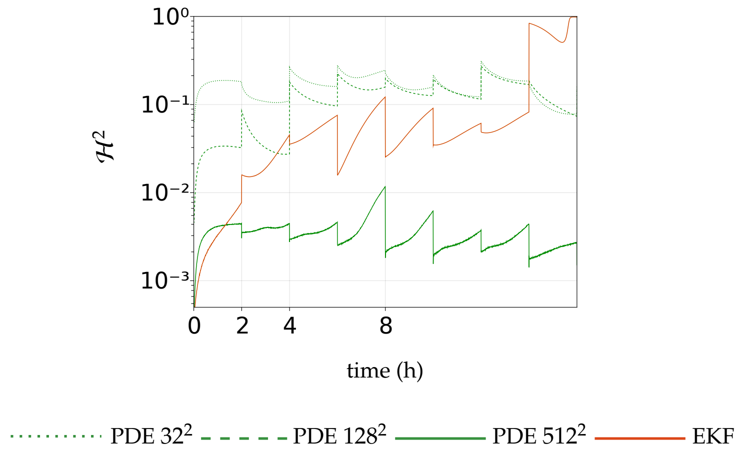

Figure 24: Squared Hellinger distance for 2B - low-frequency. BPF with $N_P = 3.5 \times 10^6$. It is clear that the approximation from the particle filter is improved. The resulting approximation is closer to the approximation with methods for PDE with a fine mesh refinement, and the mass of the distribution will be concentrated around the washout steady state as $t \to \infty$.

#### 2C - low-frequency observations (Haldane kinetics, sparse observations of biomass concentration)

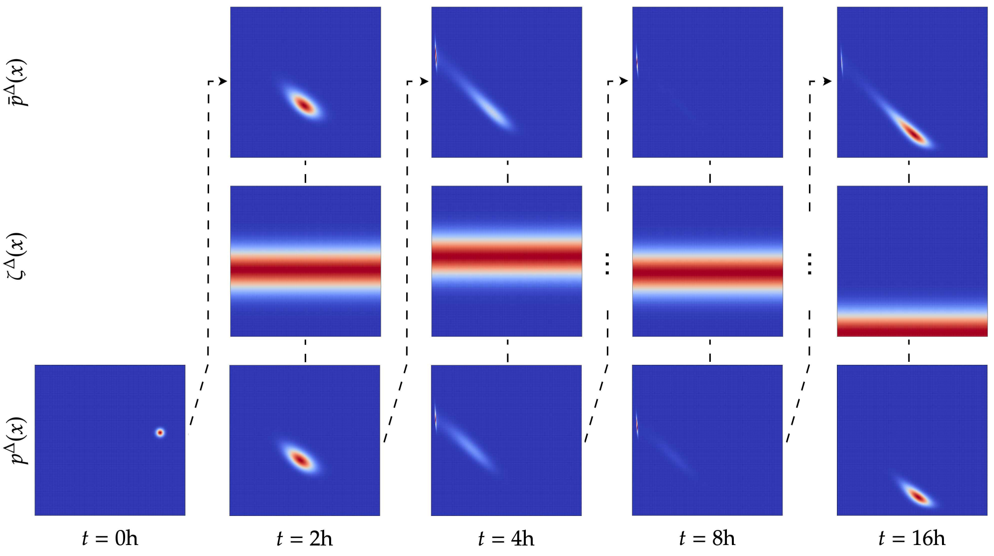

Figure 25: Splitting-up approximation for the KSE using a $256^2$ grid.

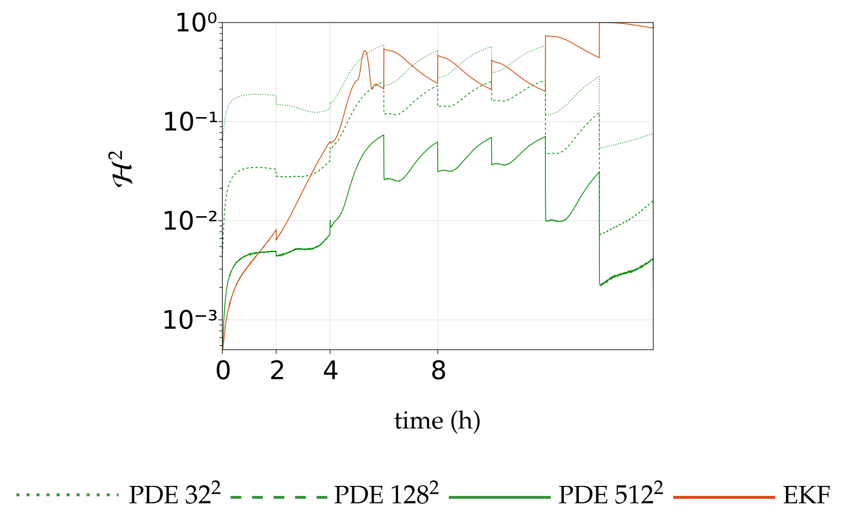

Figure 26: Squared Hellinger distance for 2C - low-frequency observations. BPF: $N_P = 2.5 \times 10^6$.

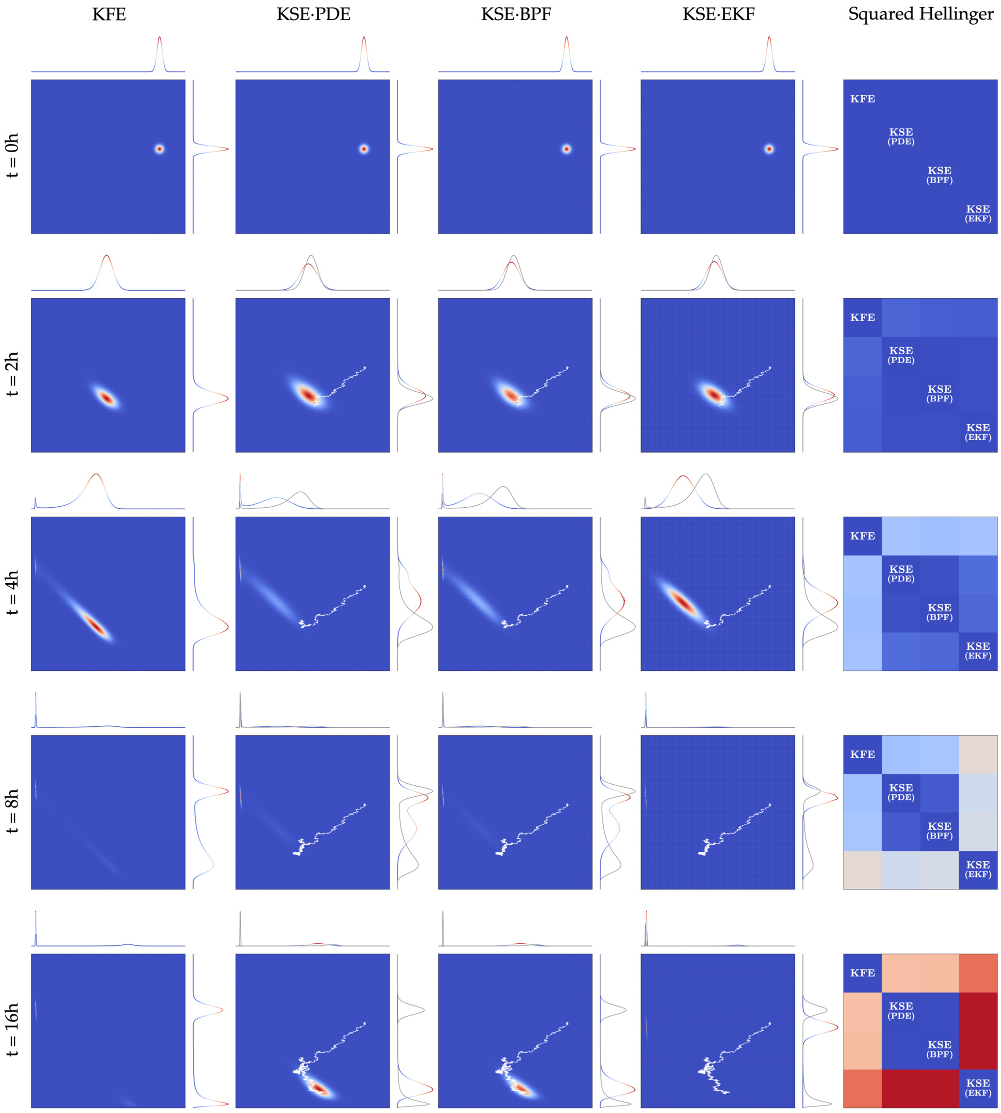

Figure 27: Approximation to the solution of the Kushner-Stratonovich equation for the growth function of the Haldane type with sparse observations of biomass concentration. $512^2$ grid. $N_P = 2.5 \times 10^6$.

**2D - low-frequency observations (Haldane kinetics, sparse observations of biomass and substrate concentrations)**

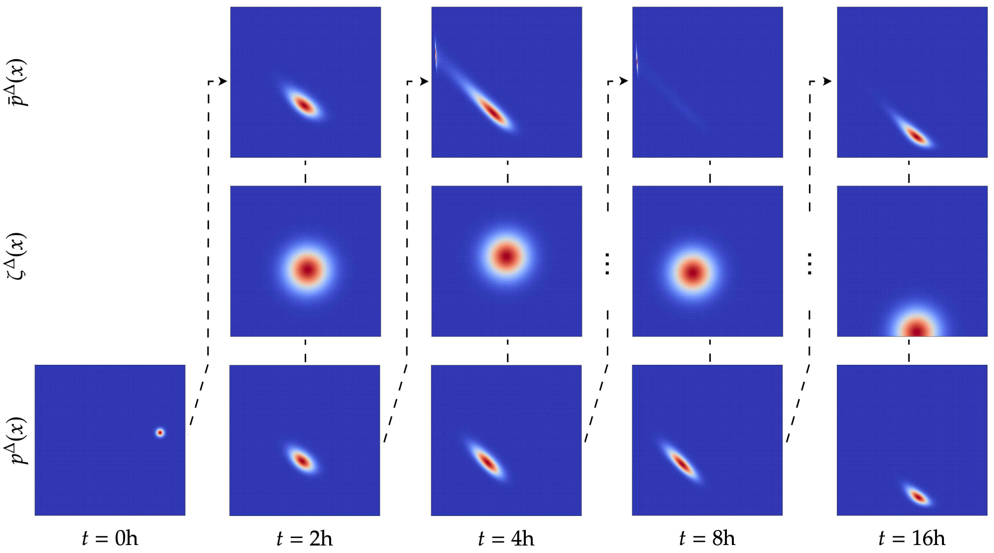

Figure 28: Splitting-up approximation for the KSE using a $256^2$ grid.

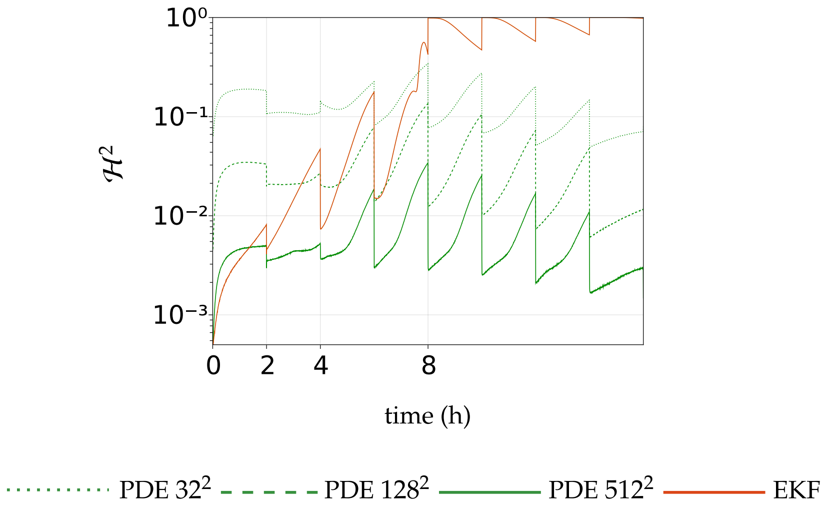

Figure 29: Squared Hellinger distance for 2D - low-frequency observations. BPF: $N_P = 2.5 \times 10^6$.

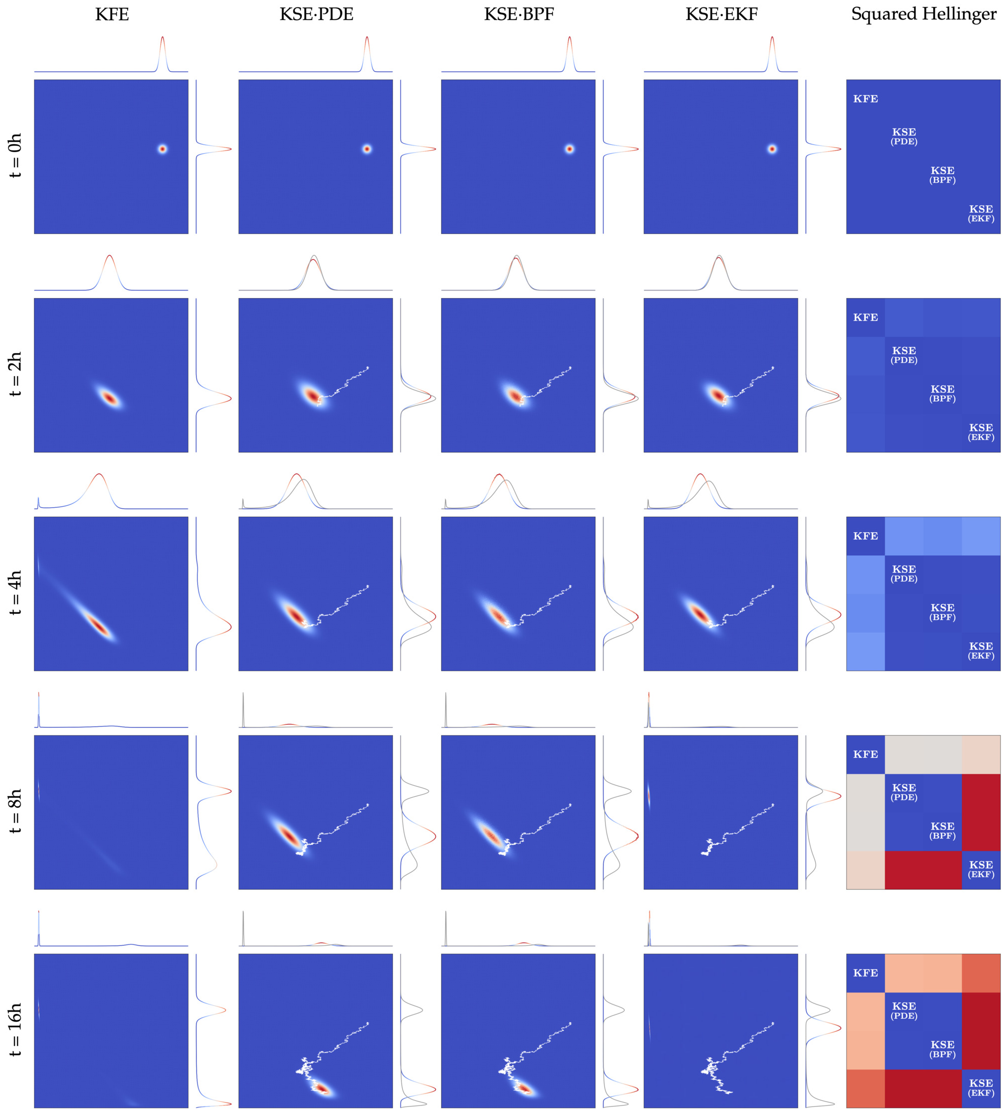

Figure 30: Approximation to the solution of the Kushner-Stratonovich equation for the growth function of the Haldane type with sparse observations of biomass and substrate concentrations. $512^2$ grid. $N_P = 2.5 \times 10^6$.
